# Selecting a Window Size for the Analysis of Whole Genome Alignments using AIC

**DOI:** 10.1101/2025.02.05.636755

**Authors:** Jeremias Ivan, Paul Frandsen, Rob Lanfear

## Abstract

The variation of evolutionary histories along the genome presents a challenge for phylogenomic methods to identify the non-recombining regions and reconstruct the phylogenetic tree for each region. To address this problem, many studies used the non-overlapping window approach, often with an arbitrary selection of fixed window sizes that potentially include intra-window recombination events. In this study, we proposed an information theoretic approach to select a window size that best reflects the underlying histories of the alignment. First, we simulated chromosome alignments that reflected the key characteristics of an empirical dataset and found that the AIC is a good predictor of window size accuracy in correctly recovering the tree topologies of the alignment. Due to the issue of missing data in empirical datasets, we then designed a stepwise non-overlapping window approach and applied this method to the genomes of *erato*-*sara Heliconius* butterflies and great apes. We found that the best window sizes for the butterflies’ chromosomes ranged from ≤125bp to 250bp, which are much shorter than those used in a previous study even though this difference in window size did not significantly change the most common topologies across the genome. On the other hand, the best window sizes for great apes’ chromosomes ranged from 500bp to 1kb with the proportion of the major topology (grouping human and chimpanzee) falling between 60% and 87%, consistent with previous findings. Additionally, we observed a notable impact of stochastic error and concatenation when using small and large windows, respectively. For instance, the proportion of the major topology for great apes was 50% when using 250bp windows, but reached almost 100% for 64kb windows. In conclusion, our study highlights the challenges associated with selecting a window size in non-overlapping window analyses and proposes the AIC as a more objective way to select the optimal window size for whole genome alignments.

## Introduction

In recent years, the variation of evolutionary trees across different loci or genomic regions, also known as gene tree discordance, has become an interest for many genome-wide phylogenetic studies (Pease et al. 2016; Copetti et al. 2017; Morales-Briones et al. 2021; Lescroart et al. 2023; McLay et al. 2023; Herrig et al. 2024). As this incongruence might reveal important evolutionary histories, such as incomplete lineage sorting (ILS) and hybridisation (Maddison 1997; Degnan and Rosenberg 2009; Mallet et al. 2016; Hibbins et al. 2023), efforts have been made to not only infer the correct tree for each locus but also to estimate the locations of the recombination events that define the loci themselves. Population genetic-based methods such as Ancestral Recombination Graphs (e.g., ARGweaver (Rasmussen et al. 2014), tsinfer (Kelleher et al. 2019), and Relate (Speidel et al. 2019)) represent a powerful approach to estimate the evolutionary history of every site in the genome. However, these methods have considerable challenges, including scalability and the estimation of population genetic statistics such as recombination rate. At deeper timescales, Hidden Markov Models (HMMs) can be used to co-infer the history of every site in the genome as well as the recombination breakpoints in the presence of extensive ILS, recombination, and hybridisation, with an assumption that neighbouring sites are likely to share evolutionary histories. However, the current implementations of HMMs do not scale to whole genome sequence analysis (e.g., PhyML_Multi (Boussau et al. 2009; Brandt et al. 2024) and PhyloNet-HMM (Liu et al. 2014)) or limited to rooted triplets at a time (e.g., Coal-HMM (Hobolth et al. 2007; Dutheil et al. 2009)). Recently, Wong et al. <u>(2024)</u> proposed the MAST model (Mixture Across Sites and Trees) that uses a mixture of bifurcating trees to infer the weight of each topology (i.e., tree without branch lengths) within the genomic sequences. This model determines the likelihood that each site in the genome follows a particular history, and can scale to large datasets. However, MAST requires a set of input topologies that needs to be derived from other methods. The upshot is that inferring the history of every site in many genomic datasets remains challenging.

One of the most common approaches to the problem of inferring locus tree topologies from non-recombining regions along a set of aligned genomes is a relatively simple method known as the non-overlapping window approach. In this approach, the alignment is first partitioned into non-overlapping blocks of equal size called ‘windows’. Then, a phylogenetic tree is reconstructed for every window with an assumption that each window has no intra-window recombination. This approach has been applied by many studies to explore the variation of phylogenetic histories across genomes <u>(e.g., Fontaine et al. 2015; Pease et al. 2016; Edelman et al. 2019; Meleshko et al. 2021; Yang et al. 2021; Feng et al. 2023; Lescroart et al. 2023; Herrig et al. 2024)</u>. However, one of the main drawbacks of this approach is that there is no objective method to determine the ideal window size (i.e., the window size that most closely reflects the recombination patterns of the alignment). For example, Yang et al. <u>(2021)</u> assessed the genome-wide topology distribution of Mediterranean lizards using six different window sizes, which ranged from 5kb to 200kb. On the other hand, Herrig et al. <u>(2024)</u> used longer windows of 10kb, 50kb, 100kb, 500kb, and 1Mb to generate the species phylogeny of *Neodiprion* sawflies. Despite this wide array of choices of window size, there remains no objective justification for the choice of any particular size.

This may not be an issue if the selected window size is sufficiently short that most windows do not contain recombination breakpoints. However, the empirical evidence suggests that this is often not the case. For example, Edelman et al. <u>(2019)</u> found that in the overwhelming majority of cases, the tree topology changed from one window to the next. This implies that many windows were likely to contain recombination breakpoints, because if the windows contained no breakpoints, then many consecutive windows would be expected to have identical tree topologies resulting from the splitting of a non-recombining region into many windows. Similarly, there is growing evidence that neighbouring exons in genes tend to have different evolutionary histories (Scornavacca and Galtier 2017; Mendes et al. 2019), and that non-recombining regions (sometimes called ‘coalescent-genes’ or ‘c-genes’) might be surprisingly small even in datasets with modest recombination rates (e.g., ∼12bp in mammals, although this remains an active area of discussion (Edwards et al. 2016; Springer and Gatesy 2016, 2018)). Because non-overlapping windows are widely used and the optimal window size is difficult to determine *a priori*, an objective approach to select the window size would provide a good starting point to generate a set of window alignments and their associated single-locus trees. Estimates of such trees can further be refined by other methods such as Espalier (Rasmussen and Guo 2023), MAST (Wong et al. 2024), and BUCKy (Larget et al. 2010), re-estimated from the window alignments with multispecies coalescent methods such as StarBEAST (Heled and Drummond 2010; Ogilvie et al. 2017; Douglas et al. 2022) and BPP (Flouri et al. 2018), or used directly as input for estimating species trees or networks with methods such as ASTRAL (Zhang et al. 2018b), PhyloNet (Wen et al. 2018), and SpeciesNetwork (Zhang et al. 2018a).

In this study, we sought to develop an objective window size selection method using information theoretic criteria commonly used in phylogenetics, such as the AIC (Akaike Information Criterion (Akaike 1974)) and BIC (Bayesian Information Criterion (Schwarz 1978)). These measures may help to choose the optimal window size because they seek to find the best balance between estimating too many parameters (i.e., having a window size that is too small) and too few (i.e., having a window size that is too large). For example, the AIC has been used to choose the optimal window size in copy-number alteration analysis on low-coverage tumour sample data (Gusnanto et al. 2014). We acknowledge that a single window size is insufficient to represent an entire genome because recombination rates are well-known to vary along chromosomes <u>(e.g.</u>, <u>Chan et al., 2012; Jensen-Seaman et al., 2004; McVean et al., 2004)</u>. Nevertheless, an objective method to determine a single best-fit window size represents a substantial improvement over the status quo as the window partitions would more closely reflect the recombination patterns of the genomes, allowing us to recover the ‘true’ history of each genomic region. In order to achieve this, we first used simulated data to develop an information theoretic approach that allows users to choose a window size that maximises the number of sites in the genome that correctly recover their true topology. We then applied this approach to select the optimal window sizes for empirical datasets of *Heliconius* butterflies and great apes, and used these window sizes to analyse the genome-wide evolutionary histories of these two groups.

## Materials & Methods

### Overview

A non-overlapping window analysis proceeds by choosing a window size, splitting the genome into non-overlapping windows of that size, and estimating a tree for each window, which in this study was done using Maximum Likelihood (ML) method. In order to ask whether information theoretic criteria (i.e., the AIC and/or BIC) are appropriate to help finding the optimal window size, we simulated a wide range of datasets that mimic a recent empirical study by Edelman et al. <u>(2019</u>) and used these to assess the performance of the AIC and BIC as criteria for choosing the best non-overlapping window size (i.e., the window size with the highest accuracy) when the truth is known. We measured the accuracy of each window size in one of two ways: the proportion of sites from the non-overlapping window analysis assigned to the true (simulated) topology, which we call site accuracy (i.e., a site accuracy of 100% indicates that every site is assigned the topology from which it is simulated), and the root mean squared error (RMSE) of the estimated distribution of tree topologies from the non-overlapping window analysis when compared to the true (simulated) distribution. We assessed the performance of the AIC and BIC by asking whether either correlates with these measures of accuracy. We then applied the best-performing method to two empirical datasets: one from *Heliconius* butterflies (Edelman et al., 2019) and another from great apes (i.e., humans, chimpanzees, gorillas, and orangutans <u>(Waterson et al. 2005; Locke et al. 2011; Scally et al. 2012; Schneider et al. 2017)</u>). All the codes necessary to reproduce the methods in this paper is available at https://github.com/jeremiasivan/SimNOW.

### Simulating Alignments Based on *Erato-Sara Heliconius* Butterflies’ Chromosome

In order to create a realistic simulation scenario, we focused on a well-studied empirical system, the *Heliconius* butterflies. Specifically, we sought to simulate data based on the seven *Heliconius* species that were the primary focus of the work in Edelman et al. <u>(2019</u>): six ingroup species from *erato-sara* clade (*H. erato*, *H. himera*, *H. hecalesia*, *H. telesiphe*, *H. demeter*, and *H. sara*) and one outgroup species (*H. melpomene*). Throughout, we took care to choose simulation parameters such that the resulting chromosome alignments closely matched the key characteristics of the empirical alignments from Edelman et al. (2019), including the proportion of informative sites and distribution of tree topologies among loci. Full details are provided in the Supplementary Information.

Our simulation design was inspired by <u>Forsythe et al. (2020)</u>, except that we used AliSim (Ly-Trong et al. 2022) instead of Seq-Gen (Rambaut and Grass 1997) to simulate the chromosome alignments as AliSim has better runtime and memory usage compared to Seq-Gen and other simulation software (Ly-Trong et al. 2022). We first simulated the locus trees using ms (Hudson 2002), along a species tree with ILS and introgression. To do this, we used the species tree topology from Figure 2B of Edelman et al. <u>(2019</u>) and incorporated the three bi-directional introgression events with introgression probabilities estimated from chr11 in Thawornwattana et al. <u>(2022)</u>: *H. himera ⇄ H. erato; H. telesiphe → H. hecalesia;* and *H. telesiphe* ⇄ (*H. demeter*, *H. sara*) (Fig. S1). We selected chromosome 11 (chr11) as the basis of our simulations because it has an intermediate length in the *erato-sara Heliconius*’ genomes. We used speciation dates from Kozak et al. <u>(2015)</u> rather than from Thawornwattana et al. <u>(2022)</u> because the former included an estimate for the outgroup divergence time. Then, we dated each introgression event on the relevant branch proportionally to the estimated time reported in Thawornwattana et al. <u>(2022)</u>.

As ms requires the speciation and introgression timings to be in coalescent units, we converted the branch lengths from Myr to coalescent units using estimates of population genetic statistics from *Heliconius* butterflies to provide sensible bounds, and within these bounds selected the parameters that best mimicked the key characteristics of the empirical dataset (Table S1-S2, see Supplementary Information for details). This resulted in using a conversion factor of 4N_e_ generations being equivalent to 0.75Myr. We set the total alignment length to 10Mb and varied the recombination rate between 0 (i.e., no recombination) to 2000 (∼126,000 loci with an average length of 80bp, Table S3) using four settings: 0, 20, 200, and 2000. In addition, we examined three degrees of ILS (low, medium, and high) by varying the branch length scaling parameters (see Supplementary Information for details). We simulated 10 replicates per scenario, resulting in 120 sets of simulated chromosomal loci in ms (3 ILS levels * 4 recombination rates * 10 replicates). Command lines for each step are provided in the Supplementary Information. Then, for each set of the simulated chromosomal loci, we used the locus trees and the locus lengths as inputs for AliSim (Ly-Trong et al. 2022) to simulate each locus under the JC model, and concatenated them to reconstruct a single 10Mb chromosome alignment containing all loci.

### Assessing the AIC, BIC, and Accuracy of Non-Overlapping Window Analyses

For each simulated 10Mb chromosome alignment, we divided the whole alignment into non-overlapping windows with 16 different window sizes, ranging from 100bp to 10Mb. At one extreme, a 100bp window would rarely contain enough phylogenetic signal to build a reliable tree, while at the other extreme, a 10Mb window incorporated the entire simulated chromosomes and would violate the assumption that there is only one tree for each window (except for simulations with zero recombination). For each window size, we estimated the window trees with JC model using IQ-TREE2 (Minh et al. 2020) with the following command: `iqtree2 -S alndir -m JC -blmin 1/window_size -cptime 1000000`, where `alndir` refers to the directory storing all window alignments. The `-cptime` flag was set to reduce I/O and speed up the analysis. The `-blmin` flag sets the minimum branch length for all trees to represent at least one substitution per branch, in order to penalise stochastic error in tree topologies from windows with very little phylogenetic information.

We then extracted a single AIC and BIC score for each window size on each simulated alignment from the IQ-TREE2 output file which calculates the scores based on the log-likelihoods, number of free parameters, and number of sites (for the BIC) across all windows following the standard formulas (Akaike 1974; Schwarz 1978; Stone 1979). We also calculated the accuracy of each of the 16 different window sizes for each simulated alignment by first unrooting the trees from ms using ape v5.6 (Paradis et al. 2004) and then calculated the site accuracy and RMSE for each window size using R v4.2.3 (R Core Team 2023) following these formulas:

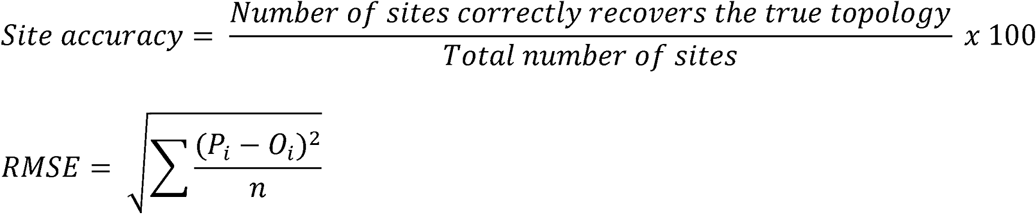

For site accuracy, the numerator counts the number of sites from the non-overlapping window analysis that recovers the true (simulated) topology from ms, while the denominator reflects the alignment length (in our case, 10Mb). For RMSE, *P_i_* and *O_i_* respectively denote the predicted weight (from ms simulation) and observed weight (from non-overlapping windows analysis) for the *i^th^* topology, while *n* denotes the number of unique topologies.

### Testing for the Accuracy of the AIC and BIC as Model Selection Criteria for Non-Overlapping Window Analyses

We compared the AIC and BIC scores to our two measures of accuracy (i.e., site accuracy and RMSE) across all window sizes for each simulation. To reflect the way that the AIC and BIC are used, we did this by calculating the loss of accuracy that would be incurred when using each metric to select the best window size (i.e., we compared the accuracy of the most accurate window size that gives the best site accuracy and lowest RMSE to the accuracy of the window size that would be selected using the AIC or BIC).

### Selecting Non-Overlapping Window Sizes on Empirical Datasets

Our simulation results showed that the AIC, but not the BIC, was a useful metric for selecting non-overlapping window sizes (see Results). Next, we sought to use the AIC to select the optimal window sizes for empirical datasets of *erato*-*sara Heliconius* butterflies and great apes. Initially, we applied the method we described above (i.e., calculating the AIC score for a wide range of non-overlapping window sizes and choosing the window size with the best score). However, this approach has a considerable limitation with empirical datasets: the shortest window size that can be analysed is that for which it is possible to calculate a likelihood for *every* window in the alignment, because comparing AIC scores (in this case between window sizes) assumes that the underlying alignment is identical. As empirical datasets tend to contain many regions where fewer than three species contain data, this can limit the shortest window size that can be analysed, precluding the calculation of a likelihood in IQ-TREE2, to quite large windows (e.g., many kilobases in the dataset we used).

To overcome this limitation, we designed a stepwise non-overlapping window approach to apply our methods to empirical datasets. In this approach, we start with the longest window size and make a series of pairwise comparisons where the window size is halved each time. Within each pairwise comparison, if a particular window cannot be analysed due to missing data, we remove that window from the alignment for both window sizes, such that the resulting alignment consists of windows that *can* be analysed for both members of the pair. This approach allows us to test much smaller window sizes but at the cost of using less of the alignment as the window size becomes smaller. This method is available in the GitHub repository under https://github.com/jeremiasivan/SimNOW/tree/main/empirical_analysis/stepwise_now/.

### Erato-sara Heliconius Butterflies

We retrieved the 21 chromosome alignments from the original dataset (Edelman et al. 2019) following the same steps when we retrieved and measured the key characteristics of chr11 for the simulations (command lines are provided in the Supplementary Information). For each chromosome, we ran stepwise non-overlapping windows with an initial window size of 64kb, then made a series of pairwise comparisons by comparing the starting window size (e.g., 64kb) to a window size of half that length (e.g., 32kb). We generated individual window trees for both window sizes using IQ-TREE2 (Minh et al. 2020) with the following command: `iqtree2 -s path_to_fasta -blmin 1/window_size -bb 1000`, where `-s` refers to the alignment file while `-bb` sets the number of UFBoot replicates. Then, we calculated the AIC for each window size by summing up the log-likelihoods and number of free parameters across windows following the original formula (Akaike 1974). When the number of unique sequences was less than four, we ran IQ-TREE2 without bootstrapping to save computational time. We continued to halve the window size (e.g., 32kb and 16kb, 16kb and 8kb, and so on) until it reached 125bp, which cannot be divided by 2. Lastly, we summarised the topology distributions from the best window sizes (i.e., window size with the best AIC score for each chromosome) and compared our results with those from Edelman et al. <u>(2019)</u>.

### Great Apes

We also ran the stepwise non-overlapping window approach on genome alignments of great apes, consisting of four species: *Homo sapiens* (hg38; human), *Pan troglodytes* (panTro4; chimpanzee), *Gorilla gorilla gorilla* (gorGor3; gorilla), and *Pongo pygmaeus abelii* (ponAbe2; orangutan). First, we downloaded the compressed MAF file for each chromosome from the UCSC Genome Browser (https://hgdownload.soe.ucsc.edu/goldenPath/hg38/multiz20way), resulting in 25 chromosome (22 autosomal, X, Y, and mitochondrial) alignments. Then, we uncompressed the files and converted them to FASTA using PHAST v1.5 (Hubisz et al. 2011) with the following commands:

~~~
$ msa_view {i}.maf -i MAF -m –G 1 > {i}.fa
$ msa_view -l hg38,gorGor3,panTro4,ponAbe2 {i}.fa >
greatapes_{i}.fa
~~~

where `i` represents the chromosome number (or name). We ran the stepwise non-overlapping window analysis on each chromosome following the same steps and parameters from the *Heliconius* analyses detailed above.

## Results

### Simulated Datasets

Summary statistics of 120 simulated datasets are shown in Table S3. We found that the degree of ILS had no meaningful effect on the results (Fig. 1-2, S2-S20), so we focus on the results from the medium ILS simulations here (which most closely mimic the empirical datasets from Edelman et al. (2019)) and present the rest in the Supplementary Information. Figure 1 shows that, as expected, the best window size (i.e., window size with the highest site accuracy and lowest RMSE) becomes shorter as the recombination rate increases due to more frequent topology switching and consequently shorter true locus lengths. For instance, the best window sizes for ρ = 0, 20, 200, and 2000 are ≥50kb, 20kb, 5kb, and 2kb respectively (Fig. 1).

**Figure 1.**
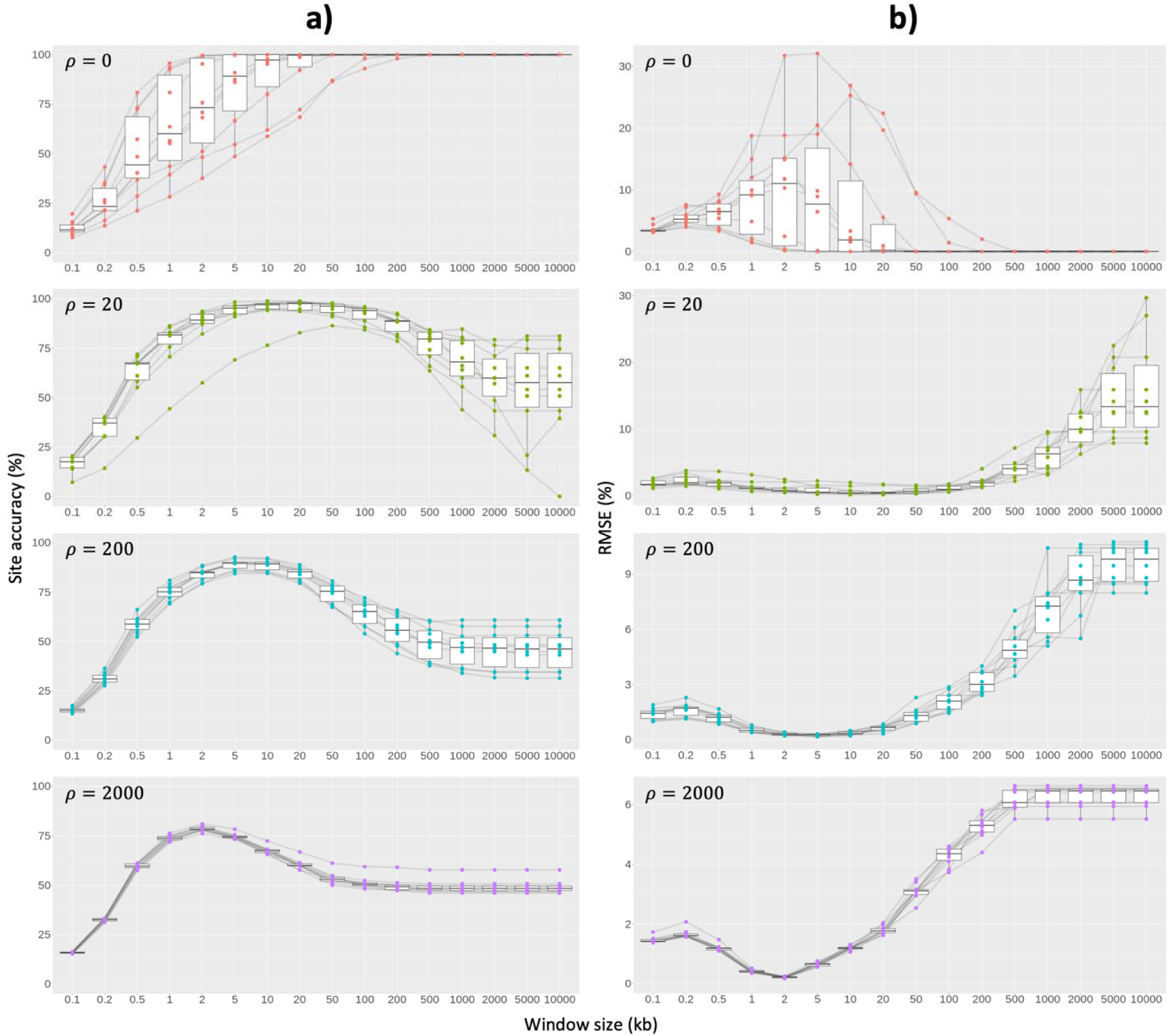
(a) Site accuracy and (b) RMSE of non-overlapping windows on simulated chromosomes with medium ILS level. Each dot represents an individual result. Black lines connect results from the same simulated alignment.

To assess the potential utility of the AIC and BIC, we compared both with the site accuracy and RMSE measured from the simulated chromosomes. The AIC frequently picks the most accurate window size according to both criteria, but the BIC does not (Fig. 2, Table S3). The BIC performs poorly in predicting accuracy (Fig. 2b, S9-S13), leading to up to 18.5% site accuracy loss and 1.5% RMSE increase (when ρ = 2000) compared to the best window size if we always choose window size with the lowest BIC (Fig. S13b, d). On the other hand, the AIC is a good predictor of both site accuracy and RMSE (Fig. 2a, S9-S13), losing up to 0.25% site accuracy with ∼0.1% RMSE increase (when ρ = 20) if we always choose window size with the lowest AIC (Fig. S13a, c). These results are consistent across datasets with different degrees of ILS (Fig. S2-S8, S14-S20), and thus we propose the AIC can be used to select the best window size on empirical alignments.

**Figure 2.**
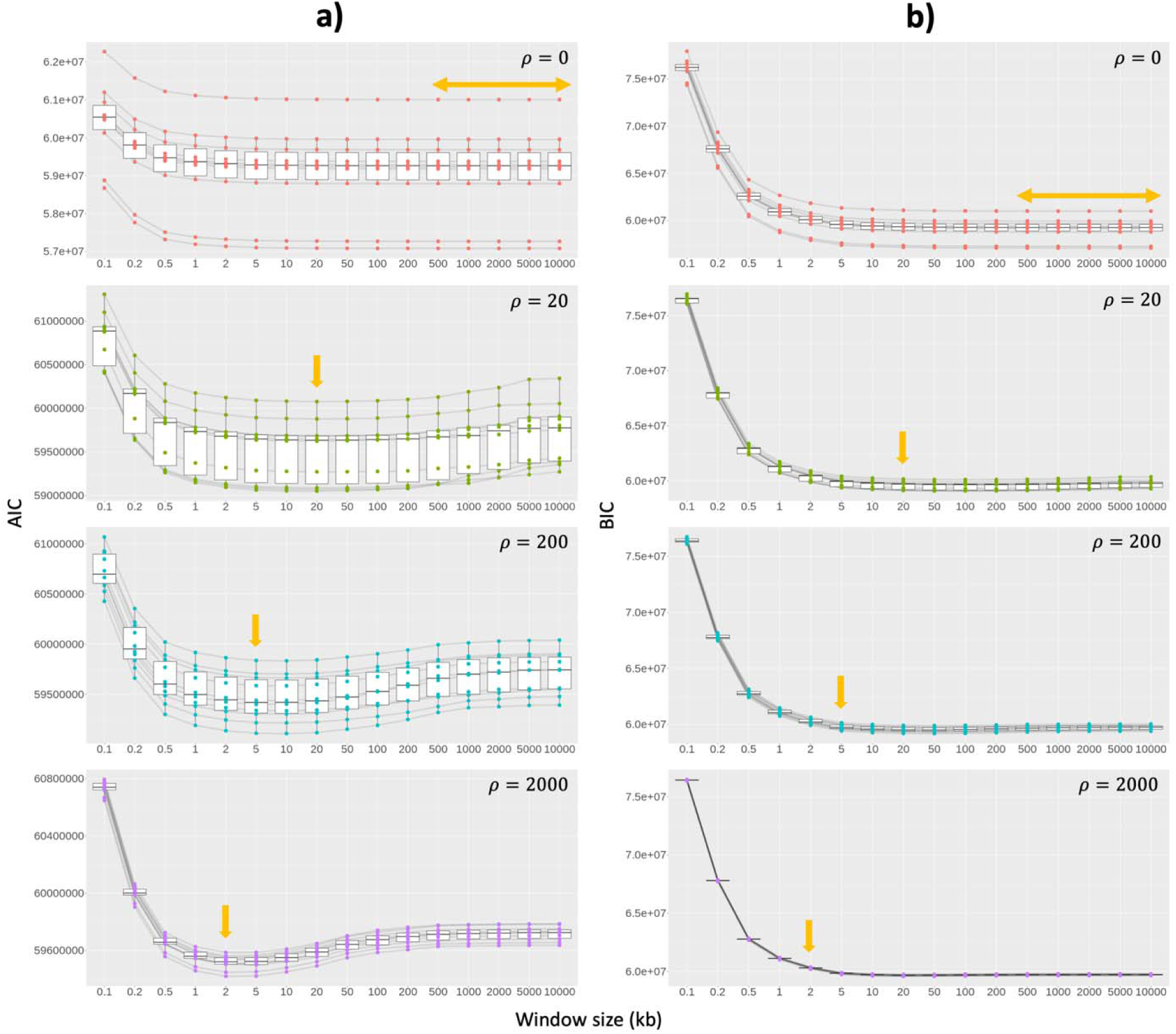
Correlation between the (a) AIC and (b) BIC with window sizes on simulated chromosomes with medium ILS level. Light orange arrows show window size(s) with the highest average site accuracy from Figure 1. Each box represents different recombination rates. Each dot represents an individual result. Black lines connect results from the same simulated alignment.

### Empirical Datasets

In order to address the issue of missing data in empirical alignments, we designed a stepwise non-overlapping window approach that compares two window sizes at a time and includes only windows that can be analysed using both window sizes. This approach enabled us to use shorter windows while ensuring the alignment consistency between the two window sizes, which is crucial when comparing the AIC scores. All results for the empirical datasets were derived from the stepwise non-overlapping window method rather than the common non-overlapping windows approach.

### Erato-sara Heliconius Butterflies

Using the stepwise non-overlapping window approach, we found that the sex chromosome and the ten longest autosomal chromosomes in *erato-sara Heliconius*’ genomes have a best window size of 250bp, while the AIC of other chromosomes was still declining after reaching 125bp with much smaller delta AIC scores than the previous step (500bp vs. 250bp) (Table S4-S5), suggesting that the best window size for these chromosomes is at most 125bp. Moreover, we also found that the selection of window sizes affects the distribution of the tree topologies recovered from the group (Table S6-S7). For example, the topology distribution of chr11 only includes <10 unique topologies and is dominated by Tree 1 using 64kb windows, but becomes much more variable as the window sizes become smaller (Fig. S21).

Figure 3 shows the most common tree topologies recovered from the sliding window (SW, first column) and non-overlapping window analyses (NOW, second column) in Edelman et al. <u>(2019)</u> as well as the stepwise non-overlapping windows (stepwise NOW, third and fourth columns) from our analyses, where the best window size was selected for each chromosome using the AIC (Table S4). The stepwise non-overlapping window approach recovers a much broader distribution of tree topologies, with the most frequently recovered 5 topologies making up only 19.1% of all window trees, most likely due to stochastic errors introduced by very short window sizes (Fig. 3, third column). This is also reflected by the number of consecutive windows that recover the same topology which is mostly one (Fig. S22). When considering only the well-supported tree topologies (i.e., trees with ≥95 average UFBoot support), the most frequent ten topologies account for 85.9% of all well-supported tree topologies (Fig. 3, fourth column). Using this approach, Tree 3 is the most common topology, closely followed by Trees 1 and 2 from the original study (in which Tree 2 is considered to be the species tree). This analysis also finds two additional topologies, Tree 9 and Tree 10, that respectively represent 8.7% and 2.1% of the well-supported trees.

**Figure 3.**
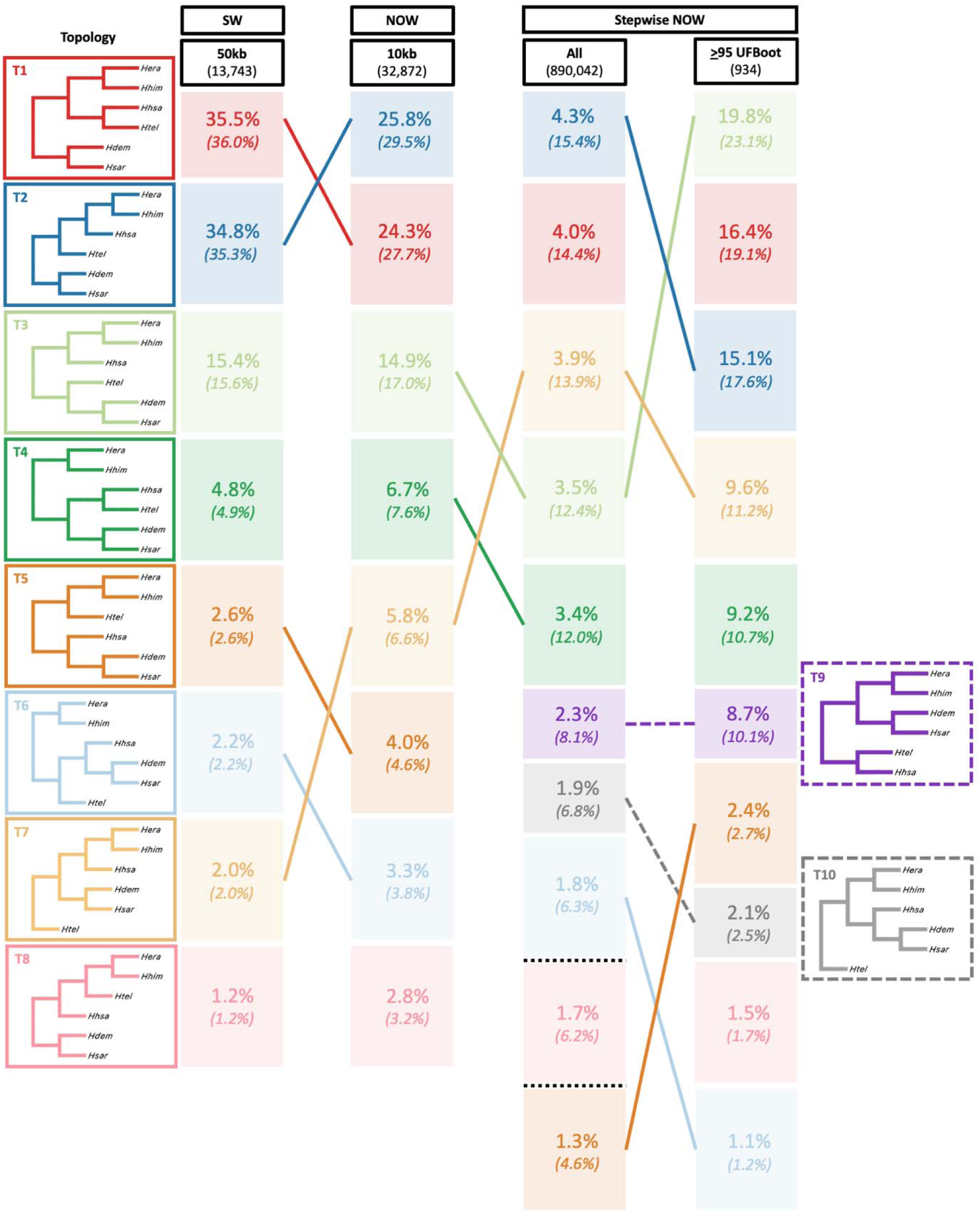
Comparison of topology frequency of the most common tree topologies from *erato-sara Heliconius*’ genomes between 50kb sliding windows (SW) and 10kb non-overlapping windows (NOW) from Edelman et al. <u>(2019</u>), as well as stepwise NOW using the best window size for each chromosome (Table S4). The number under each method reflects the total number of windows. Dashed lines refer to additional topologies that are not present in the original eight, while black dotted lines refer to other topologies not listed. Numbers in brackets show the normalised value for each distribution, rounded to one decimal place so the total might not be exactly 100%. Topologies are visualised using FigTree v1.4.4 (http://tree.bio.ed.ac.uk/software/figtree) without branch lengths.

### Great Apes

Table S8 shows the results of the stepwise non-overlapping window analyses for the great apes. The best window size for most chromosomes is either 500bp or 1kb, except for the mitochondrial DNA (best window size of 4kb and all windows recover the same topology; Fig. S23), and the Y chromosome (best window size of 500bp, but noting that this alignment included just 2.24% of the initial alignment, reflecting the presence of all-gapped taxa in most of the windows).

The distribution of tree topologies from the stepwise non-overlapping windows follows the expected pattern, in which the (major) topology that groups human and chimpanzee is the most common, followed by a roughly equal proportion of the other two (minor) topologies. For example, if we count all window trees based on the best window size for each chromosome in Table S8, the ratio between the major topology and the two minor topologies is roughly 60.7: 19.8 : 19.5 (Table S9 (left), Fig. 4 (first column)). If we consider only highly-supported window trees with ≥95 average UFBoot support, the proportions change to 86.8 : 6.9 : 6.3 (Table S9 (right), Fig. 4 (second column)). However, both of these approaches may not reflect the underlying distribution of tree topologies, as including all trees might introduce stochastic error while highly-supported trees would bias towards the proportion of major topology (see Discussion). Moreover, Figure S24 and Figure 4 (rightmost panel) show that most non-recombining blocks (i.e., number of consecutive windows that recover the same topology) are 20-window long or less by considering all trees. Separating the trees by at least 50 windows, which is likely to be sufficient to ensure that each tree is independent of each other, shows that the proportions approach an asymptote of approximately 60 : 20 : 20 when considering all trees and 82 : 9 : 9 for highly supported ones (Fig. 4, S25 (third and fourth columns)). This might suggest that the effect of double (or multiple) counting of one non-recombing block is less apparent when considering all window trees (proportion of major topology changed from 60.7% to 59.7%, Fig. 4) compared to the highly-supported trees (proportion of major topology changed from 86.8% to 81.8%, Fig. 4).

**Figure 4.**
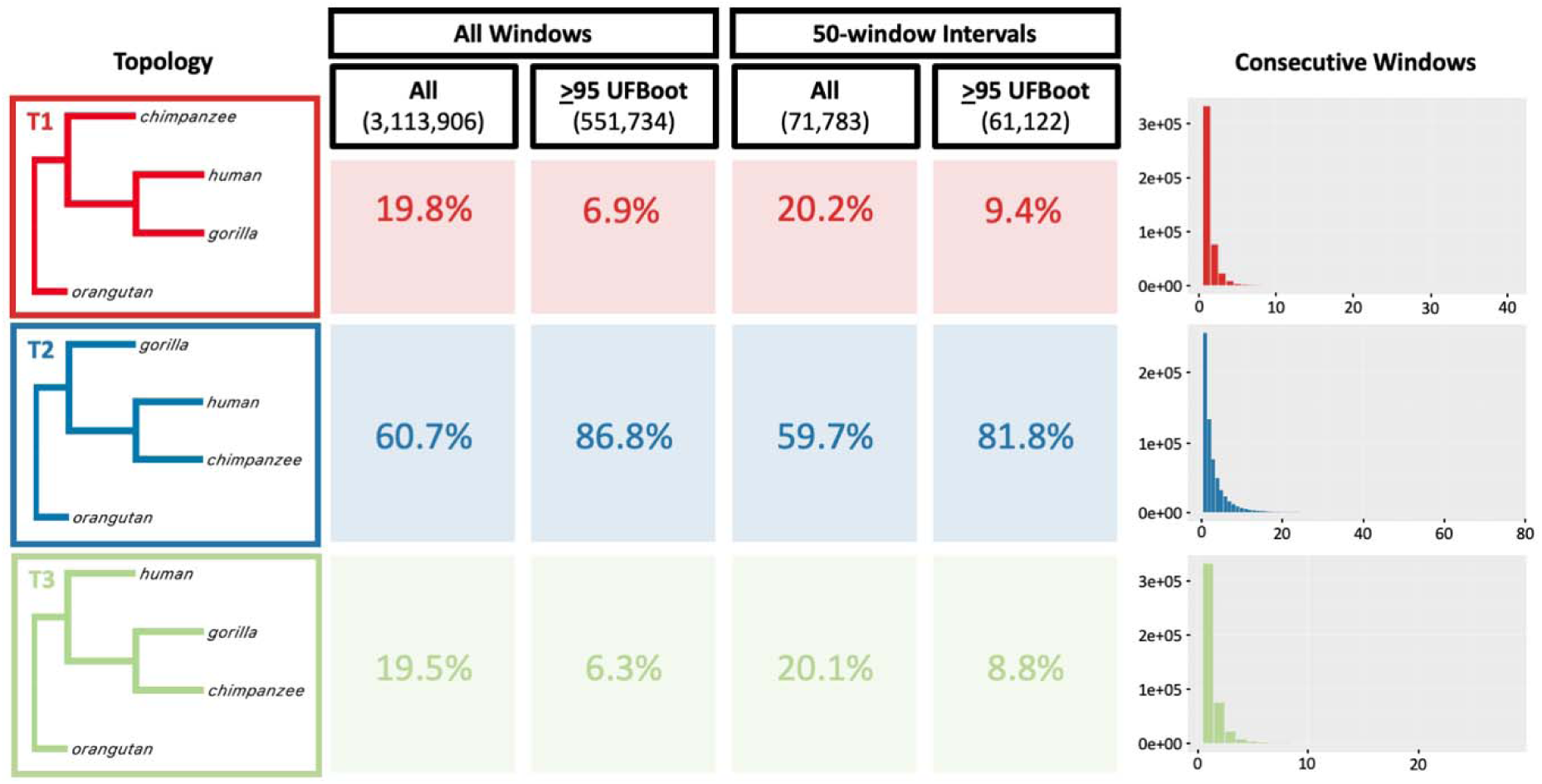
Comparison of topology frequency from great apes’ genomes when including all windows (first and second columns) and window trees with a minimum of 50-window interval (third and fourth columns) based on stepwise non-overlapping windows using the best window size for each chromosome (Table S8). The rightmost panels show the number of consecutive windows that recovers each topology from stepwise NOW considering all window trees. The number under each method reflects the total number of windows. Topologies are visualised using FigTree v1.4.4 (http://tree.bio.ed.ac.uk/software/figtree) without branch lengths.

We further investigated the impact of window size on empirical tree distributions by comparing the distribution of tree topologies across the whole range of window sizes (Fig. 5). At small window sizes, such proportions are likely to be highly affected by stochastic error, while at large window sizes they are likely to be affected by concatenation (the inclusion of one or more recombination breakpoints in a single locus). Figure 5 shows the genome-wide proportion of the major topology (grouping human and chimpanzee) calculated from all trees (Fig. 5a) and highly supported trees (i.e., trees with ≥95 average UFBoot support, Fig. 5b) across all window sizes. Consistent with the effect of stochastic error, the inferred proportion of the major topology is much lower across all trees than it is for highly-supported trees, particularly at smaller window sizes (compare Fig. 5a to Fig. 5b, also Fig. S26-27). On the other hand, the proportion of the major topology shows a monotonic increase in most chromosomes as the window size increases, which follows the effect of concatenation (see Discussion for more details).

**Figure 5.**
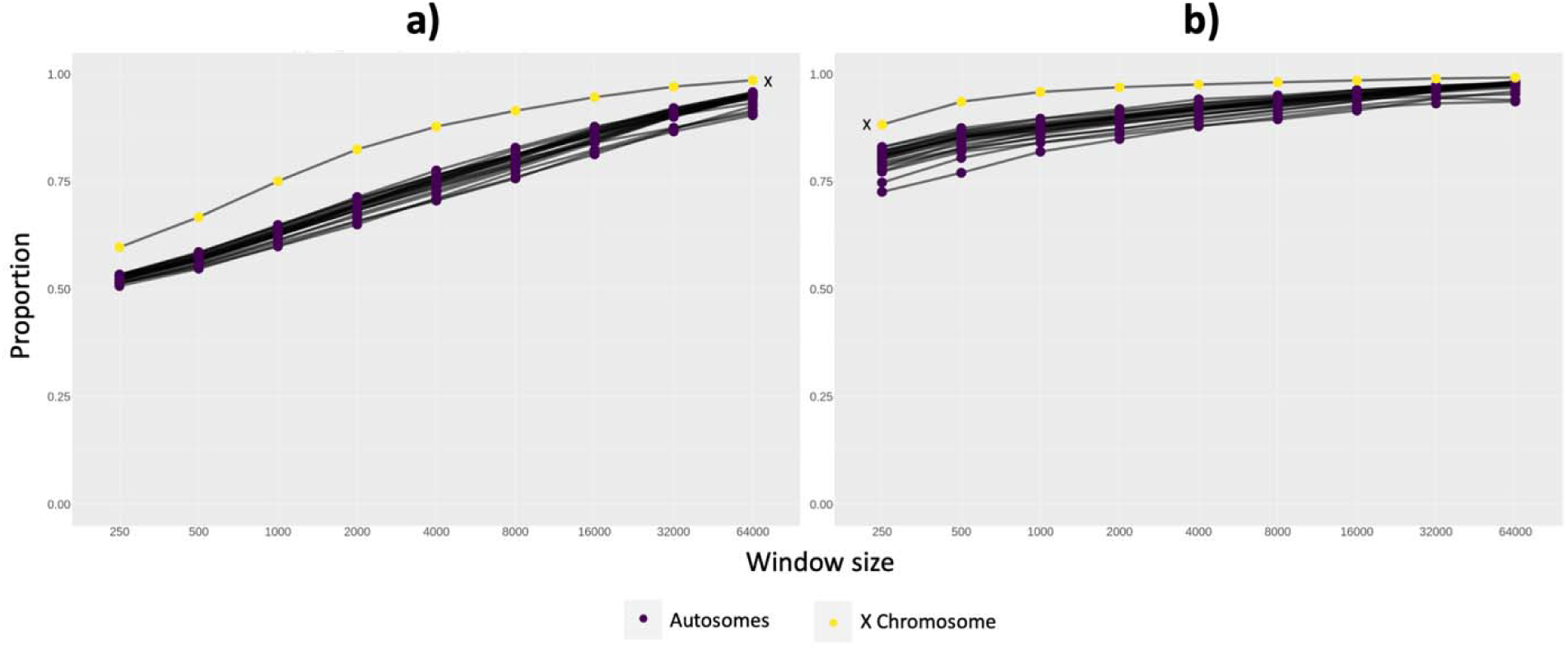
Proportion of major topology of great apes (grouping human and chimpanzee) from stepwise non-overlapping windows based on (a) all window trees, and (b) window trees with ≥95 average UFBoot support. Black lines connect the same chromosome across window sizes.

## Discussion

The non-overlapping window method is commonly used in phylogenomic studies to infer local histories along a set of genomes. However, the selection of the window size is often arbitrary and might not closely reflect the optimal size to extract non-recombining regions, affecting the inference of tree topologies in different genomic regions. In order to address this problem, we simulated chromosomes that mimic the empirical alignments of *erato*-*sara Heliconius* butterflies (Edelman et al. 2019) with varying recombination rates. We then analysed these simulated datasets using different non-overlapping window sizes and showed that the AIC (but not the BIC) is a strong correlate of two different measures of accuracy: the proportion of sites in the genome assigned to the correct tree topology (i.e., site accuracy), and the distribution of tree topologies recovered from the analysis (i.e., RMSE). We then developed a stepwise approach that uses the AIC to select the optimal window size for empirical alignments and applied this to empirical datasets from butterflies and great apes.

Applying the stepwise non-overlapping windows method on *erato-sara Heliconius*’ genomes from Edelman et al. (2019), the best window size for each chromosome ranges from ≤125bp to 250bp (Table S4), smaller than the predicted linkage blocks in *H. erato* by Tobler et al. (2005) and Counterman et al. <u>(2010)</u>. As our dataset does not only include *H. erato* but also other closely-related species, our estimates should include older recombination events that have occurred during the evolution of the group, resulting in c-genes that are shorter than the linkage blocks in any single species if there is a recombination ratchet (Springer and Gatesy 2016). These window sizes are also vastly shorter than those used in the original study (Edelman et al. 2019), which used 10kb and 50kb windows in different analyses. Somewhat reassuringly, the different window-based methods that we and others have applied to this dataset do not significantly change the most common topologies inferred across the genomes of the group, particularly when considering the highly-supported trees (Fig. 3, Table S7). Thus, if the focus is simply on identifying the most common topologies, then the exact choice of window size may not significantly affect the approximation. However, if the *distribution* of the topologies or their locations along the genome are the focus (e.g., in the estimation of the extent of recombination and hybridisation, reconstruction of recombination-based linkage maps, or association between genotype and phenotype), then the selection of window size matters (Table S6-S7, Fig. S21).

For great apes, the best window size for each chromosome ranges from 500bp to 1kb (and 4kb for the mitochondrial genome) (Table S8). These are much longer than the best window sizes from *Heliconius,* likely reflecting the lower average recombination rates in great apes compared to butterflies (Tobler et al. 2005; The International HapMap Consortium 2007; Stevison et al. 2016) due to their smaller effective population sizes (Prado-Martinez et al. 2013; Van Belleghem et al. 2018) despite their older divergence times (Kozak et al. 2015; Reis et al. 2018). Previous work comprising human, chimpanzee, and orangutan has suggested that the length of genomic regions that recover the species phylogeny (grouping human and chimpanzee) ranged from 900bp to 7.8kb, while genomic fragments with alternative genealogies can be very short (<100bp; <u>Hobolth et al., 2011)</u>. On the other hand, Springer and Gatesy (2016) estimates that the mean c-genes for human, chimpanzee, and gorilla to be around ∼109bp. Our results are 5x-10x larger than these estimates, likely due to the lack of phylogenetic signal in shorter windows.

The best window size of 4kb for the mitochondrial genome might be counterintuitive at first – this 16kb genome is inherited as a single locus and recombination is vanishingly rare (though likely not completely absent (Kraytsberg et al. 2004)), suggesting that a single window should be recovered, not four. However, all four of the 4kb windows for the mitochondrial genome have the same topology, and it is simply the branch lengths that differ (Fig. S23). The selection of four windows instead of one likely occurs because of model misspecification, such that allowing for four windows with identical topologies and slightly different branch lengths is a better fit to the data than having two windows or a single window. Similar observations were made by Richards et al. <u>(2018)</u>, who showed that the inferred tree topologies of different regions of the mitochondrial genome often differ because of model misspecification.

Our analyses highlight the challenges of estimating the distribution of tree topologies from whole genome alignments. In the analysis of great apes, there are only three possible topologies. Figure 5 shows that the frequency of the species tree (major) topology (grouping human and chimpanzee) is just 50% when using 250bp windows, but increases to almost 100% when using 64kb windows. This dramatic change occurs because of two competing effects: stochastic error and concatenation. Stochastic error in shorter windows is caused by a lack of phylogenetic signal and tends to push the topologies towards a purely random distribution (i.e., a frequency of 33.3% for the species tree topology). On the other hand, longer windows are more likely to include recombination breakpoints (i.e., concatenation), which is likely to favour the species tree topology because it tends to dominate the phylogenetic signals of the great apes’ genomes. Previous studies have sometimes attempted to ameliorate the effects of stochastic error by analysing ‘highly supported’ trees, for example those with average bootstrap support ≥95%. However, Bryant & Hahn <u>(2020)</u> pointed out that this approach will bias the distribution towards locus trees that agree with the species tree, because coalescent theory dictates that the internal branches of such trees will tend to be longer. This effect can be seen clearly in our data – when we analysed only the trees with average bootstrap support ≥95% (Fig. 5b), the estimated frequency of tree grouping human and chimpanzee is far higher across almost every chromosome than when we analysed all trees (Fig. 5a), and this effect is repeated on the trees from the optimal window sizes (Fig. 4). Nevertheless, these approaches might allow us to estimate a plausible range within which the frequency of the species tree lies. Using the optimal window size for each chromosome, spaced 50-windows apart to prevent multiple counting of the same non-recombining blocks (Fig. 4), the complete distribution of tree topologies suggests that the species tree has a frequency of 60% (with the other two possible topologies having a frequency of 20% each). This estimate can be considered a lower bound since it would be driven down by stochastic error in short windows. When we analysed only those trees from the same set with high bootstrap support, the frequency becomes 82% for the species tree (and 9% each for the alternative topologies). This estimate represents an upper bound since it is biased upwards by the longer internal branches in the loci that match the species tree. The truth is likely to be somewhere in-between, which agrees closely with many previous estimates of this frequency, ranging from 60% to 77% depending on the methods being used (Ebersberger et al. 2007; Vanderpool et al. 2020; Wong et al. 2024).

A number of approaches can be used to further refine the locus trees estimated from the methods we present here, particularly when the best window size is very short and ML methods are limited in their ability to infer a reliable set of locus trees. Such methods include ARG reconstruction using Espalier (Rasmussen and Guo 2023) and multi-tree mixture model from MAST (Wong et al. 2024). All of these methods take different approaches to the same end – the estimation of the topology underlying every site in the alignment. Most also benefit considerably from the initial selection of the optimal window size and the estimate of the tree associated with each window. In conclusion, our AIC-based stepwise non-overlapping method provides a more objective way to select the best window size given the alignments. One key limitation of this method is that only a single window size is selected for each aligned chromosome. Future work will focus on estimating variable window sizes across the genome by dividing long windows and/or joining together two neighbouring windows.

## Supplementary Materials

Supplementary materials will be available online upon the acceptance of the paper.

## Acknowledgements

We thank Matt Hahn from Indiana University for providing constructive feedback during the writing of the manuscript. We also thank Minh Bui, Justin Borevitz, and the members of Lanfear and Bui Lab from Australian National University for comments on the study design and the manuscript.

