## Supplementary File for "Selecting a Window Size for the Analysis of Whole Genome Alignments using AIC"

### **Supplementary Text**

#### **Measuring Key Characteristics of the Empirical Dataset**

To simulate chromosome alignments that match the key characteristics of the data from Edelman et al. (2019), we first measured the summary statistics of the alignment (i.e., chr11) used in that study. We downloaded the whole genome alignment of the *Lepidoptera* group from Dryad (doi: 10.5061/dryad.b7bj832). As the *H. erato* reference genome (*HeraRef*) consisted of 16 sequences for chr11, we converted them individually into MAF using HAL toolkit v2.2 (Hickey et al. 2013):

hal2maf --refGenome HeraRef --targetGenomes Hdem,Hsar,Hhim,Hhsa,Htel,HmelRef --refSequence Herato_chr11_{i} edelman2019_lepidopteraAlignmen.hal {i}.maf

In Edelman et al. (2019), the authors used --refTargets (with BED file as input) to retrieve the MAF blocks. In this study, as we only have one target chromosome, we extracted the blocks using the sequence names, where i represents the sequence number from 1 to 16. Following the original study, we then deleted all MAF blocks with multiple entries from the same taxon using the code provided by the authors (getSingleCopy.py) and sorted the blocks using maf-sort (<https://github.com/UCSantaCruzComputationalGenomicsLab/last>). We then converted each MAF alignment into FASTA and concatenated the alignments using PHAST v1.5 (Hubisz et al. 2011) with the following commands:

$ msa_view {i}.maf -i MAF -m –G 1 > {i}.fa

$ msa_view --aggregate HmelRef,HeraRef,Hhim,Hhsa,Htel,Hdem,Hsar {i}.fa {i+1}.fa ... > concat.fa

We calculated the summary statistics of the concatenated alignment using IQ-TREE2 (Minh et al. 2020) with the following command: iqtree2 -s concat.fa -alninfo. The final 10,454,565bp alignment of chr11 from the 7 focal species had 4.987% informative sites. Based on this analysis, we designed our simulations to produce a 10Mb alignment with approximately 5% informative sites.

Next, we sought to determine the degree of ILS that we should simulate in our dataset. While the precise degree of ILS remains unknown in this clade, we tried to approximate the empirical data in this regard by ensuring that our simulated datasets resulted in roughly the same distribution of tree topologies as observed in the original study. Specifically, Edelman et al. (2019) found that the eight most common tree topologies accounted for roughly 98% of the recovered trees from 50kb sliding windows (and 87.6% for 10kb non-overlapping windows), and that this distribution was almost identical for both the whole genome and chr11 analysed individually. Thus, we also sought to simulate data for which ~98% of the simulated locus trees along the chromosome were concentrated onto approximately 8 different topologies. The precise distribution of tree topologies in our simulation may differ from that in the original study because we incorporated the introgression events estimated in a more recent study (Thawornwattana et al. 2022). Nevertheless, by aiming for a similar distribution of tree topologies, we seek to simulate a dataset that approximates the levels of introgression and ILS in the empirical data.

#### **Simulating Single Chromosome Alignments**

We simulated single chromosome alignments along the tree in Figure S1 using ms (Hudson 2002) and AliSim (Ly-Trong et al. 2022). We chose parameters for these simulations by considering their biological plausibility, and ensuring that they produced simulated chromosomes that match the key characteristics listed above. First, we varied the scaling of divergence times (i.e., conversion from Myr shown in Figure S1 to 4N_e_ generations used in ms) and branch length (i.e., conversion from 4N_e_ generations of the trees produced by ms to the number of substitutions per site used by AliSim). Second, we varied the recombination rates to simulate chromosomes that recover similar sets of topologies with the empirical dataset. The full set of ms simulation parameters is depicted in Figure S1.

##### **Simulating Trees in ms**

We used ms to simulate trees evolving along the species tree shown in Figure S1. To do this, we first had to convert the divergence times from Kozak et al. (2015) to the units of 4N_e_ generations used in ms, which requires estimates of both the effective population size and the generation time for the clade of interest. The effective population size of *erato-sara* clade has been estimated to be of the order of 10^6^ to 10^7^ (Van Belleghem et al. 2018), and the generation time has been estimated to be around one to three months (Kronforst 2008; Van Belleghem et al. 2018). These figures imply that 4N_e_ generations equates to somewhere between 0.3 to 10Myr. We initially set the recombination rate (ρ) to a value that resulted in approximately 1,000 loci being simulated along the chromosome, to generate a large enough number of loci to determine the distribution of the tree topologies (this number was later varied to generate simulation conditions where the optimal window size differs).

Initial analyses revealed that setting 4N_e_ generations to be larger than or equal to 2Myr led to at least 1% of trees placing *H. melpomene* as ingroup species due to ILS. Thus, we only considered ms scaling factor between 0.1 (the lower bound from the previous estimation) and 1Myr. From 100 replicates (Table S1), ms scaling factor of 0.75Myr has the closest average number of topologies to the empirical dataset. Thus, we selected this scaling factor for our simulation. In order to simulate a 10Mb chromosome with ρ = 17 and 4N_e_ generations = 0.75Myr, we used the following command:

ms 7 1 -T -I 7 1 1 1 1 1 1 1 -ej 2.13 2 1 -ej 5.60 3 1 -ej 7.07 4 1 -ej 6.67 5 6 -ej 8.13 6 1 -ej 15.47 7 1 -es 0.21 2 0.966 -ej 0.21 8 1 -es 0.21 1 0.796 -ej 0.21 9 2 -es 5.52 3 0.115 -ej 5.52 10 4 -es 6.84 6 0.778 -ej 6.84 11 4 -es 6.84 4 0.853 -ej 6.84 12 6 -r 17 10000000

In this command, the first two numbers represent the number of taxa and sampling, respectively. -T set the ms output as locus trees, while the -I 7 1 1 1 1 1 1 1 argument indicates seven populations with one sample per population. Each -ej flag represents a speciation event, which is followed by three numbers describing the speciation timing and the two resulting populations. In this case, each *Heliconius* species represented one population: 1: *H. himera*, 2: *H. erato*, 3: *H. hecalesia*, 4: *H. telesiphe*, 5: *H. sara*, 6: *H. demeter*, and 7: *H. melpomene* (outgroup). The -es flags represent introgression events, followed by introgression timing, recipient taxon, and inheritance probability (i.e., 1 – rate of introgression), whereas the subsequent -ej flag tells the new population (second number) to coalesce with the donor taxon (third number). The recombination rate was set by the -r flag, followed by alignment length.

##### **Simulating Alignments with AliSim**

We used the set of locus trees and their respective lengths from ms to simulate DNA alignments with AliSim (Ly-Trong et al. 2022). To do this, we first converted the branch lengths from the ms output (in units of 4N_e_ generations) to units of substitutions per site. In principle, this scaling factor could be estimated using estimates of the effective population size, generation time, and per-generation mutation rate from *Heliconius* (which Keightley et al. (2015) and Van Belleghem et al. (2018) have estimated to be of the order of 2x10^-9^ substitutions per site per generation). Based on the effective population size and generation time from the previous paragraph, the branch length scaling factor would be around 0.004 to 0.12 substitutions per site per 4N_e_ generations. Within this range, we sought to select a scaling factor that would lead to an alignment with approximately 5% informative sites as observed in the empirical alignment.

For each set of locus trees generated by ms, we simulated one alignment for each tree with its respective length using a Jukes-Cantor (JC (Jukes and Cantor 1969)) model of evolution in AliSim. We did not incorporate indels or gaps into the alignment because we did not expect the distributions of indels and gaps to fundamentally affect the appropriate window size for a chromosome. We tried a range of branch length scaling factors and, based on the average proportion of informative sites across 100 independent 10Mb simulated alignments (Table S2), we selected a branch length scaling factor of 0.005.

##### **Simulating Datasets with Different Recombination Rates**

Next, we sought to simulate datasets with a wide range of recombination rates, such that the best non-overlapping window size should vary among simulation conditions. To do this, we varied the recombination rate from 0 to 2000 (the highest recombination rate for which ms finished simulating the locus trees in <1 hour using a server with Intel(R) Xeon(R) CPU E5-2690 v4 @2.60GHz). These extremes resulted in simulating one locus with a size of 10Mb (i.e., the entire simulated chromosome) that evolved along one unique tree topology when the recombination rate is zero, to ~126,000 loci with an average size of 80bp that evolved along approximately 70 tree topologies when the recombination rate is 2000 (Table S3). These extremes include a broad range of conditions and represent different challenges for non-overlapping window analyses. Following the same ms command as mentioned above but varying the recombination rate, we selected four different recombination rates (0, 20, 200, 2000) and generated 10 replicates for each, for a total of 40 simulated chromosomes, hitherto referred to as scenario with a medium level of ILS.

##### **Simulating Datasets with Different Degrees of ILS**

In order to validate the consistency of the results across a range of different conditions, we simulated two more scenarios where the degree of ILS was altered, using the smallest and largest possible values of the ms scaling factor (Table S1). The analyses exactly followed the previous analyses, except that the ms and branch length scaling factors were set to be 1.0 and 0.007 (for datasets with high ILS), and 0.1 and 0.0007 (for datasets with low ILS). We adjusted the branch length scale to achieve 5% informative sites, and the recombination rates so that the ms simulation completed in under one hour. The highest recombination rate for high ILS simulations was 2500, resulting in ~100 unique topologies (around 1.4x the number of topologies from the scenario with medium ILS). On the other hand, low ILS simulations had a maximum recombination rate of 200, resulting in ~10 unique topologies (around 0.14x the number of topologies from the scenario with medium ILS).

### **Supplementary Tables and Figures**

**Table S1.** Average numbers of locus and unique topology across different ms scaling factors. ms scale = ms scaling factor to the unit of 4N_e_ generations; ρ = simulated recombination rate to generate ~1,000 loci; #locus = average number of locus; #topology = average number of unique topology that accounts for ≥98.5% of all loci. The averages are derived from 100 simulated 10Mb alignment.

|  | **ms scale (Myr)** | **ρ** | **#locus** | **#topology** |
| --- | --- | --- | --- | --- |
| 1 | 1.00 | 22.5 | 1087.11 | 11.95 |
| 2 | 0.95 | 21.0 | 1061.39 | 10.95 |
| 3 | 0.90 | 20.0 | 1068.47 | 10.27 |
| 4 | 0.85 | 19.0 | 1073.56 | 9.92 |
| 5 | 0.80 | 18.0 | 1073.91 | 9.13 |
| 6 | 0.75 | 17.0 | 1073.49 | 7.88 |
| 7 | 0.70 | 16.0 | 1080.43 | 7.46 |
| 8 | 0.65 | 15.0 | 1089.68 | 7.15 |
| 9 | 0.60 | 14.0 | 1092.07 | 5.73 |
| 10 | 0.55 | 12.5 | 1065.89 | 5.82 |
| 11 | 0.50 | 11.5 | 1072.43 | 5.34 |
| 12 | 0.25 | 6.0 | 1095.38 | 3.44 |
| 13 | 0.10 | 2.5 | 1129.66 | 1.66 |

**Table S2.** Average proportion of informative sites across different branch length (bl) scaling factors. ms scale = ms scaling factor to the unit of 4N_e_ generations; bl scale = branch length scaling factor to the unit of substitution per site. The average is derived from 100 simulated 10Mb alignment.

|  | **ms scale (Myr)** | **bl scale** | **informative sites (%)** |
| --- | --- | --- | --- |
| 1 | 0.75 | 0.010 | 10.610 |
| 2 |  | 0.009 | 9.521 |
| 3 |  | 0.008 | 8.381 |
| 4 |  | 0.007 | 7.215 |
| 5 |  | 0.006 | 6.096 |
| 6 |  | 0.005 | 5.085 |
| 7 |  | 0.004 | 3.981 |
| 8 |  | 0.003 | 2.883 |
| 9 |  | 0.002 | 1.913 |
| 10 |  | 0.001 | 0.918 |

**Table S3**. Average statistics of the simulated alignments and their respective best window sizes across replicates. ρ = simulated recombination rate; locus length = average length of locus across 10 replicates based on ms simulation (*raw*), merged when consecutive loci have the same topology and branch length (*top+bl*), and merged when consecutive loci have the same topology only (*top*); #locus = average number of locus from ms simulation (*raw*) across 10 replicates; #topology = average number of unique locus topology across 10 replicates; best window size = window size(s) with the highest site accuracy and the lowest RMSE according to the non-overlapping windows (NOW), and median size selected by the AIC and BIC across 10 replicates. Numbers in brackets reflect median of the locus lengths.

| **ILS level** | **ρ** | **locus length (bp)** | | | **#locus** | **#topology** | **best window size (bp)** | | |
| --- | --- | --- | --- | --- | --- | --- | --- | --- | --- |
|  |  | **raw** | **top+bl** | **top** |  |  | **NOW** | **AIC** | **BIC** |
| low | 0 | 10,000,000 | 10,000,000 | 10,000,000 | 1 | 1 | ≥500,000 | 10,000,000 | 10,000,000 |
|  | 2 | 10,915.5  (10,935.9) | 702,615.4  (645,833.3) | 10,000,000  (10,000,000) | 917.1 | 1.0 | ≥5,000 | 1,000,000 | 10,000,000 |
|  | 20 | 1,102.8  (1,099.8) | 58,561.0  (58,487.5) | 3,075,260.0  (1,010,101.0) | 9069.2 | 3.4 | 20,000 | 50,000 | 100,000 |
|  | 200 | 111.6  (111.5) | 6,070.7  (6,123.7) | 102,011.6  (93,457.9) | 89,606.7 | 11.1 | 10,000 | 10,000 | 50,000 |
| medium | 0 | 10,000,000 | 10,000,000 | 10,000,000 | 1 | 1 | ≥500,000 | 10,000,000 | 10,000,000 |
|  | 20 | 7,880.3  (7,782.2) | 62,797.8  (60,276.0) | 445,152.4  (444,664.0) | 1,270.7 | 9.9 | 20,000 | 20,000 | 100,000 |
|  | 200 | 789.3  (789.8) | 6,530.0  (6,474.8) | 39,858.4  (39,685.0) | 12,669.5 | 35.6 | 5,000 | 10,000 | 50,000 |
|  | 2000 | 79.3  (79.3) | 651.3  (653.1) | 4,216.3  (4,226.6) | 126,113.2 | 71.4 | 2,000 | 2,000 | 20,000 |
| high | 0 | 10,000,000 | 10,000,000 | 10,000,000 | 1 | 1 | ≥50,000 | 10,000,000 | 10,000,000 |
|  | 25 | 8,306.7  (8,316.0) | 58,318.7  (58,874.5) | 481,287.9  (385,185.2) | 1,204.9 | 11.9 | 20,000 | 20,000 | 100,000 |
|  | 250 | 831.7  (832.9) | 5,505.5  (5,473.5) | 27,208.9  (26,846.1) | 12,024.8 | 51.5 | 5,000 | 5,000 | 20,000 |
|  | 2500 | 83.2  (83.2) | 536.9  (536.6) | 2,678.3  (2,637.6) | 120,173.0 | 101.2 | 2,000 | 2,000 | 20,000 |

**Table S4.** Best window size for each *erato-sara Heliconius* butterflies’ chromosome based on stepwise non-overlapping windows. Length is in the units of base pairs. Initial length = length of chromosome before filtering; 64kb length = length of chromosome after divided into 64kb non-overlapping windows; final length = length of chromosome when the best window size is found; best window size = window size with the best AIC score. Percentage of final length refers to the initial length of the alignment.

| **chr** | **initial length** | **64kb length** | **final length** | **best window size** |
| --- | --- | --- | --- | --- |
| 1 | 16,810,322 | 16,768,000 | 12,690,000 (75.49%) | 250 |
| 2 | 8,186,463 | 8,064,000 | 4,049,000 (49.46%) | ≤125 |
| 3 | 9,505,348 | 9,472,000 | 5,711,250 (60.09%) | ≤125 |
| 4 | 8,396,326 | 8,384,000 | 4,777,750 (56.90%) | ≤125 |
| 5 | 8,937,605 | 8,896,000 | 5,355,250 (59.92%) | ≤125 |
| 6 | 13,137,424 | 13,120,000 | 9,149,000 (69.64%) | 250 |
| 7 | 13,846,482 | 13,824,000 | 10,108,500 (73.00%) | 250 |
| 8 | 8,351,654 | 8,320,000 | 4,878,000 (58.41%) | ≤125 |
| 9 | 7,667,912 | 7,616,000 | 4,416,250 (57.59%) | ≤125 |
| 10 | 18,909,427 | 18,880,000 | 13,259,000 (70.12%) | 250 |
| 11 | 10,454,565 | 10,432,000 | 6,524,250 (62.41%) | ≤125 |
| 12 | 16,081,187 | 16,064,000 | 11,245,500 (69.93%) | 250 |
| 13 | 16,998,850 | 16,960,000 | 12,081,500 (71.07%) | 250 |
| 14 | 8,407,831 | 8,384,000 | 4,746,250 (56.45%) | ≤125 |
| 15 | 9,302,932 | 9,216,000 | 4,853,250 (52.17%) | ≤125 |
| 16 | 9,322,555 | 9,280,000 | 5,434,750 (58.30%) | ≤125 |
| 17 | 14,100,382 | 14,016,000 | 9,905,000 (70.25%) | 250 |
| 18 | 16,500,162 | 16,448,000 | 11,987,500 (72.65%) | 250 |
| 19 | 16,952,300 | 16,896,000 | 11,335,000 (66.86%) | 250 |
| 20 | 14,467,297 | 14,464,000 | 10,687,000 (73.87%) | 250 |
| 21 | 13,727,181 | 13,696,000 | 8,570,500 (62.44%) | 250 |

**Table S5.** Summary table of stepwise non-overlapping windows on *Heliconius* butterflies’ genomes. Alignment length reflects the total length of the concatenated windows (bp) and its percentage to the original alignment (%). L = long window size; S = short window size; #window = number of windows with ≥95 average UFBoot support; #topology = number of unique topologies with ≥95 average UFBoot support.

| **chr** | **step** | **window sizes (bp)** | | **AIC (L–S)** | **alignment length** | | **window trees with ≥95 UFBoot support** | | | |
| --- | --- | --- | --- | --- | --- | --- | --- | --- | --- | --- |
|  |  | **L** | **S** |  | **bp** | **%** | **L** | | **S** | |
|  |  |  |  |  |  |  | **#window** | **#topology** | **#window** | **#topology** |
| 1 | 1 | 64000 | 32000 | -49474.972 | 16768000 | 99.748 | 175 | 6 | 269 | 10 |
|  | 2 | 32000 | 16000 | -63582.861 | 16768000 | 99.748 | 269 | 10 | 458 | 10 |
|  | 3 | 16000 | 8000 | -94498.921 | 16752000 | 99.653 | 458 | 10 | 685 | 13 |
|  | 4 | 8000 | 4000 | -141487.366 | 16680000 | 99.225 | 682 | 13 | 813 | 20 |
|  | 5 | 4000 | 2000 | -190974.206 | 16392000 | 97.512 | 813 | 20 | 736 | 30 |
|  | 6 | 2000 | 1000 | -231149.387 | 15666000 | 93.193 | 728 | 30 | 447 | 28 |
|  | 7 | 1000 | 500 | -282700.605 | 14356000 | 85.400 | 444 | 28 | 223 | 27 |
|  | 8 | 500 | 250 | -306751.431 | 12690000 | 75.489 | 225 | 26 | 86 | 20 |
|  | 9 | 250 | 125 | 34279.475 | 11225000 | 66.774 | 84 | 19 | 28 | 15 |
| 2 | 1 | 64000 | 32000 | -20698.652 | 8064000 | 98.504 | 114 | 4 | 204 | 5 |
|  | 2 | 32000 | 16000 | -27531.477 | 8032000 | 98.113 | 205 | 5 | 318 | 8 |
|  | 3 | 16000 | 8000 | -47037.878 | 7904000 | 96.550 | 314 | 8 | 477 | 12 |
|  | 4 | 8000 | 4000 | -63904.900 | 7728000 | 94.400 | 477 | 11 | 568 | 17 |
|  | 5 | 4000 | 2000 | -84914.013 | 7356000 | 89.856 | 560 | 17 | 526 | 18 |
|  | 6 | 2000 | 1000 | -103901.424 | 6750000 | 82.453 | 522 | 18 | 332 | 20 |
|  | 7 | 1000 | 500 | -121293.274 | 5846000 | 71.411 | 328 | 20 | 148 | 22 |
|  | 8 | 500 | 250 | -119066.995 | 4882000 | 59.635 | 145 | 19 | 52 | 14 |
|  | 9 | 250 | 125 | -11739.172 | 4049000 | 49.460 | 54 | 14 | 16 | 6 |
| 3 | 1 | 64000 | 32000 | -24376.406 | 9472000 | 99.649 | 119 | 3 | 190 | 8 |
|  | 2 | 32000 | 16000 | -30838.113 | 9376000 | 98.639 | 190 | 8 | 288 | 9 |
|  | 3 | 16000 | 8000 | -47786.917 | 9344000 | 98.303 | 288 | 9 | 409 | 9 |
|  | 4 | 8000 | 4000 | -77350.650 | 9232000 | 97.124 | 407 | 9 | 417 | 14 |
|  | 5 | 4000 | 2000 | -103047.917 | 9004000 | 94.726 | 415 | 14 | 323 | 14 |
|  | 6 | 2000 | 1000 | -127917.181 | 8512000 | 89.550 | 322 | 14 | 181 | 16 |
|  | 7 | 1000 | 500 | -163895.241 | 7628000 | 80.250 | 183 | 16 | 97 | 16 |
|  | 8 | 500 | 250 | -182717.521 | 6608500 | 69.524 | 100 | 16 | 40 | 16 |
|  | 9 | 250 | 125 | -7149.424 | 5711250 | 60.085 | 38 | 14 | 13 | 8 |
| 4 | 1 | 64000 | 32000 | -25626.186 | 8384000 | 99.853 | 107 | 5 | 181 | 8 |
|  | 2 | 32000 | 16000 | -31339.994 | 8384000 | 99.853 | 181 | 8 | 255 | 10 |
|  | 3 | 16000 | 8000 | -49441.395 | 8352000 | 99.472 | 255 | 10 | 309 | 10 |
|  | 4 | 8000 | 4000 | -65344.142 | 8224000 | 97.948 | 309 | 10 | 318 | 12 |
|  | 5 | 4000 | 2000 | -91453.131 | 7980000 | 95.042 | 316 | 12 | 247 | 18 |
|  | 6 | 2000 | 1000 | -116719.793 | 7418000 | 88.348 | 243 | 18 | 113 | 17 |
|  | 7 | 1000 | 500 | -149381.867 | 6604000 | 78.653 | 114 | 17 | 64 | 11 |
|  | 8 | 500 | 250 | -163074.197 | 5603000 | 66.732 | 64 | 11 | 31 | 12 |
|  | 9 | 250 | 125 | -18251.510 | 4777750 | 56.903 | 30 | 11 | 6 | 6 |
| 5 | 1 | 64000 | 32000 | -23898.664 | 8896000 | 99.534 | 114 | 4 | 195 | 6 |
|  | 2 | 32000 | 16000 | -28888.389 | 8896000 | 99.534 | 195 | 6 | 324 | 9 |
|  | 3 | 16000 | 8000 | -51299.781 | 8800000 | 98.460 | 322 | 9 | 384 | 11 |
|  | 4 | 8000 | 4000 | -75017.507 | 8704000 | 97.386 | 383 | 11 | 378 | 16 |
|  | 5 | 4000 | 2000 | -98929.299 | 8484000 | 94.925 | 377 | 15 | 268 | 19 |
|  | 6 | 2000 | 1000 | -132289.945 | 8062000 | 90.203 | 270 | 20 | 150 | 16 |
|  | 7 | 1000 | 500 | -167521.726 | 7245000 | 81.062 | 148 | 15 | 60 | 14 |
|  | 8 | 500 | 250 | -175992.494 | 6268000 | 70.131 | 57 | 12 | 21 | 10 |
|  | 9 | 250 | 125 | -13706.988 | 5355250 | 59.918 | 21 | 10 | 11 | 7 |
| 6 | 1 | 64000 | 32000 | -35511.784 | 13120000 | 99.867 | 154 | 5 | 251 | 8 |
|  | 2 | 32000 | 16000 | -47154.419 | 13120000 | 99.867 | 251 | 8 | 382 | 8 |
|  | 3 | 16000 | 8000 | -73218.312 | 13104000 | 99.746 | 382 | 8 | 570 | 13 |
|  | 4 | 8000 | 4000 | -103312.134 | 12928000 | 98.406 | 568 | 13 | 659 | 19 |
|  | 5 | 4000 | 2000 | -138078.476 | 12628000 | 96.122 | 656 | 19 | 539 | 22 |
|  | 6 | 2000 | 1000 | -171370.909 | 11856000 | 90.246 | 533 | 21 | 323 | 25 |
|  | 7 | 1000 | 500 | -210774.211 | 10652000 | 81.081 | 315 | 25 | 161 | 22 |
|  | 8 | 500 | 250 | -218177.263 | 9149000 | 69.641 | 157 | 21 | 62 | 19 |
|  | 9 | 250 | 125 | 7632.817 | 7845500 | 59.719 | 63 | 19 | 17 | 9 |
| 7 | 1 | 64000 | 32000 | -41417.460 | 13824000 | 99.838 | 160 | 8 | 254 | 10 |
|  | 2 | 32000 | 16000 | -59536.204 | 13824000 | 99.838 | 254 | 10 | 397 | 10 |
|  | 3 | 16000 | 8000 | -85932.365 | 13792000 | 99.607 | 396 | 10 | 540 | 14 |
|  | 4 | 8000 | 4000 | -125661.184 | 13696000 | 98.913 | 538 | 14 | 647 | 20 |
|  | 5 | 4000 | 2000 | -159811.978 | 13472000 | 97.295 | 644 | 20 | 553 | 25 |
|  | 6 | 2000 | 1000 | -191502.611 | 12744000 | 92.038 | 548 | 25 | 342 | 21 |
|  | 7 | 1000 | 500 | -237269.417 | 11589000 | 83.696 | 340 | 21 | 180 | 23 |
|  | 8 | 500 | 250 | -242447.025 | 10108500 | 73.004 | 181 | 23 | 60 | 17 |
|  | 9 | 250 | 125 | 21665.512 | 8773000 | 63.359 | 61 | 17 | 22 | 9 |
| 8 | 1 | 64000 | 32000 | -19224.729 | 8320000 | 99.621 | 114 | 3 | 199 | 6 |
|  | 2 | 32000 | 16000 | -29124.113 | 8256000 | 98.855 | 198 | 6 | 334 | 8 |
|  | 3 | 16000 | 8000 | -44484.522 | 8208000 | 98.280 | 335 | 8 | 478 | 11 |
|  | 4 | 8000 | 4000 | -70015.300 | 8144000 | 97.514 | 477 | 11 | 505 | 14 |
|  | 5 | 4000 | 2000 | -93484.034 | 7880000 | 94.353 | 502 | 13 | 347 | 16 |
|  | 6 | 2000 | 1000 | -122000.575 | 7464000 | 89.372 | 344 | 16 | 170 | 12 |
|  | 7 | 1000 | 500 | -151842.123 | 6652000 | 79.649 | 165 | 11 | 91 | 16 |
|  | 8 | 500 | 250 | -164469.232 | 5693500 | 68.172 | 89 | 15 | 28 | 9 |
|  | 9 | 250 | 125 | -25253.051 | 4878000 | 58.408 | 31 | 10 | 16 | 6 |
| 9 | 1 | 64000 | 32000 | -16926.971 | 7616000 | 99.323 | 104 | 3 | 184 | 6 |
|  | 2 | 32000 | 16000 | -23577.271 | 7616000 | 99.323 | 184 | 6 | 274 | 7 |
|  | 3 | 16000 | 8000 | -41429.491 | 7600000 | 99.114 | 274 | 7 | 388 | 8 |
|  | 4 | 8000 | 4000 | -60965.586 | 7504000 | 97.862 | 388 | 8 | 400 | 12 |
|  | 5 | 4000 | 2000 | -85818.267 | 7328000 | 95.567 | 400 | 12 | 263 | 18 |
|  | 6 | 2000 | 1000 | -109804.969 | 6842000 | 89.229 | 262 | 19 | 137 | 15 |
|  | 7 | 1000 | 500 | -141392.167 | 6136000 | 80.022 | 137 | 16 | 64 | 14 |
|  | 8 | 500 | 250 | -145409.624 | 5259000 | 68.585 | 62 | 14 | 28 | 12 |
|  | 9 | 250 | 125 | -17955.228 | 4416250 | 57.594 | 29 | 12 | 10 | 8 |
| 10 | 1 | 64000 | 32000 | -50607.680 | 18880000 | 99.844 | 215 | 7 | 344 | 10 |
|  | 2 | 32000 | 16000 | -63257.218 | 18880000 | 99.844 | 344 | 10 | 531 | 14 |
|  | 3 | 16000 | 8000 | -98172.536 | 18864000 | 99.760 | 531 | 14 | 812 | 20 |
|  | 4 | 8000 | 4000 | -145130.211 | 18736000 | 99.083 | 810 | 20 | 918 | 25 |
|  | 5 | 4000 | 2000 | -196267.193 | 18244000 | 96.481 | 910 | 25 | 726 | 32 |
|  | 6 | 2000 | 1000 | -238544.198 | 17072000 | 90.283 | 724 | 32 | 445 | 32 |
|  | 7 | 1000 | 500 | -285027.116 | 15306000 | 80.944 | 443 | 31 | 216 | 26 |
|  | 8 | 500 | 250 | -297461.083 | 13259000 | 70.118 | 214 | 25 | 89 | 15 |
|  | 9 | 250 | 125 | 51143.472 | 11530500 | 60.978 | 92 | 15 | 31 | 8 |
| 11 | 1 | 64000 | 32000 | -28474.591 | 10432000 | 99.784 | 120 | 5 | 201 | 7 |
|  | 2 | 32000 | 16000 | -48752.613 | 10400000 | 99.478 | 201 | 7 | 317 | 9 |
|  | 3 | 16000 | 8000 | -65133.132 | 10368000 | 99.172 | 317 | 9 | 427 | 12 |
|  | 4 | 8000 | 4000 | -87057.478 | 10312000 | 98.636 | 427 | 12 | 455 | 17 |
|  | 5 | 4000 | 2000 | -123334.427 | 10144000 | 97.029 | 453 | 17 | 359 | 22 |
|  | 6 | 2000 | 1000 | -152983.515 | 9594000 | 91.769 | 359 | 22 | 201 | 21 |
|  | 7 | 1000 | 500 | -181804.559 | 8626000 | 82.509 | 202 | 21 | 96 | 22 |
|  | 8 | 500 | 250 | -186879.888 | 7524500 | 71.973 | 97 | 22 | 44 | 14 |
|  | 9 | 250 | 125 | -10241.757 | 6524250 | 62.406 | 43 | 14 | 13 | 7 |
| 12 | 1 | 64000 | 32000 | -46310.466 | 16064000 | 99.893 | 184 | 6 | 301 | 9 |
|  | 2 | 32000 | 16000 | -60955.978 | 16064000 | 99.893 | 301 | 9 | 460 | 9 |
|  | 3 | 16000 | 8000 | -88290.088 | 15968000 | 99.296 | 459 | 9 | 665 | 15 |
|  | 4 | 8000 | 4000 | -125850.907 | 15760000 | 98.003 | 660 | 15 | 750 | 17 |
|  | 5 | 4000 | 2000 | -164515.890 | 15384000 | 95.665 | 743 | 17 | 616 | 25 |
|  | 6 | 2000 | 1000 | -199116.755 | 14430000 | 89.732 | 608 | 23 | 338 | 24 |
|  | 7 | 1000 | 500 | -242121.008 | 12980000 | 80.715 | 334 | 25 | 160 | 30 |
|  | 8 | 500 | 250 | -254579.043 | 11245500 | 69.930 | 161 | 30 | 85 | 21 |
|  | 9 | 250 | 125 | 29286.631 | 9746000 | 60.605 | 85 | 21 | 27 | 13 |
| 13 | 1 | 64000 | 32000 | -38828.140 | 16960000 | 99.771 | 181 | 5 | 323 | 8 |
|  | 2 | 32000 | 16000 | -57137.797 | 16928000 | 99.583 | 323 | 8 | 507 | 12 |
|  | 3 | 16000 | 8000 | -85991.711 | 16896000 | 99.395 | 505 | 12 | 726 | 15 |
|  | 4 | 8000 | 4000 | -126619.101 | 16680000 | 98.124 | 721 | 15 | 925 | 15 |
|  | 5 | 4000 | 2000 | -172790.392 | 16216000 | 95.395 | 922 | 15 | 778 | 26 |
|  | 6 | 2000 | 1000 | -210345.090 | 15308000 | 90.053 | 761 | 25 | 494 | 29 |
|  | 7 | 1000 | 500 | -262705.419 | 13860000 | 81.535 | 493 | 29 | 227 | 30 |
|  | 8 | 500 | 250 | -285605.070 | 12081500 | 71.072 | 228 | 27 | 93 | 19 |
|  | 9 | 250 | 125 | 37207.670 | 10580500 | 62.242 | 93 | 21 | 22 | 12 |
| 14 | 1 | 64000 | 32000 | -17042.403 | 8384000 | 99.717 | 110 | 4 | 194 | 7 |
|  | 2 | 32000 | 16000 | -30383.203 | 8288000 | 98.575 | 194 | 7 | 298 | 11 |
|  | 3 | 16000 | 8000 | -44525.607 | 8160000 | 97.052 | 296 | 11 | 415 | 13 |
|  | 4 | 8000 | 4000 | -73402.879 | 8112000 | 96.481 | 417 | 13 | 411 | 14 |
|  | 5 | 4000 | 2000 | -92883.360 | 7840000 | 93.246 | 411 | 14 | 291 | 16 |
|  | 6 | 2000 | 1000 | -121675.253 | 7388000 | 87.870 | 290 | 16 | 163 | 20 |
|  | 7 | 1000 | 500 | -146785.529 | 6574000 | 78.189 | 161 | 19 | 66 | 13 |
|  | 8 | 500 | 250 | -148395.702 | 5581000 | 66.379 | 65 | 13 | 33 | 15 |
|  | 9 | 250 | 125 | -10162.100 | 4746250 | 56.450 | 34 | 15 | 11 | 6 |
| 15 | 1 | 64000 | 32000 | -21407.502 | 9216000 | 99.066 | 110 | 7 | 177 | 10 |
|  | 2 | 32000 | 16000 | -29002.174 | 9120000 | 98.034 | 176 | 10 | 274 | 9 |
|  | 3 | 16000 | 8000 | -46498.938 | 8912000 | 95.798 | 272 | 9 | 355 | 12 |
|  | 4 | 8000 | 4000 | -71446.709 | 8728000 | 93.820 | 354 | 12 | 377 | 17 |
|  | 5 | 4000 | 2000 | -99140.872 | 8380000 | 90.079 | 378 | 17 | 326 | 20 |
|  | 6 | 2000 | 1000 | -121606.935 | 7812000 | 83.974 | 321 | 20 | 184 | 21 |
|  | 7 | 1000 | 500 | -143307.301 | 6849000 | 73.622 | 181 | 18 | 84 | 17 |
|  | 8 | 500 | 250 | -142516.331 | 5762500 | 61.943 | 86 | 17 | 30 | 11 |
|  | 9 | 250 | 125 | -10929.092 | 4853250 | 52.169 | 30 | 11 | 3 | 2 |
| 16 | 1 | 64000 | 32000 | -22304.790 | 9280000 | 99.544 | 125 | 5 | 212 | 6 |
|  | 2 | 32000 | 16000 | -29770.506 | 9248000 | 99.200 | 212 | 6 | 320 | 10 |
|  | 3 | 16000 | 8000 | -49463.419 | 9216000 | 98.857 | 320 | 10 | 387 | 12 |
|  | 4 | 8000 | 4000 | -76990.871 | 9136000 | 97.999 | 386 | 12 | 392 | 15 |
|  | 5 | 4000 | 2000 | -102167.502 | 8900000 | 95.467 | 392 | 15 | 330 | 23 |
|  | 6 | 2000 | 1000 | -135289.745 | 8344000 | 89.503 | 323 | 24 | 179 | 26 |
|  | 7 | 1000 | 500 | -166517.136 | 7405000 | 79.431 | 177 | 26 | 90 | 20 |
|  | 8 | 500 | 250 | -182257.417 | 6354000 | 68.157 | 91 | 18 | 43 | 11 |
|  | 9 | 250 | 125 | -15964.329 | 5434750 | 58.297 | 44 | 11 | 11 | 7 |
| 17 | 1 | 64000 | 32000 | -40315.595 | 14016000 | 99.402 | 170 | 6 | 261 | 8 |
|  | 2 | 32000 | 16000 | -50123.711 | 14016000 | 99.402 | 261 | 8 | 409 | 13 |
|  | 3 | 16000 | 8000 | -81386.265 | 13952000 | 98.948 | 409 | 13 | 579 | 16 |
|  | 4 | 8000 | 4000 | -113562.900 | 13840000 | 98.153 | 578 | 16 | 692 | 19 |
|  | 5 | 4000 | 2000 | -148739.071 | 13428000 | 95.231 | 687 | 19 | 568 | 28 |
|  | 6 | 2000 | 1000 | -184126.195 | 12648000 | 89.700 | 565 | 27 | 390 | 25 |
|  | 7 | 1000 | 500 | -223865.479 | 11378000 | 80.693 | 384 | 26 | 179 | 18 |
|  | 8 | 500 | 250 | -247551.521 | 9905000 | 70.246 | 171 | 17 | 63 | 12 |
|  | 9 | 250 | 125 | 16497.137 | 8611750 | 61.075 | 69 | 13 | 19 | 9 |
| 18 | 1 | 64000 | 32000 | -54694.590 | 16448000 | 99.684 | 179 | 7 | 279 | 11 |
|  | 2 | 32000 | 16000 | -62317.475 | 16448000 | 99.684 | 279 | 11 | 427 | 13 |
|  | 3 | 16000 | 8000 | -87807.717 | 16416000 | 99.490 | 426 | 13 | 603 | 15 |
|  | 4 | 8000 | 4000 | -133075.438 | 16312000 | 98.860 | 602 | 15 | 698 | 19 |
|  | 5 | 4000 | 2000 | -179237.541 | 15968000 | 96.775 | 699 | 19 | 574 | 27 |
|  | 6 | 2000 | 1000 | -218314.684 | 15122000 | 91.648 | 561 | 25 | 355 | 27 |
|  | 7 | 1000 | 500 | -270227.822 | 13626000 | 82.581 | 354 | 28 | 164 | 22 |
|  | 8 | 500 | 250 | -300450.566 | 11987500 | 72.651 | 163 | 21 | 69 | 20 |
|  | 9 | 250 | 125 | 20140.528 | 10526500 | 63.796 | 68 | 19 | 21 | 15 |
| 19 | 1 | 64000 | 32000 | -41591.974 | 16896000 | 99.668 | 182 | 5 | 307 | 8 |
|  | 2 | 32000 | 16000 | -55784.910 | 16832000 | 99.290 | 306 | 8 | 503 | 10 |
|  | 3 | 16000 | 8000 | -86510.716 | 16752000 | 98.818 | 502 | 10 | 679 | 14 |
|  | 4 | 8000 | 4000 | -130401.349 | 16600000 | 97.922 | 677 | 14 | 774 | 19 |
|  | 5 | 4000 | 2000 | -175196.226 | 16060000 | 94.736 | 772 | 19 | 705 | 28 |
|  | 6 | 2000 | 1000 | -208583.745 | 14962000 | 88.259 | 691 | 27 | 392 | 30 |
|  | 7 | 1000 | 500 | -257253.300 | 13183000 | 77.765 | 394 | 29 | 208 | 28 |
|  | 8 | 500 | 250 | -270933.873 | 11335000 | 66.864 | 203 | 28 | 78 | 18 |
|  | 9 | 250 | 125 | 33384.040 | 9851000 | 58.110 | 85 | 18 | 16 | 9 |
| 20 | 1 | 64000 | 32000 | -45931.268 | 14464000 | 99.977 | 157 | 5 | 279 | 9 |
|  | 2 | 32000 | 16000 | -56609.539 | 14432000 | 99.756 | 278 | 9 | 431 | 14 |
|  | 3 | 16000 | 8000 | -88131.051 | 14352000 | 99.203 | 430 | 14 | 561 | 16 |
|  | 4 | 8000 | 4000 | -123699.942 | 14248000 | 98.484 | 559 | 16 | 672 | 24 |
|  | 5 | 4000 | 2000 | -163357.084 | 14004000 | 96.798 | 670 | 24 | 585 | 27 |
|  | 6 | 2000 | 1000 | -199800.436 | 13356000 | 92.319 | 585 | 26 | 352 | 24 |
|  | 7 | 1000 | 500 | -241940.680 | 12141000 | 83.920 | 345 | 23 | 174 | 26 |
|  | 8 | 500 | 250 | -254045.172 | 10687000 | 73.870 | 179 | 26 | 86 | 18 |
|  | 9 | 250 | 125 | 24165.410 | 9388000 | 64.891 | 89 | 18 | 24 | 11 |
| 21 | 1 | 64000 | 32000 | -34245.840 | 13696000 | 99.773 | 155 | 2 | 248 | 3 |
|  | 2 | 32000 | 16000 | -43223.131 | 13632000 | 99.307 | 247 | 3 | 393 | 7 |
|  | 3 | 16000 | 8000 | -59955.957 | 13392000 | 97.558 | 390 | 7 | 573 | 5 |
|  | 4 | 8000 | 4000 | -84023.952 | 12848000 | 93.595 | 570 | 5 | 655 | 14 |
|  | 5 | 4000 | 2000 | -104084.816 | 12204000 | 88.904 | 646 | 13 | 563 | 14 |
|  | 6 | 2000 | 1000 | -119577.441 | 11084000 | 80.745 | 552 | 12 | 288 | 16 |
|  | 7 | 1000 | 500 | -152118.614 | 9811000 | 71.471 | 289 | 15 | 135 | 12 |
|  | 8 | 500 | 250 | -143163.041 | 8570500 | 62.435 | 137 | 11 | 53 | 12 |
|  | 9 | 250 | 125 | 68503.023 | 7515500 | 54.749 | 55 | 12 | 10 | 7 |

**Table S6.** Distribution of the ten most common topologies from *erato*-*sara* *Heliconius* butterflies’ genomes based on stepwise non-overlapping windows using all window trees. Top = tree topologies based on Edelman et al. (2019) and Figure 3 on the main manuscript; best = best window sizes from Table S4. Numbers in bracket refer to the proportion of windows that recover each topology compared to the total number of windows for every window size.

| **top** | **window sizes (kb)** | | | | | | | | | | |
| --- | --- | --- | --- | --- | --- | --- | --- | --- | --- | --- | --- |
|  | **64** | **32** | **16** | **8** | **4** | **2** | **1** | **0.5** | **0.25** | **0.125** | **best** |
| T1 | 1507 (37.2) | 2752 (34.0) | 4860 (30.1) | 8247 (25.7) | 13278 (20.9) | 19740 (16.0) | 26176 (11.3) | 31687 (7.6) | 35177 (4.9) | 38195 (3.1) | 35773 (4.0) |
| T2 | 1862 (46.0) | 3237 (40.0) | 5438 (33.7) | 8971 (27.9) | 14324 (22.5) | 21611 (17.5) | 28583 (12.3) | 33966 (8.1) | 37409 (5.2) | 38890 (3.1) | 38313 (4.3) |
| T3 | 444 (11.0) | 1110 (13.7) | 2481 (15.4) | 5116 (15.9) | 9818 (15.5) | 15960 (12.9) | 22779 (9.8) | 27598 (6.6) | 30200 (4.2) | 31446 (2.5) | 30844 (3.5) |
| T4 | 80 (2.0) | 301 (3.7) | 865 (5.4) | 2184 (6.8) | 4841 (7.6) | 9388 (7.6) | 15687 (6.7) | 22483 (5.4) | 28209 (3.9) | 33834 (2.7) | 29939 (3.4) |
| T5 | 42 (1.0) | 168 (2.1) | 516 (3.2) | 1264 (3.9) | 2631 (4.1) | 4644 (3.8) | 6834 (2.9) | 8879 (2.1) | 10787 (1.5) | 12522 (1.0) | 11324 (1.3) |
| T6 | 35 (0.9) | 134 (1.7) | 447 (2.8) | 1063 (3.3) | 2325 (3.7) | 4273 (3.5) | 6942 (3.0) | 10307 (2.5) | 14372 (2.0) | 18915 (1.5) | 15779 (1.8) |
| T7 | 42 (1.0) | 195 (2.4) | 674 (4.2) | 1967 (6.1) | 4713 (7.4) | 9829 (7.9) | 17374 (7.5) | 25802 (6.2) | 32935 (4.6) | 37832 (3.0) | 34506 (3.9) |
| T8 | 21 (0.5) | 104 (1.3) | 309 (1.9) | 816 (2.5) | 1975 (3.1) | 3912 (3.2) | 6627 (2.9) | 10006 (2.4) | 14072 (1.9) | 18569 (1.5) | 15439 (1.7) |
| T9 | 5 (0.1) | 19 (0.2) | 184 (1.1) | 786 (2.4) | 2396 (3.8) | 5550 (4.5) | 10245 (4.4) | 14911 (3.6) | 19009 (2.6) | 22008 (1.8) | 20049 (2.3) |
| T10 | 8 (0.2) | 52 (0.6) | 201 (1.2) | 650 (2.0) | 1593 (2.5) | 3452 (2.8) | 6270 (2.7) | 10448 (2.5) | 15307 (2.1) | 20268 (1.6) | 16795 (1.9) |
| Other | 4 (0.1) | 28 (0.3) | 185 (1.1) | 1074 (3.3) | 5644 (8.9) | 25289 (20.5) | 84917 (36.5) | 220807 (53.0) | 484741 (67.1) | 978235 (78.2) | 641281 (72.1) |
| Total | 4050 | 8100 | 16160 | 32138 | 63538 | 123648 | 232434 | 416894 | 722218 | 1250714 | 890042 |

**Table S7.** Distribution of the ten most common topologies from *erato*-*sara* *Heliconius* butterflies’ genomes based on stepwise non-overlapping windows (Figure 3 on the main manuscript) using window trees with ≥95 average UFBoot support. Top = tree topologies based on Edelman et al. (2019) and Figure 3 on the main manuscript; best = best window sizes from Table S4. Numbers in bracket refer to the proportion of windows that recover each topology compared to the total number of windows for every window size.

| **top** | **window sizes (kb)** | | | | | | | | | | |
| --- | --- | --- | --- | --- | --- | --- | --- | --- | --- | --- | --- |
|  | **64** | **32** | **16** | **8** | **4** | **2** | **1** | **0.5** | **0.25** | **0.125** | **best** |
| T1 | 1302 (42.7) | 2117 (41.9) | 3160 (40.0) | 4043 (36.7) | 3954 (31.8) | 2780 (27.2) | 1410 (23.6) | 615 (21.3) | 218 (18.6) | 55 (15.9) | 153 (16.4) |
| T2 | 1406 (46.1) | 2066 (40.9) | 2796 (35.4) | 3428 (31.1) | 3365 (27.1) | 2359 (23.1) | 1092 (18.3) | 420 (14.5) | 159 (13.5) | 52 (15.0) | 141 (15.1) |
| T3 | 249 (8.2) | 549 (10.9) | 1121 (14.2) | 1880 (17.1) | 2499 (20.1) | 2263 (22.1) | 1383 (23.1) | 586 (20.3) | 214 (18.2) | 56 (16.1) | 185 (19.8) |
| T4 | 38 (1.2) | 123 (2.4) | 283 (3.6) | 533 (4.8) | 795  (6.4) | 855 (8.4) | 568 (9.5) | 313 (10.8) | 120 (10.2) | 38 (11.0) | 86 (9.2) |
| T5 | 21 (0.7) | 61 (1.2) | 140 (1.8) | 277 (2.5) | 323 (2.6) | 281 (2.7) | 161 (2.7) | 85 (2.9) | 28 (2.4) | 5 (1.4) | 22  (2.4) |
| T6 | 13 (0.4) | 41 (0.8) | 104 (1.3) | 158 (1.4) | 210 (1.7) | 147 (1.4) | 85 (1.4) | 41 (1.4) | 11 (0.9) | 2 (0.6) | 10 (1.1) |
| T7 | 6 (0.2) | 36 (0.7) | 136 (1.7) | 317 (2.9) | 584 (4.7) | 616 (6.0) | 458 (7.7) | 246 (8.5) | 101 (8.6) | 27 (7.8) | 90 (9.6) |
| T8 | 11 (0.4) | 39 (0.8) | 92 (1.2) | 138 (1.3) | 151 (1.2) | 138 (1.3) | 70 (1.2) | 43 (1.5) | 18 (1.5) | 8 (2.3) | 14 (1.5) |
| T9 | 1 (<0.1) | 5 (0.1) | 21 (0.3) | 112 (1.0) | 257 (2.1) | 353 (3.5) | 340 (5.7) | 212 (7.3) | 114 (9.7) | 29 (8.4) | 81 (8.7) |
| T10 | 1 (<0.1) | 9 (0.2) | 22 (0.3) | 59 (0.5) | 94 (0.8) | 88 (0.9) | 65 (1.1) | 44 (1.5) | 21 (1.8) | 8 (2.3) | 20 (2.1) |
| Other | 1 (<0.1) | 7 (<0.1) | 25 (0.1) | 77 (0.3) | 192 (0.9) | 343 (2.6) | 344 (5.6) | 282 (9.1) | 170 (15.6) | 67 (20.0) | 132 (13.3) |
| Total | 3049 | 5053 | 7900 | 11022 | 12424 | 10223 | 5976 | 2887 | 1174 | 347 | 934 |

**Table S8.** Best window size for each great apes’ chromosome based on stepwise non-overlapping windows. Length is in the units of base pairs. Initial length = length of chromosome before filtering; 64kb length = length of chromosome after divided into 64kb non-overlapping windows; final length = length of chromosome when the best window size is found; best window size = window size with the best AIC score. Percentage of final length refers to the initial length of the alignment. *Mitochondrial genome starts with 16kb instead of 64kb window size.

| **chr** | **Initial length** | **64kb length** | **final length** | **best window size** |
| --- | --- | --- | --- | --- |
| 1 | 227,082,584 | 227,072,000 | 192,610,000 (84.82%) | 1,000 |
| 2 | 240,480,912 | 240,448,000 | 208,534,000 (86.72%) | 1,000 |
| 3 | 194,975,254 | 194,944,000 | 171,902,000 (88.17%) | 1,000 |
| 4 | 187,599,630 | 187,584,000 | 163,660,000 (87.24%) | 1,000 |
| 5 | 181,263,364 | 181,248,000 | 153,328,000 (84.59%) | 1,000 |
| 6 | 170,078,522 | 170,048,000 | 147,120,000 (86.50%) | 1,000 |
| 7 | 158,966,559 | 158,912,000 | 126,537,000 (79.60%) | 500 |
| 8 | 144,766,429 | 144,704,000 | 123,918,000 (85.60%) | 1,000 |
| 9 | 119,357,962 | 119,296,000 | 97,018,000 (81.28%) | 1,000 |
| 10 | 133,258,722 | 133,248,000 | 108,989,000 (81.79%) | 500 |
| 11 | 134,426,162 | 134,400,000 | 110,742,000 (82.38%) | 1,000 |
| 12 | 129,897,401 | 129,856,000 | 113,700,000 (87.53%) | 1,000 |
| 13 | 95,930,070 | 95,872,000 | 83,011,000 (86.53%) | 500 |
| 14 | 88,310,862 | 88,256,000 | 75,576,000 (85.58%) | 1,000 |
| 15 | 81,856,706 | 81,856,000 | 66,643,000 (81.41%) | 500 |
| 16 | 81,730,111 | 81,728,000 | 62,160,000 (76.06%) | 500 |
| 17 | 82,919,899 | 82,880,000 | 63,168,000 (76.18%) | 500 |
| 18 | 74,650,691 | 74,624,000 | 64,954,000 (87.01%) | 500 |
| 19 | 55,192,149 | 55,168,000 | 39,050,000 (70.75%) | 500 |
| 20 | 60,890,884 | 60,864,000 | 51,026,000 (83.80%) | 500 |
| 21 | 37,406,299 | 37,376,000 | 28,655,000 (76.61%) | 500 |
| 22 | 39,008,215 | 38,976,000 | 26,745,000 (68.56%) | 500 |
| X | 154,858,262 | 154,816,000 | 112,756,000 (72.81%) | 1,000 |
| Y | 25,917,497 | 25,856,000 | 581,000 (2.24%) | 500 |
| M* | 16,568 | 16,000 | 16,000 (96.57%) | 4,000 |

**Table S9.** Distribution of window topologies for each great apes’ chromosome based on all window trees (left) and window trees with ≥95 average UFBoot support (right). Counts are based on the best window sizes from stepwise non-overlapping windows (Table S8). H: human; O: orangutan; G: gorilla; C: chimpanzee.

| **chr** | **all window trees** | | | **window trees ≥95 UFBoot support** | | |
| --- | --- | --- | --- | --- | --- | --- |
|  | **[H,C\|O,G]** | **[H,G\|O,C]** | **[H,O\|C,G]** | **[H,C\|O,G]** | **[H,G\|O,C]** | **[H,O\|C,G]** |
| 1 | 124022 | 34425 | 34163 | 38093 | 2614 | 2366 |
| 2 | 133350 | 37657 | 37527 | 42057 | 2936 | 2604 |
| 3 | 108178 | 31578 | 32146 | 33757 | 2580 | 2483 |
| 4 | 99458 | 32218 | 31984 | 32145 | 3020 | 3056 |
| 5 | 95741 | 28819 | 28768 | 30059 | 2209 | 2275 |
| 6 | 92044 | 27587 | 27489 | 28711 | 2183 | 2265 |
| 7 | 144555 | 55108 | 53411 | 24386 | 2269 | 1926 |
| 8 | 77561 | 23371 | 22986 | 25191 | 2097 | 1971 |
| 9 | 62993 | 17106 | 16919 | 20476 | 1191 | 1171 |
| 10 | 125897 | 46609 | 45472 | 22172 | 2011 | 1742 |
| 11 | 71639 | 19502 | 19601 | 23054 | 1647 | 1542 |
| 12 | 72321 | 20780 | 20599 | 22421 | 1641 | 1494 |
| 13 | 92711 | 37001 | 36310 | 15785 | 1682 | 1641 |
| 14 | 47861 | 14029 | 13686 | 15021 | 1175 | 956 |
| 15 | 75786 | 29102 | 28398 | 11956 | 1222 | 1010 |
| 16 | 72885 | 25931 | 25504 | 13703 | 1078 | 1000 |
| 17 | 71055 | 27638 | 27643 | 10296 | 775 | 793 |
| 18 | 72268 | 29019 | 28621 | 12188 | 1343 | 1264 |
| 19 | 42988 | 17759 | 17353 | 7220 | 981 | 765 |
| 20 | 58214 | 21931 | 21907 | 10010 | 852 | 776 |
| 21 | 31389 | 13232 | 12689 | 5903 | 998 | 758 |
| 22 | 30632 | 11659 | 11199 | 5458 | 709 | 417 |
| X | 84691 | 14250 | 13815 | 28759 | 727 | 514 |
| Y | 618 | 245 | 299 | 169 | 3 | 9 |
| M | 4 | 0 | 0 | 3 | 0 | 0 |
| Total | 1888861 (60.7%) | 616556 (19.8%) | 608489 (19.5%) | 478993 (86.8%) | 37943 (6.9%) | 34798 (6.3%) |

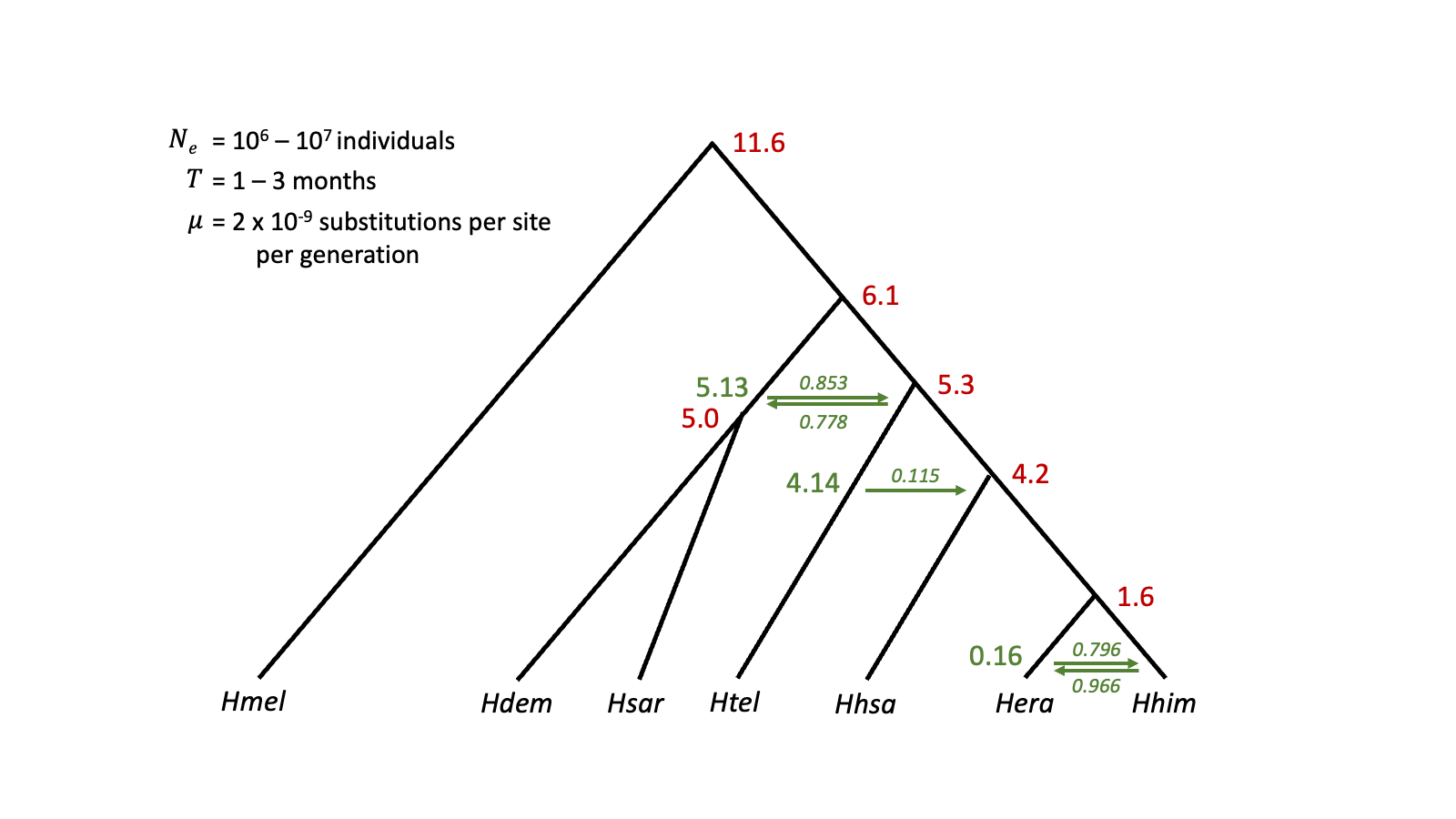
**Figure S1.** Parameters for simulating locus trees in ms. Divergence and introgression times (in Myr) are coloured red and green, respectively. The direction of introgression is shown as arrow, while the italic numbers show inheritance probability. N_e_ = effective population size, T = generation time, µ = mutation rate.

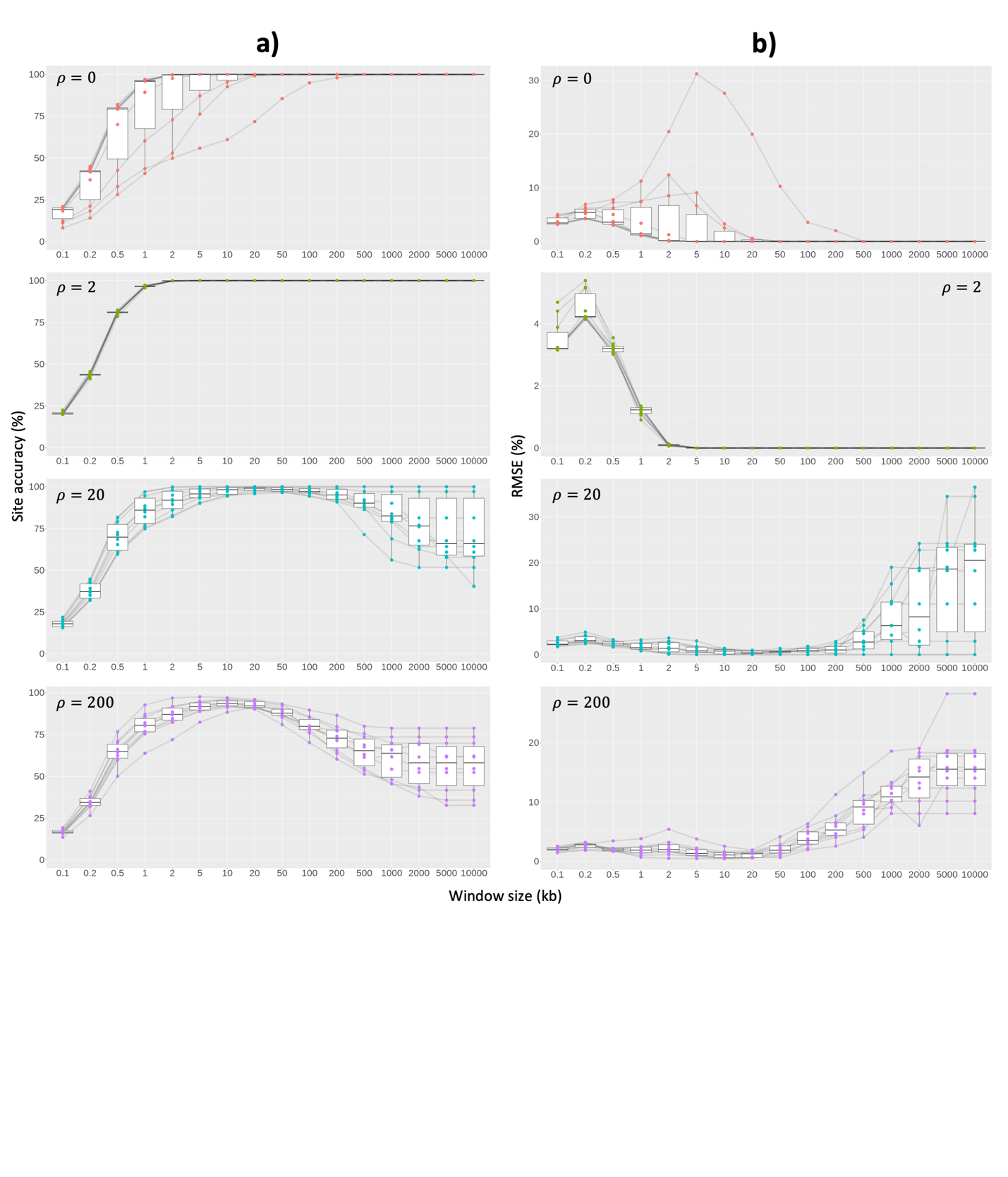

**Figure S2**. (a) Site accuracy and (b) RMSE of non-overlapping windows on simulated chromosomes with low ILS level. Each box represents different recombination rates. Each dot represents an individual result. Black lines connect results from the same simulated alignment.

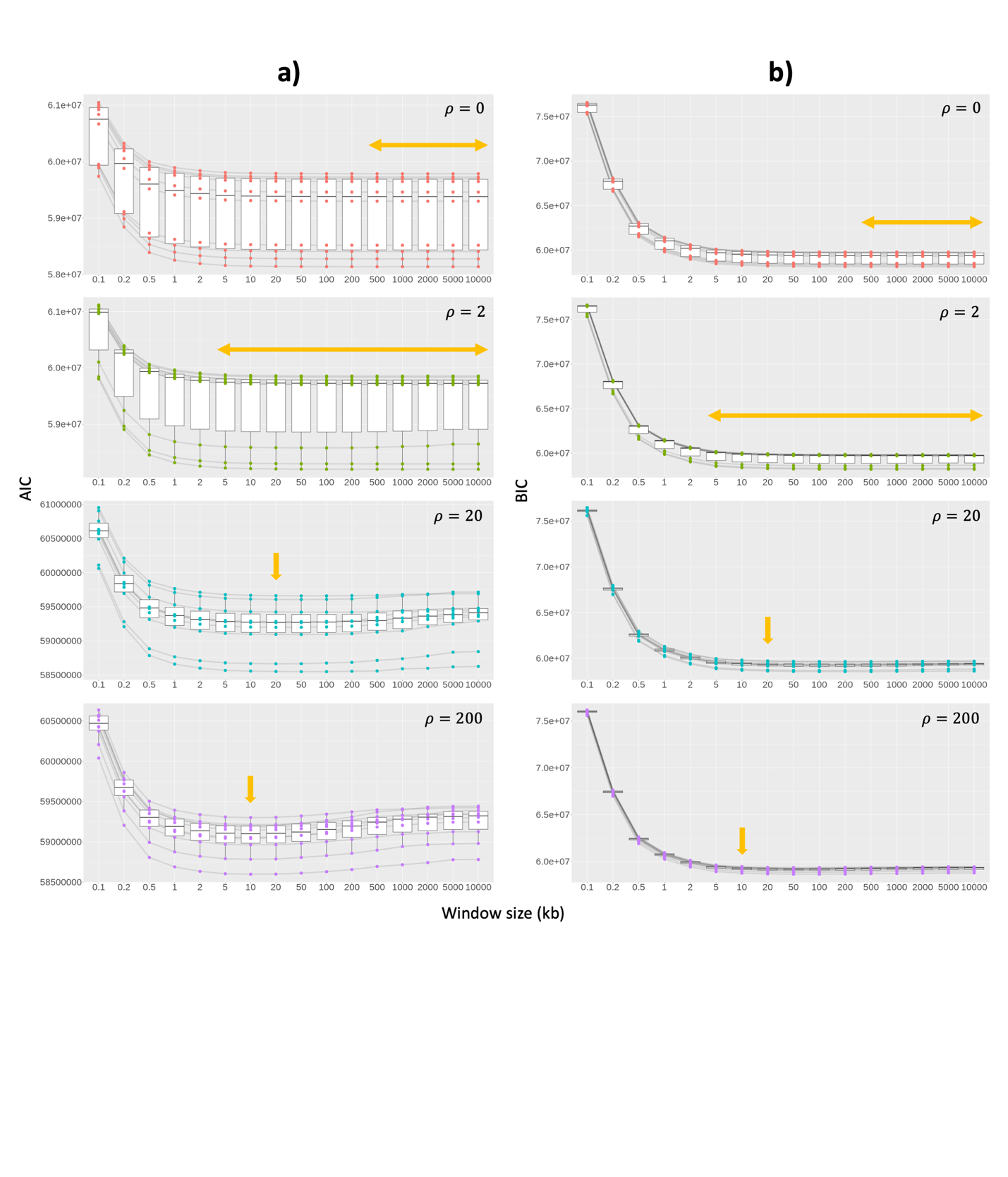

**Figure S3**. Correlation between the (a) AIC and (b) BIC with window sizes on simulated chromosomes with low ILS level. Light orange arrows show window size(s) with the highest average site accuracy from Figure S2. Each box represents different recombination rates. Each dot represents an individual result. Black lines connect results from the same simulated alignment.

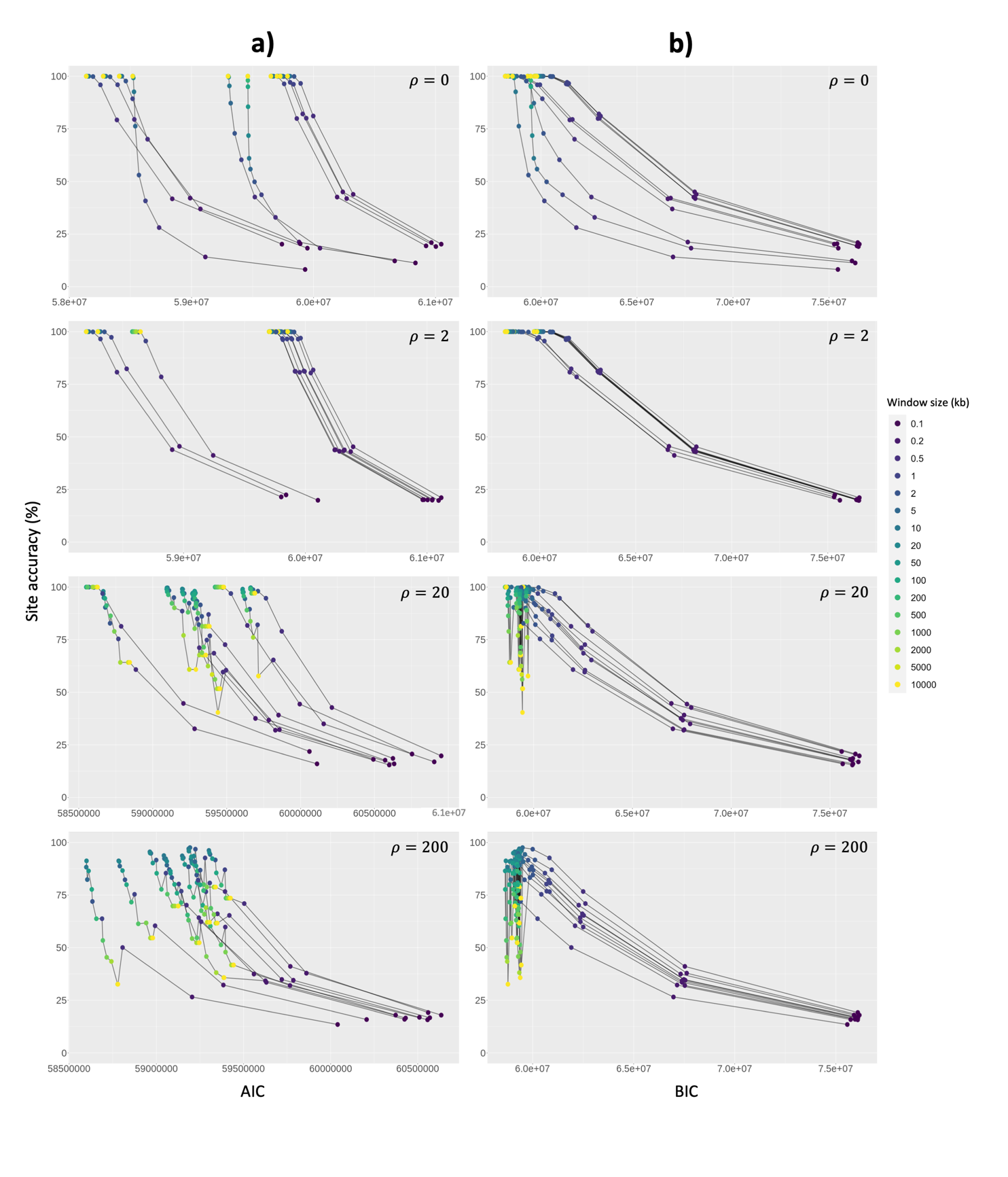

**Figure S4.** Correlation between the (a) AIC and (b) BIC with site accuracy on simulated chromosomes with low ILS level. Each box represents different recombination rates. Each dot represents an individual result and coloured by window size. Black lines connect results from the same simulated alignment.

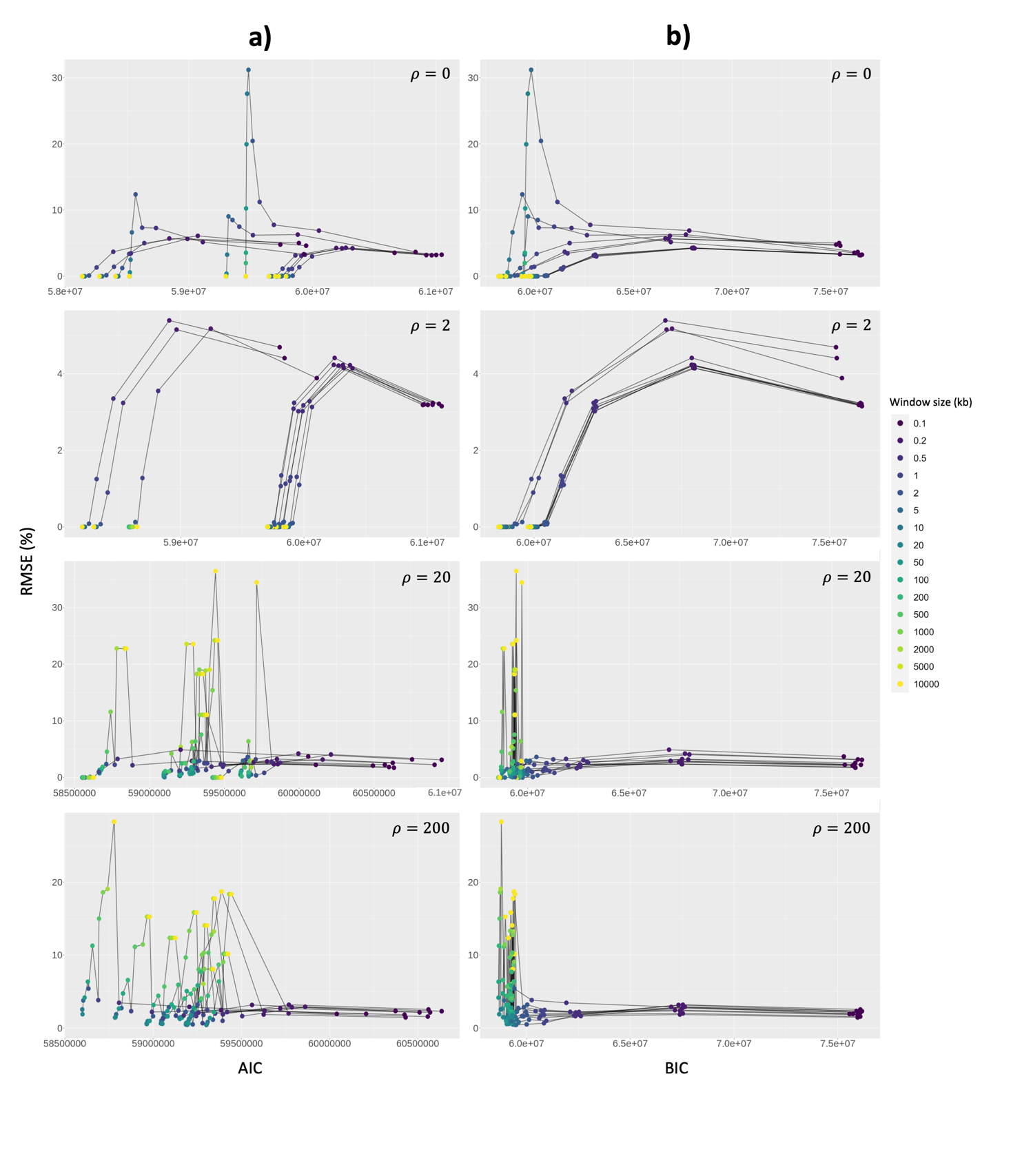

**Figure S5.** Correlation between the (a) AIC and (b) BIC with RMSE on simulated chromosomes with low ILS level. Each box represents different recombination rates. Each dot represents an individual result and coloured by window size. Black lines connect results from the same simulated alignment.

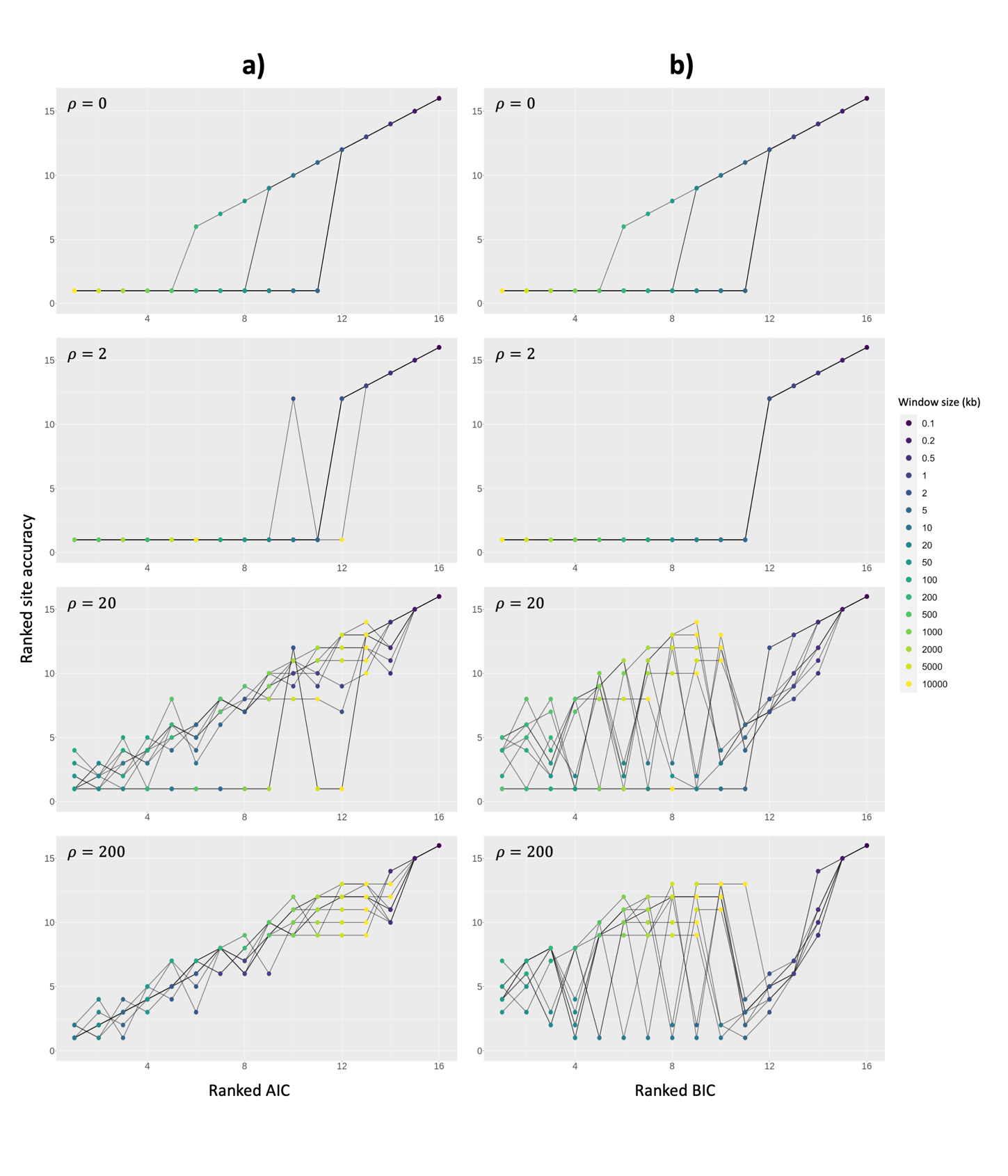

**Figure S6.** Correlation between (a) ranked AIC and (b) ranked BIC with ranked site accuracy on simulated chromosomes with low ILS level. For site accuracy, we multiplied the value by minus one, so that the highest site accuracy is ranked one. In case of a tie, the best rank was applied for the respective window sizes using min_rank() function in R. Each box represents different recombination rates. Each dot represents an individual result and coloured by window size. Black lines connect results from the same simulated alignment.

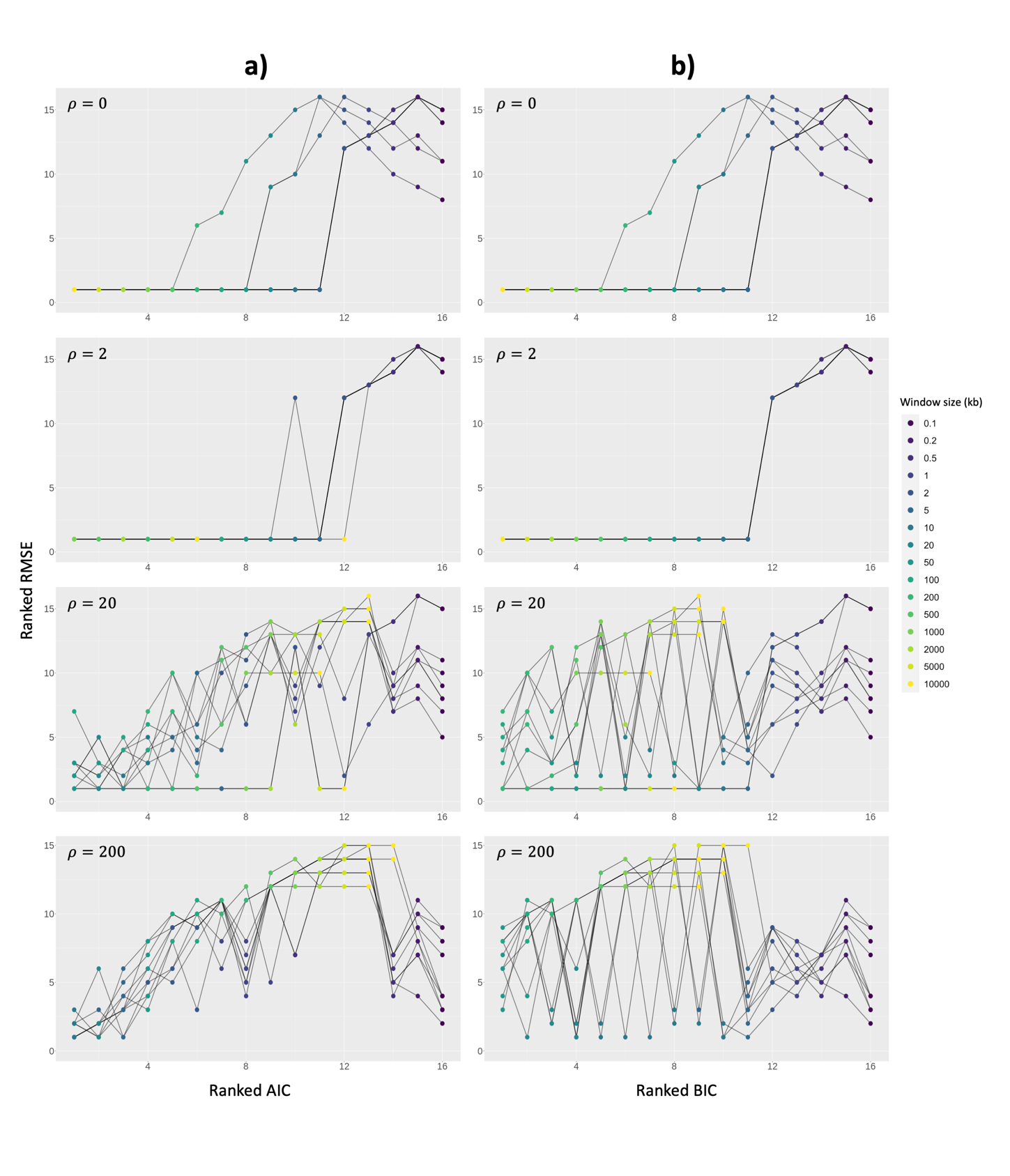

**Figure S7.** Correlation between (a) ranked AIC and (b) ranked BIC with RMSE on simulated chromosomes with low ILS level. For RMSE, the lowest RMSE is ranked one. In case of a tie, the best rank was applied for respective window sizes using min_rank() function in R. Each box represents different recombination rates. Each dot represents an individual result, connected based on replicate and coloured by window size.

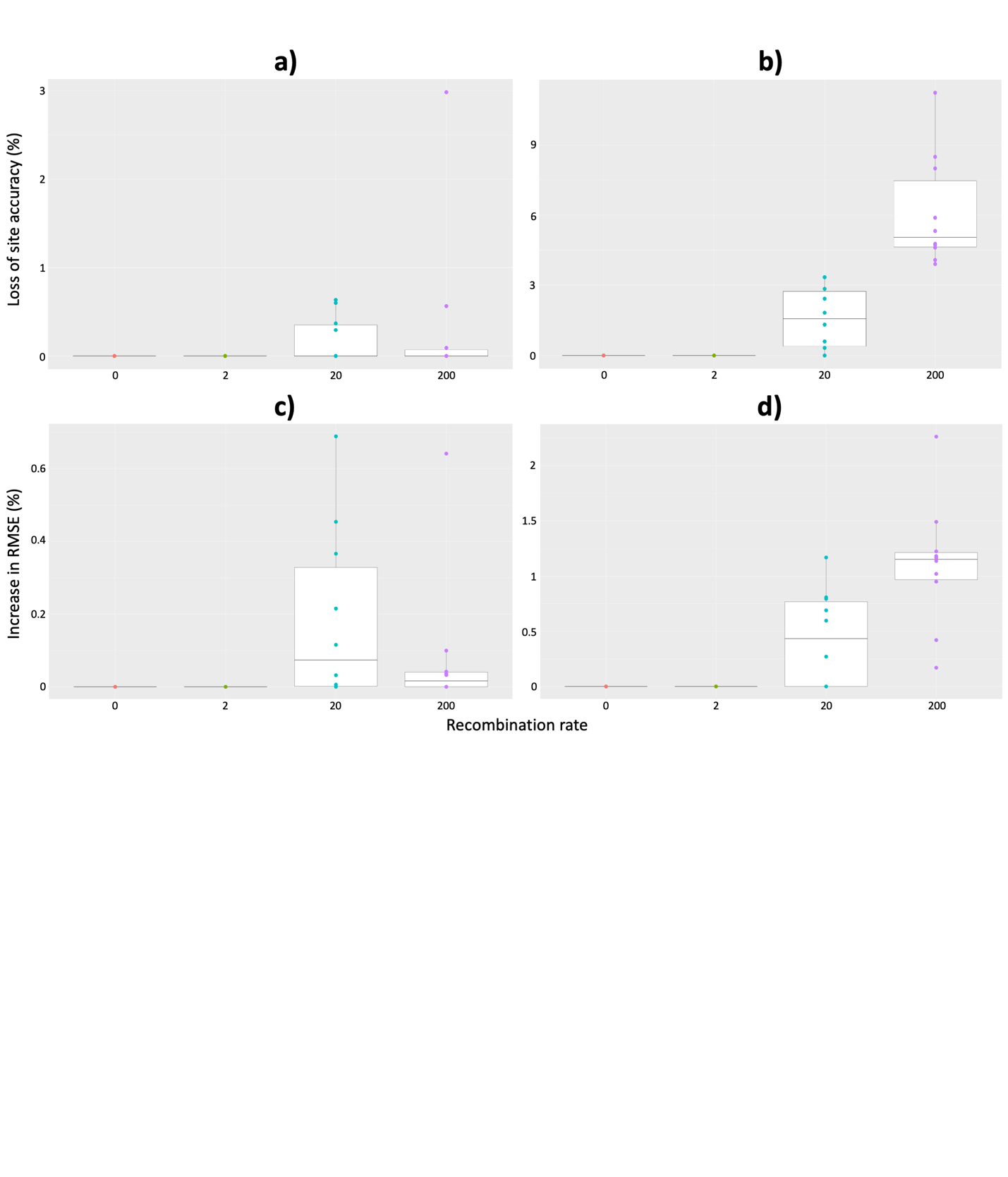

**Figure S8.** (a-b) Site accuracy loss and (c-d) RMSE gain of AIC (left) and BIC (right) across recombination rates on simulated chromosomes with low ILS level. Each dot represents an individual result (i.e., ten replicates per recombination rate).

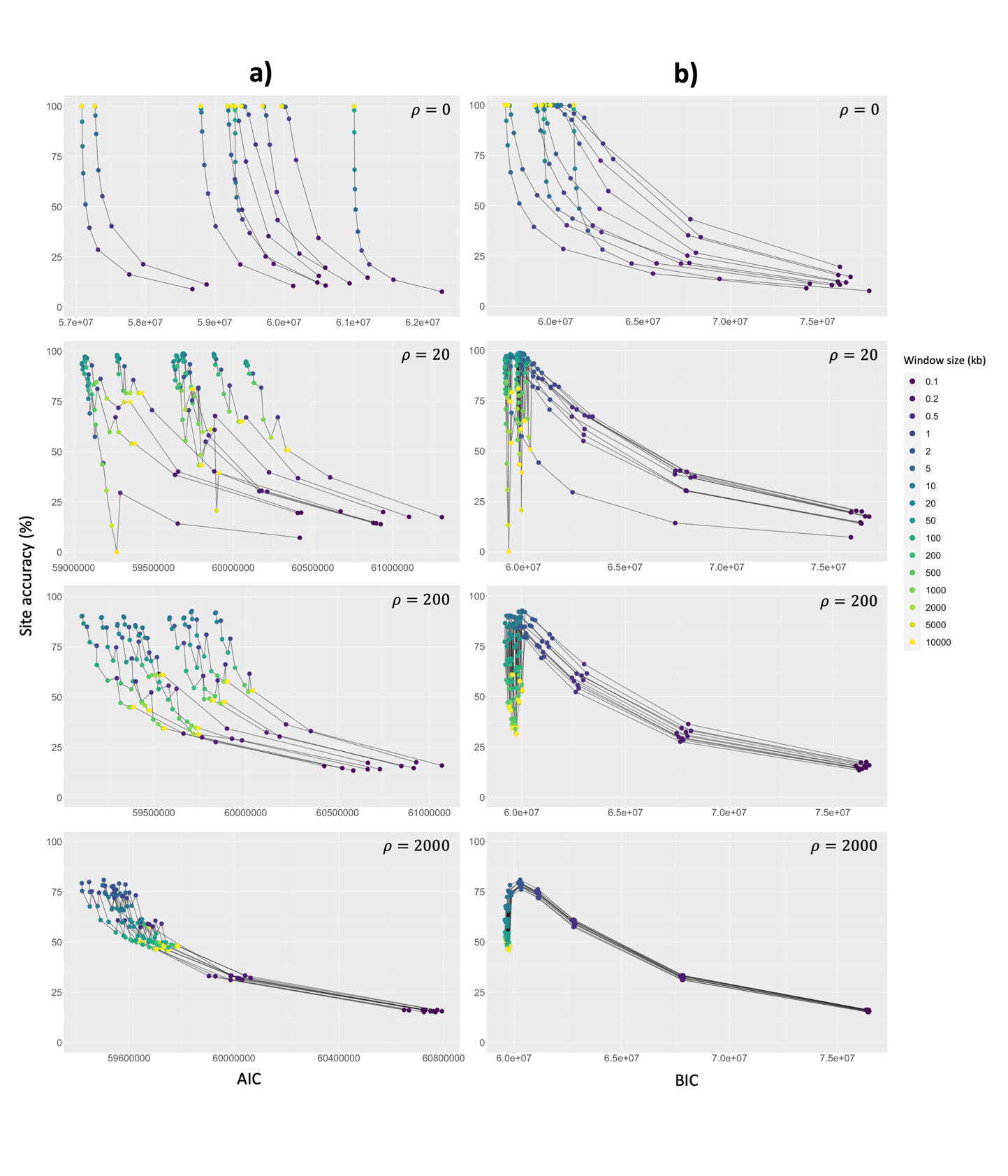

**Figure S9.** Correlation between the (a) AIC and (b) BIC with site accuracy on simulated chromosomes with medium ILS level. Each box represents different recombination rates. Each dot represents an individual result and coloured by window size. Black lines connect results from the same simulated alignment.

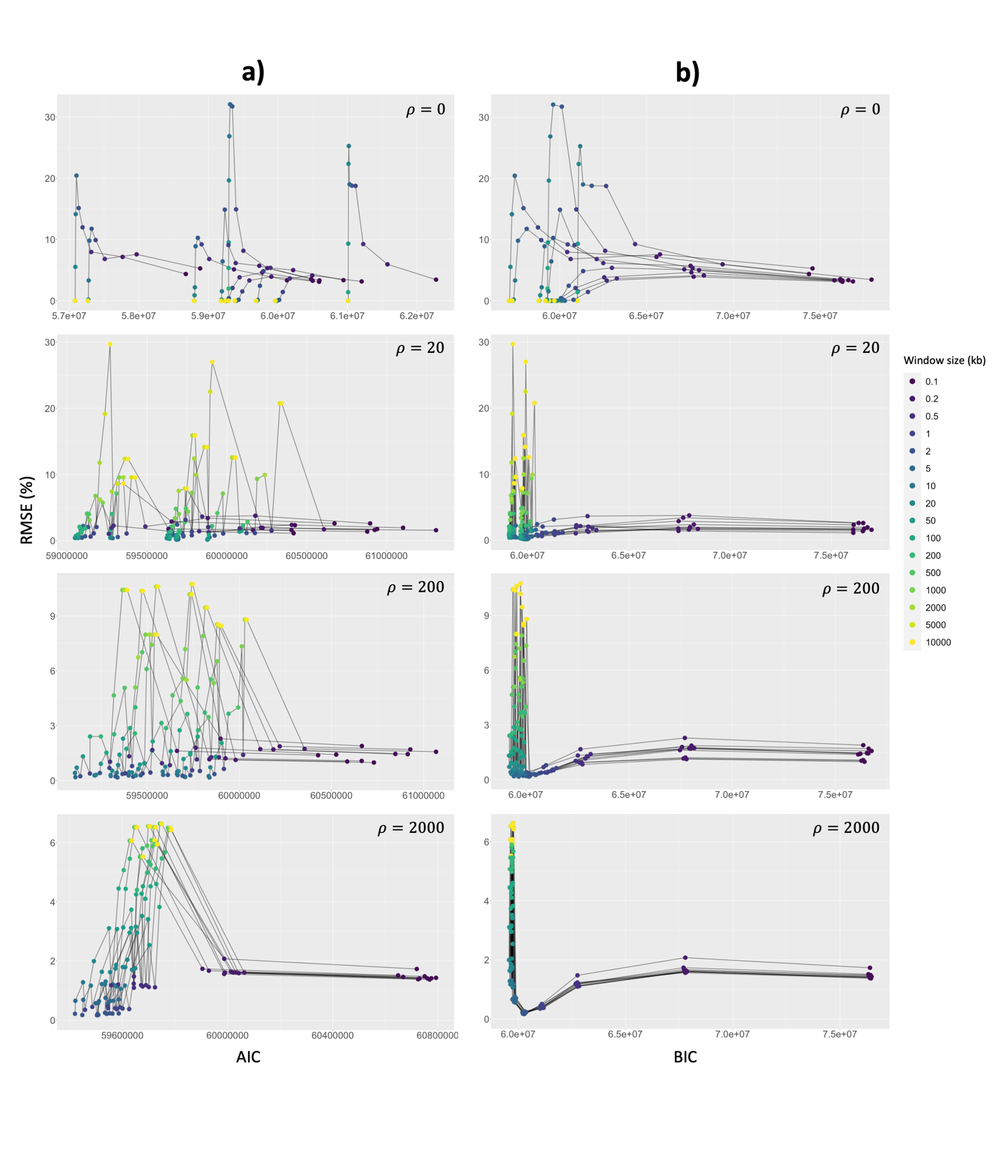

**Figure S10.** Correlation between the (a) AIC and (b) BIC with RMSE on simulated chromosomes with medium ILS level. Each box represents different recombination rates. Each dot represents an individual result and coloured by window size. Black lines connect results from the same simulated alignment.

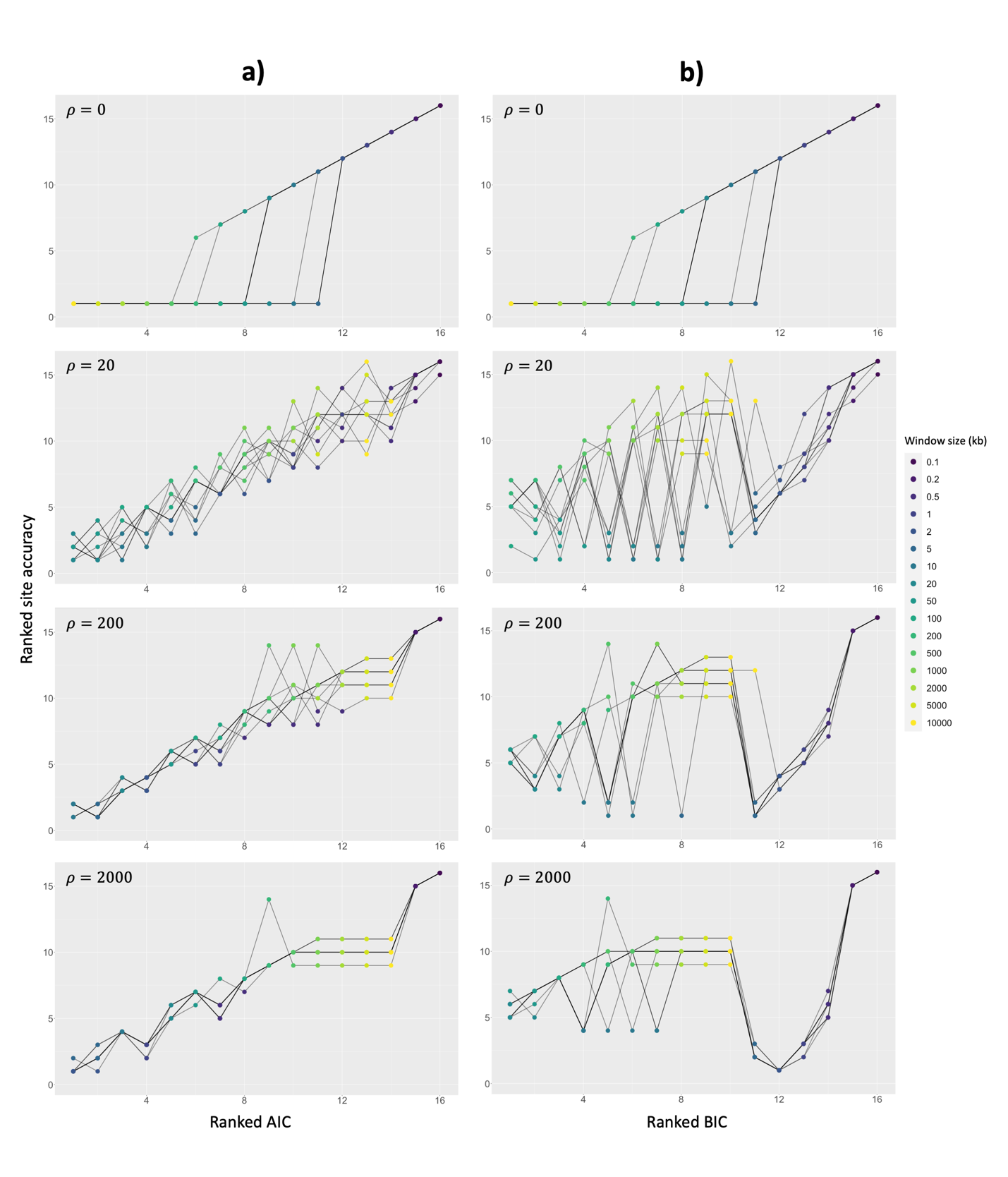

**Figure S11.** Correlation between (a) ranked AIC and (b) ranked BIC with ranked site accuracy on simulated chromosomes with medium ILS level. For site accuracy, we multiplied the value by minus one, so that the highest site accuracy is ranked one. In case of a tie, the best rank was applied for respective window sizes using min_rank() function in R. Each box represents different recombination rates. Each dot represents an individual result and coloured by window size. Black lines connect results from the same simulated alignment.

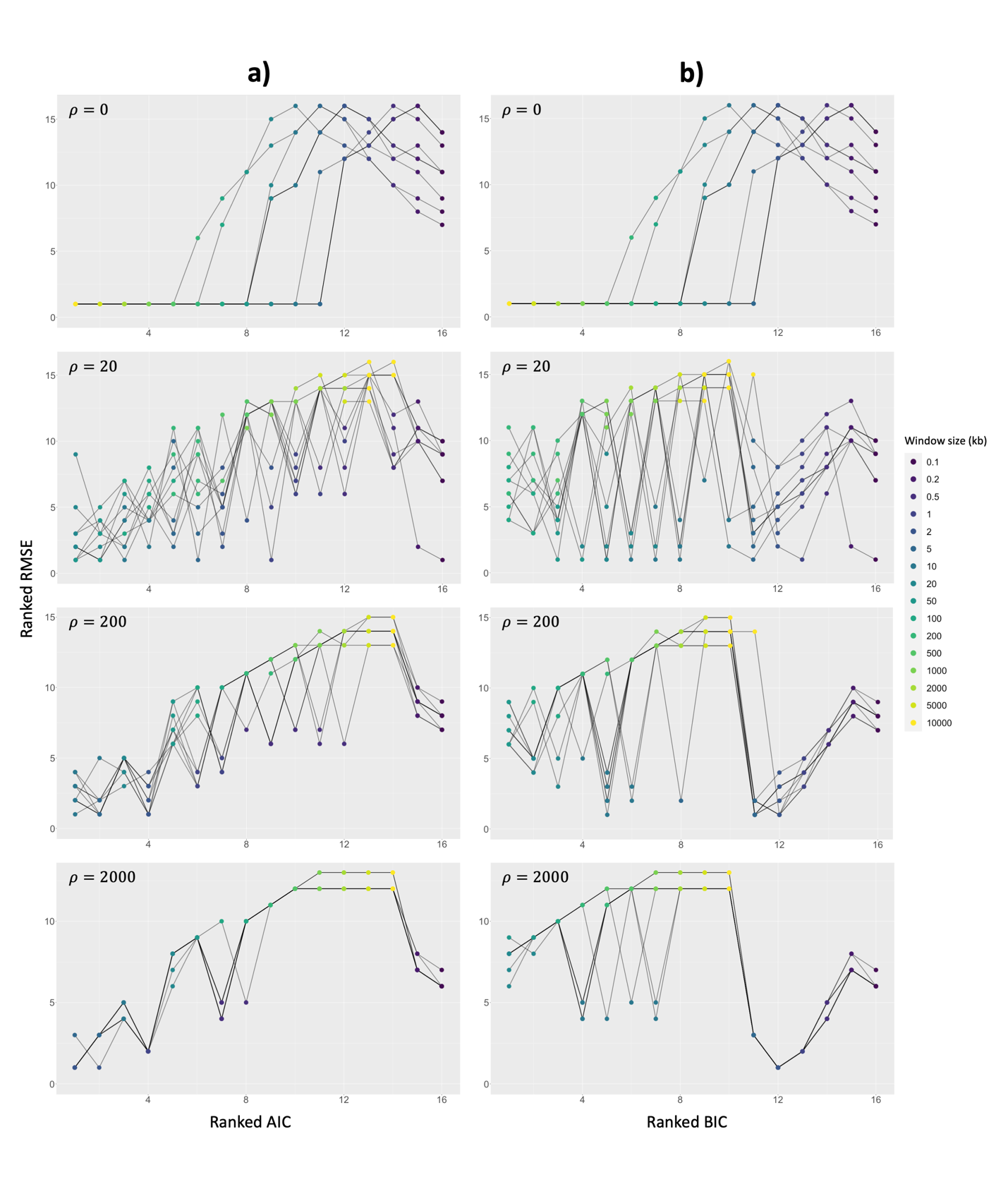

**Figure S12.** Correlation between (a) ranked AIC and (b) ranked BIC with RMSE on simulated chromosomes with medium ILS level. For RMSE, the lowest RMSE is ranked one. In case of a tie, the best rank was applied for respective window sizes using min_rank() function in R. Each box represents different recombination rates. Each dot represents an individual result and coloured by window size. Black lines connect results from the same simulated alignment.

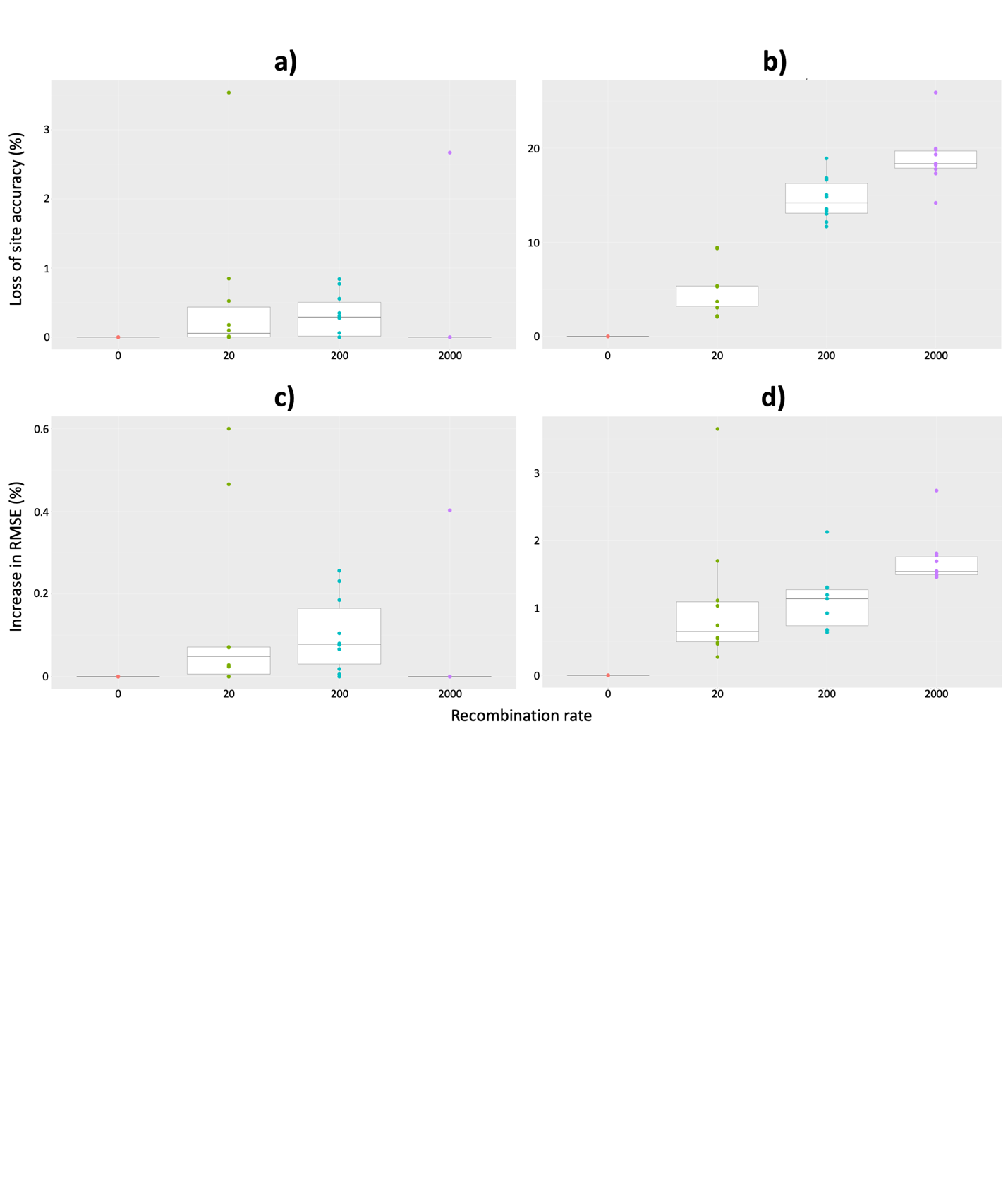

**Figure S13.** (a-b) Site accuracy loss and (c-d) RMSE gain of AIC (left) and BIC (right) across recombination rates on simulated chromosomes with medium ILS level. Each dot represents an individual result (i.e., ten replicates per recombination rate).

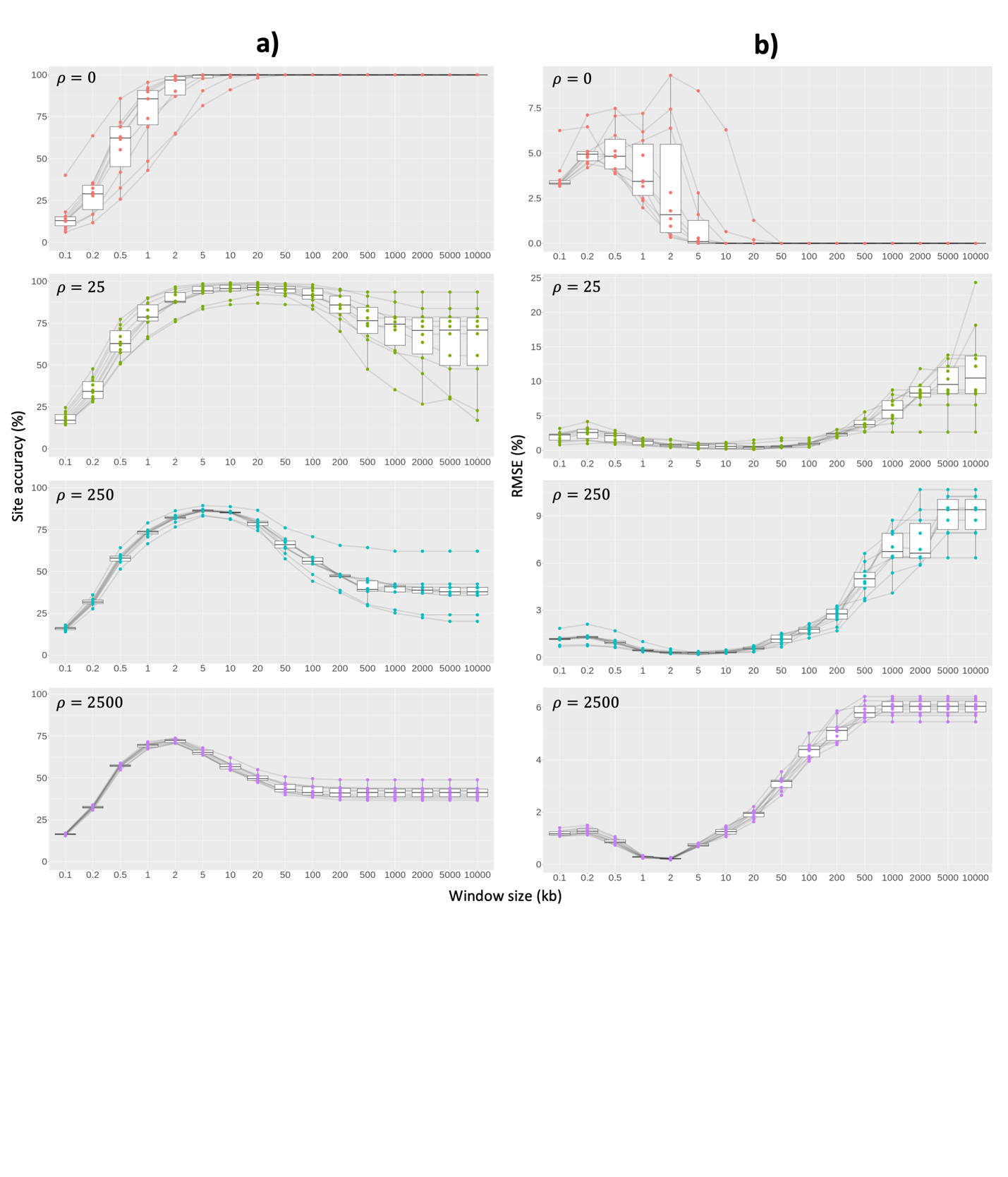

**Figure S14**. (a) Site accuracy and (b) RMSE of non-overlapping windows on simulated chromosomes with high ILS level. Each box represents different recombination rates. Each dot represents an individual result. Black lines connect results from the same simulated alignment.

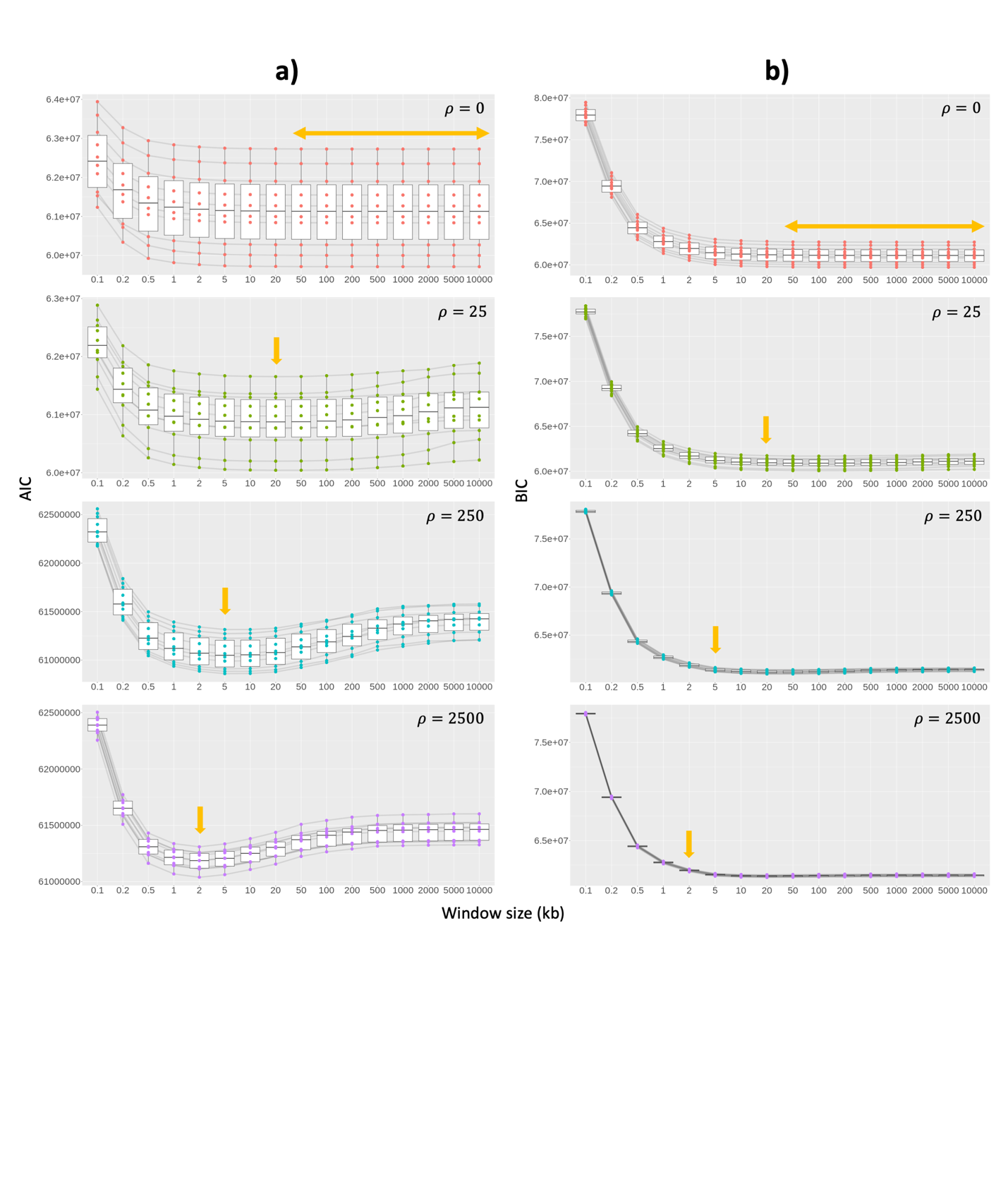

**Figure S15**. Correlation between the (a) AIC and (b) BIC with window sizes on simulated chromosomes with high ILS level. Light orange arrows show window size(s) with the highest average site accuracy from Figure S14. Each box represents different recombination rates. Each dot represents an individual result. Black lines connect results from the same simulated alignment.

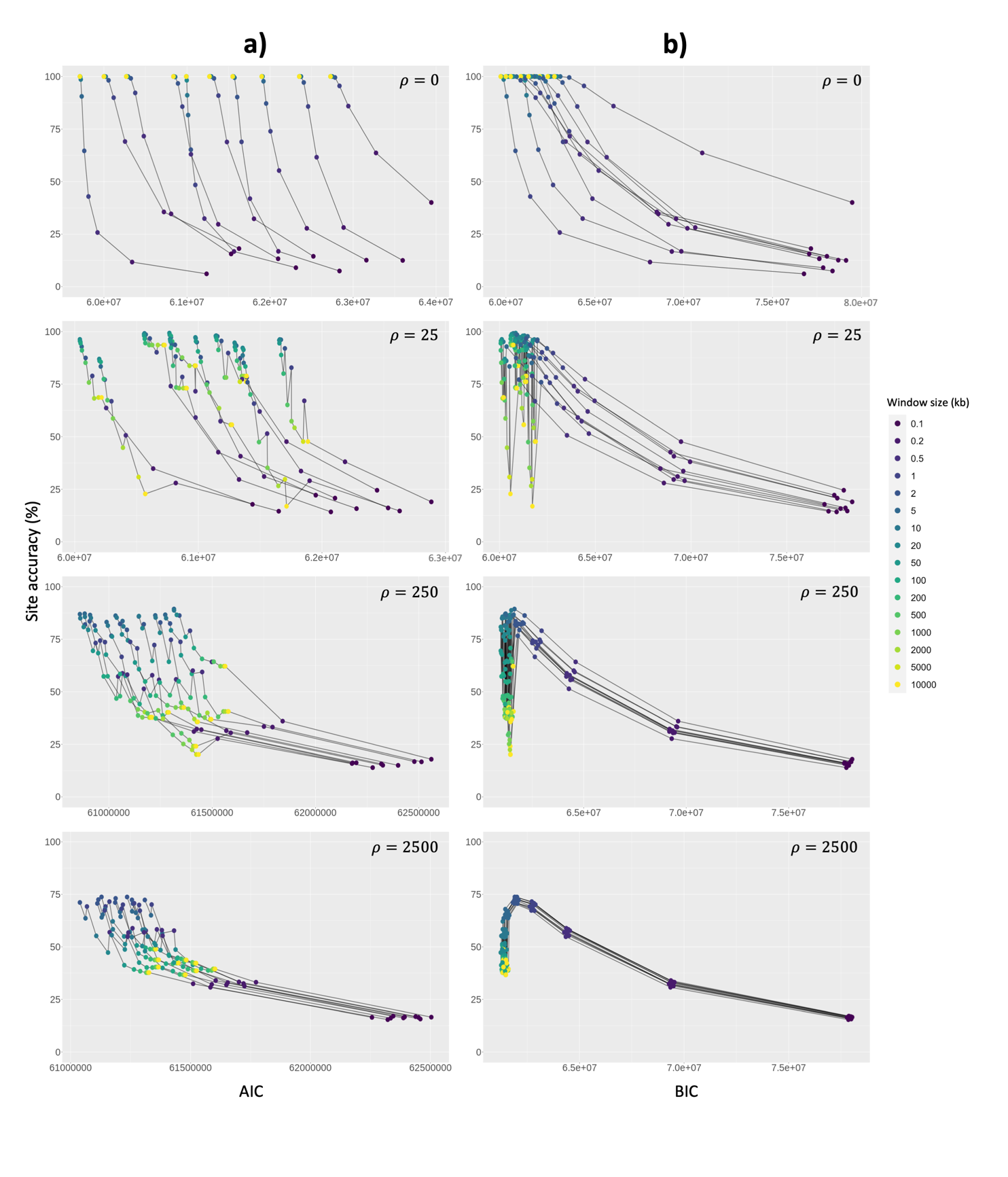

**Figure S16.** Correlation between the (a) AIC and (b) BIC with site accuracy on simulated chromosomes with high ILS level. Each box represents different recombination rates. Each dot represents an individual result and coloured by window size. Black lines connect results from the same simulated alignment.

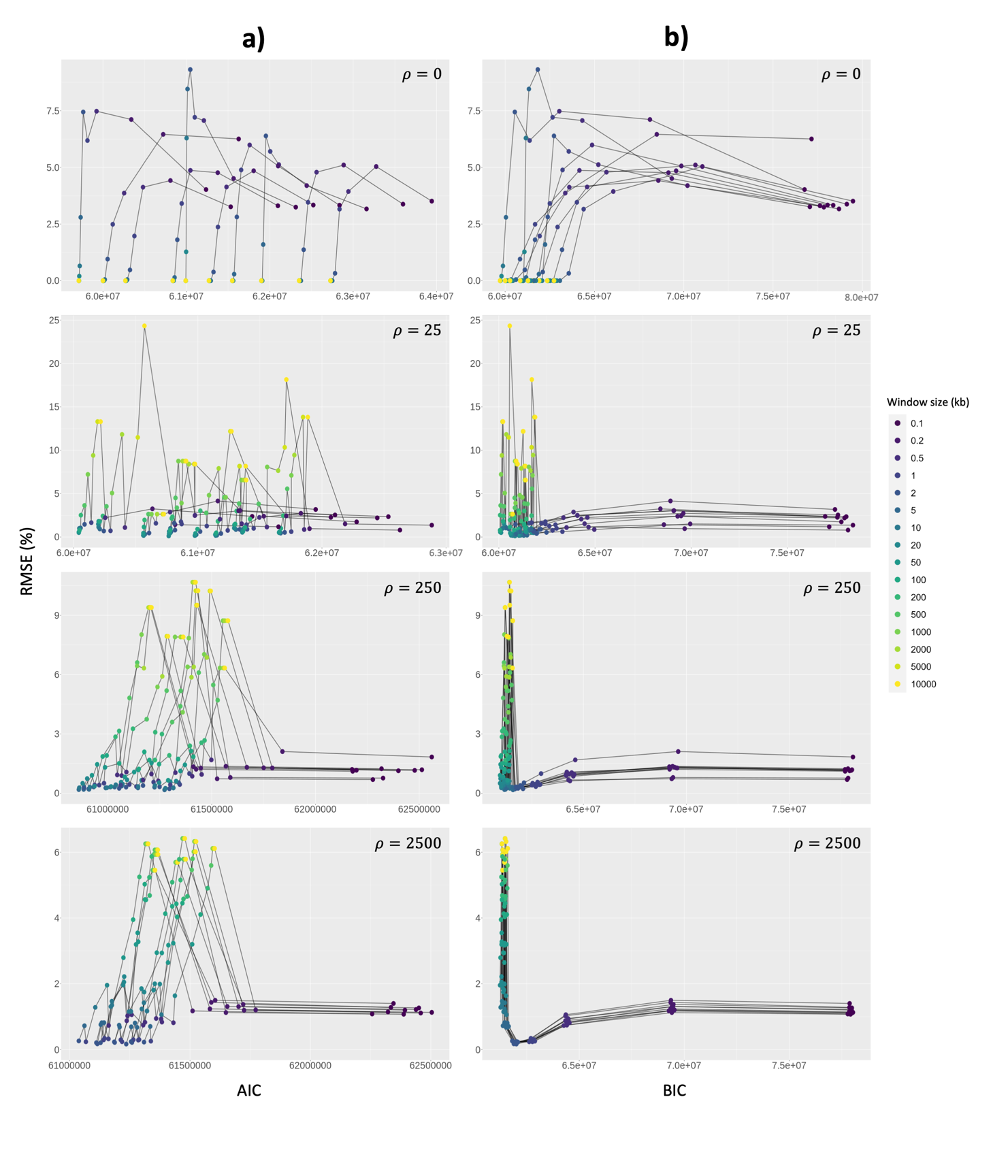

**Figure S17.** Correlation between the (a) AIC and (b) BIC with RMSE on simulated chromosomes with high ILS level. Each box represents different recombination rates. Each dot represents an individual result and coloured by window size. Black lines connect results from the same simulated alignment.

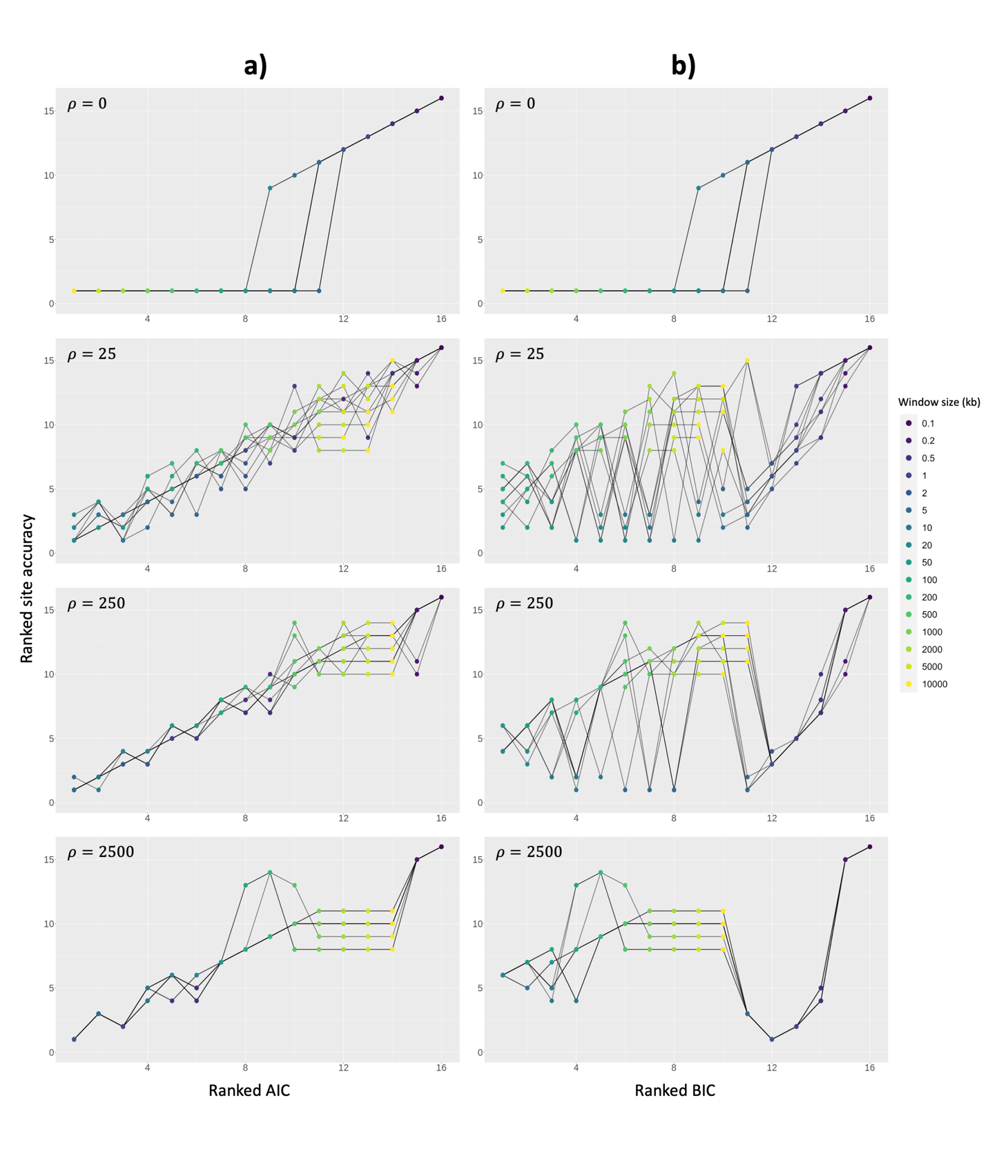

**Figure S18.** Correlation between (a) ranked AIC and (b) ranked BIC with ranked site accuracy on simulated chromosomes with high ILS level. For site accuracy, we multiplied the value by minus one, so that the highest site accuracy is ranked one. In case of a tie, the best rank was applied for respective window sizes using min_rank() function in R. Each box represents different recombination rates. Each dot represents an individual result and coloured by window size. Black lines connect results from the same simulated alignment.

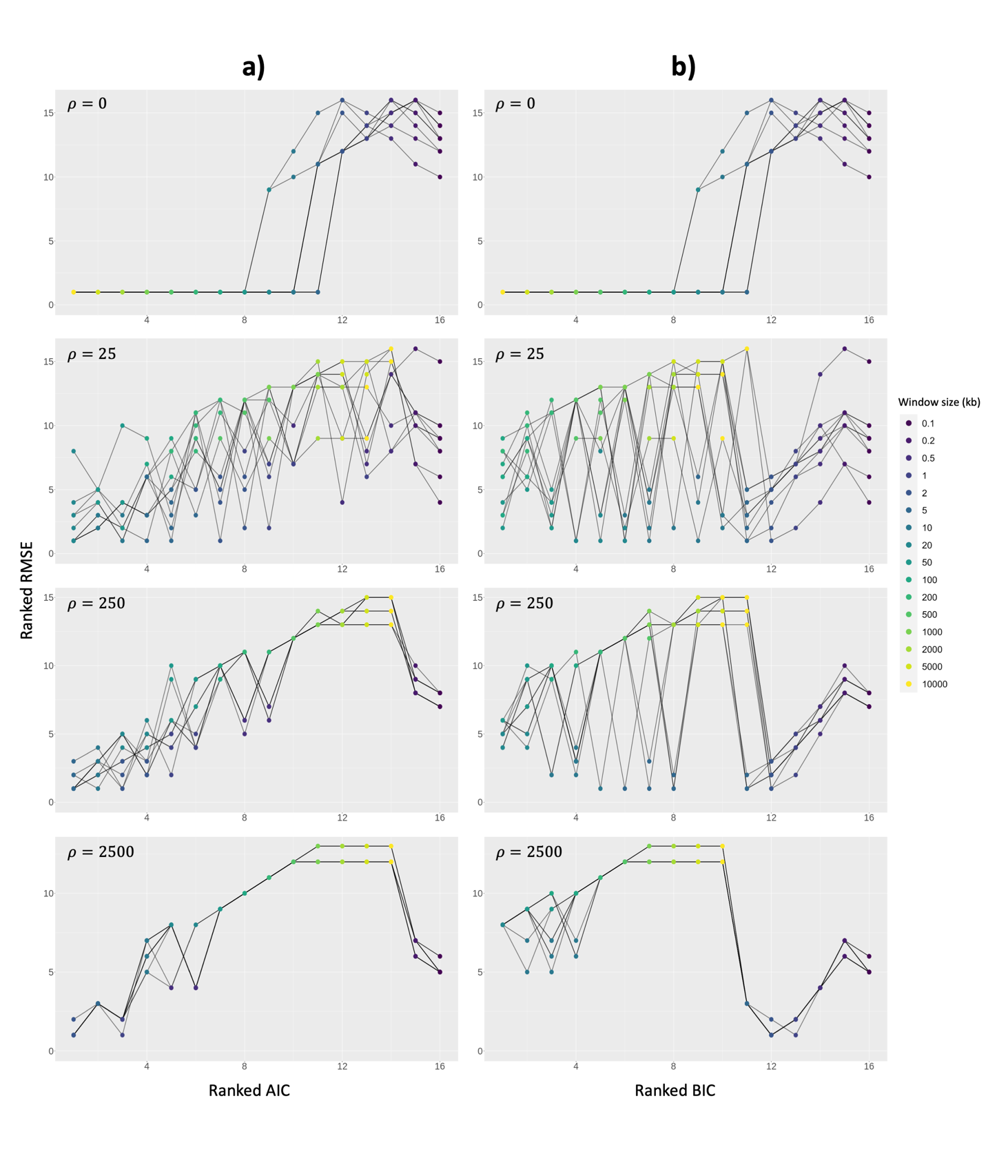

**Figure S19.** Correlation between (a) ranked AIC and (b) ranked BIC with RMSE on simulated chromosomes with high ILS level. For RMSE, the lowest RMSE is ranked one. In case of a tie, the best rank was applied for respective window sizes using min_rank() function in R. Each box represents different recombination rates. Each dot represents an individual result and coloured by window size. Black lines connect results from the same simulated alignment.

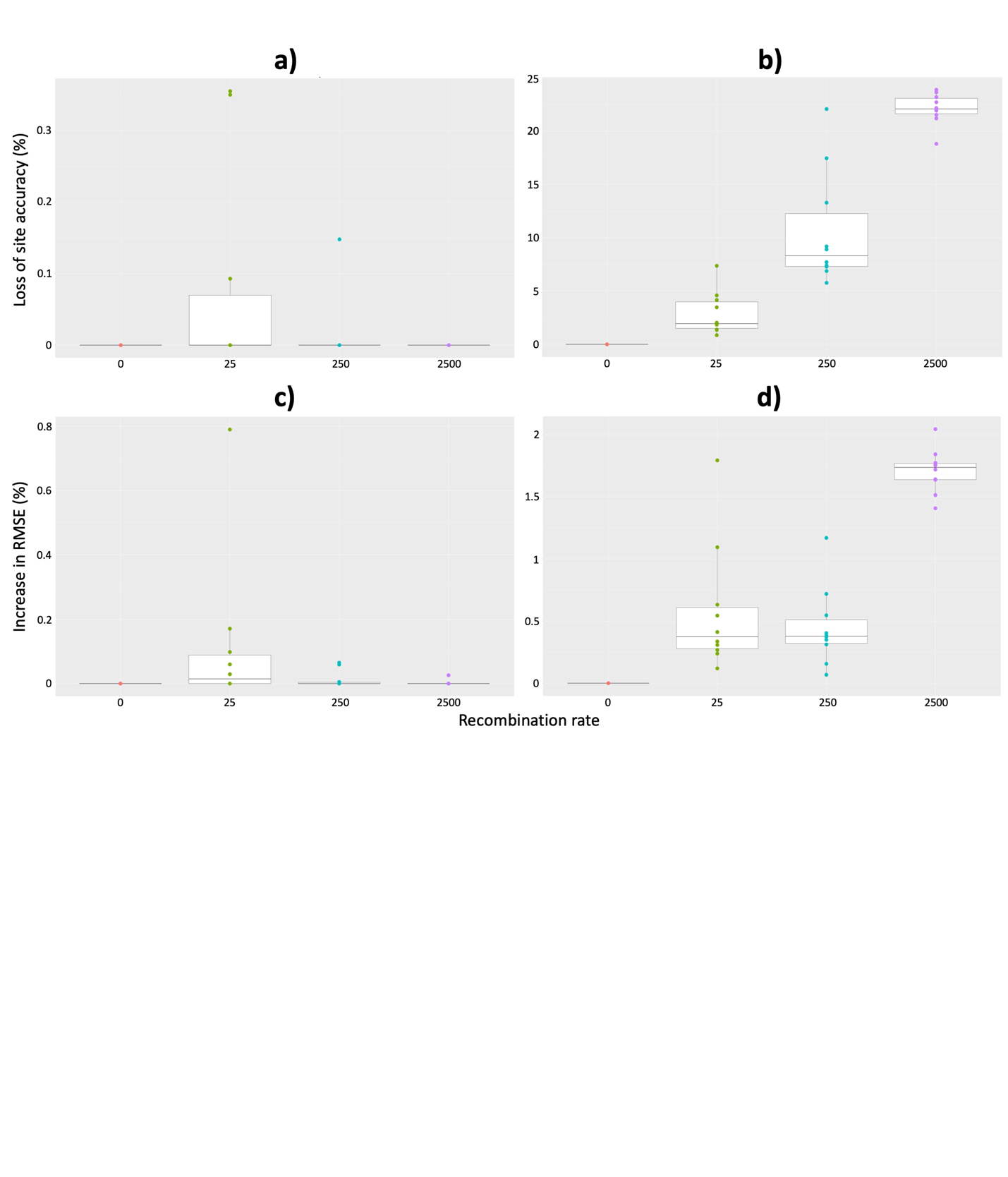

**Figure S20.** (a-b) Site accuracy loss and (c-d) RMSE gain of AIC (left) and BIC (right) across recombination rates on simulated chromosomes with high ILS level. Each dot represents an individual result (i.e., ten replicates per recombination rate).

­­­
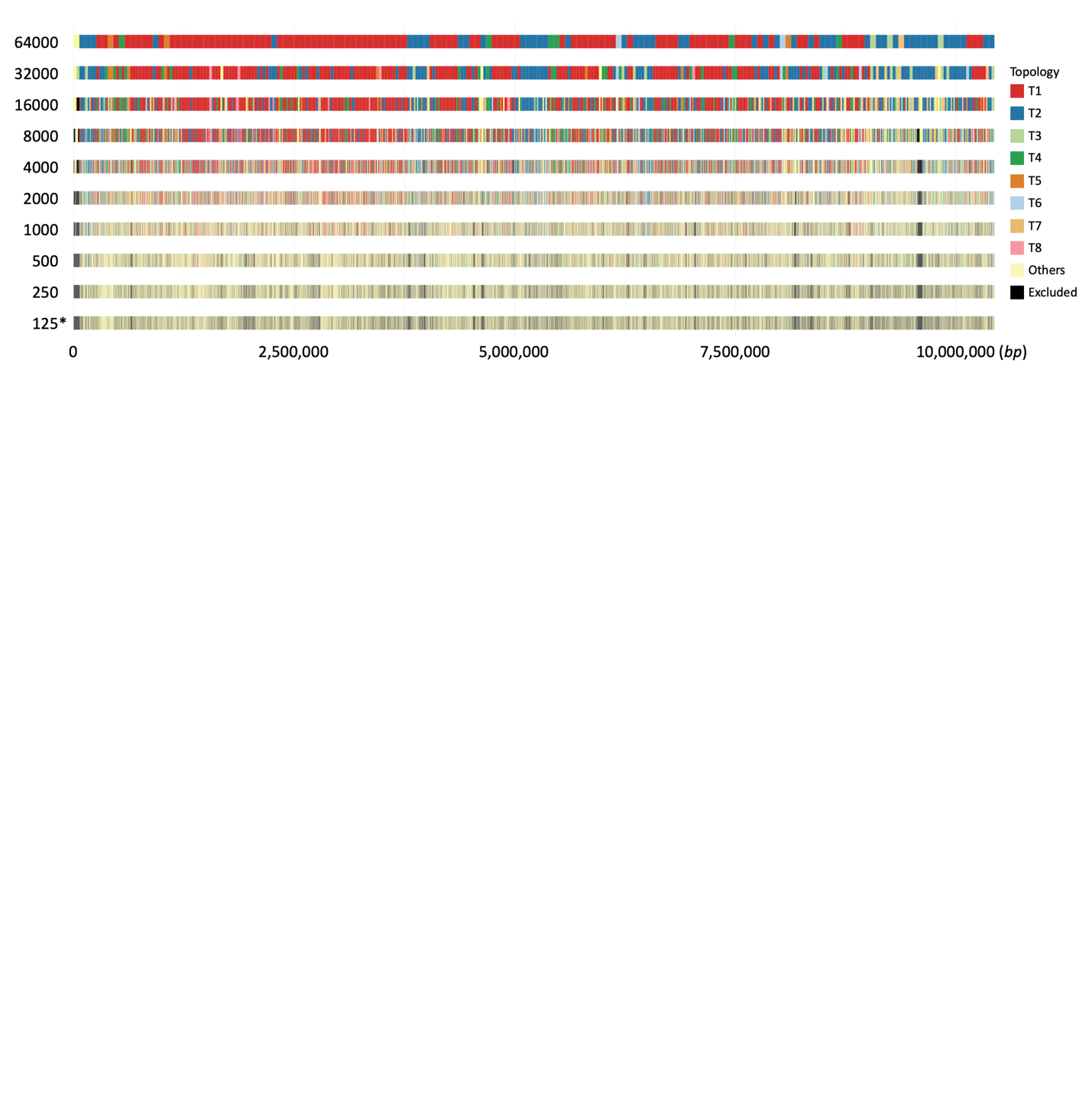

**Figure S21.** Topology distribution ­­­­­­of chromosome 11 of *erato-sara* *Heliconius* butterflies across window sizes. Asterisk (*) shows the best window size from stepwise non-overlapping windows (Table S4). Colouring is based on tree topology from Edelman et al. (2019), with *Others* for other topologies and *Excluded* for win­­dows that are excluded from the analyses. When one window size was compared to several other window sizes (e.g., 32kb windows were compared with 64kb windows and 16kb windows), we took the distribution that includes more windows.

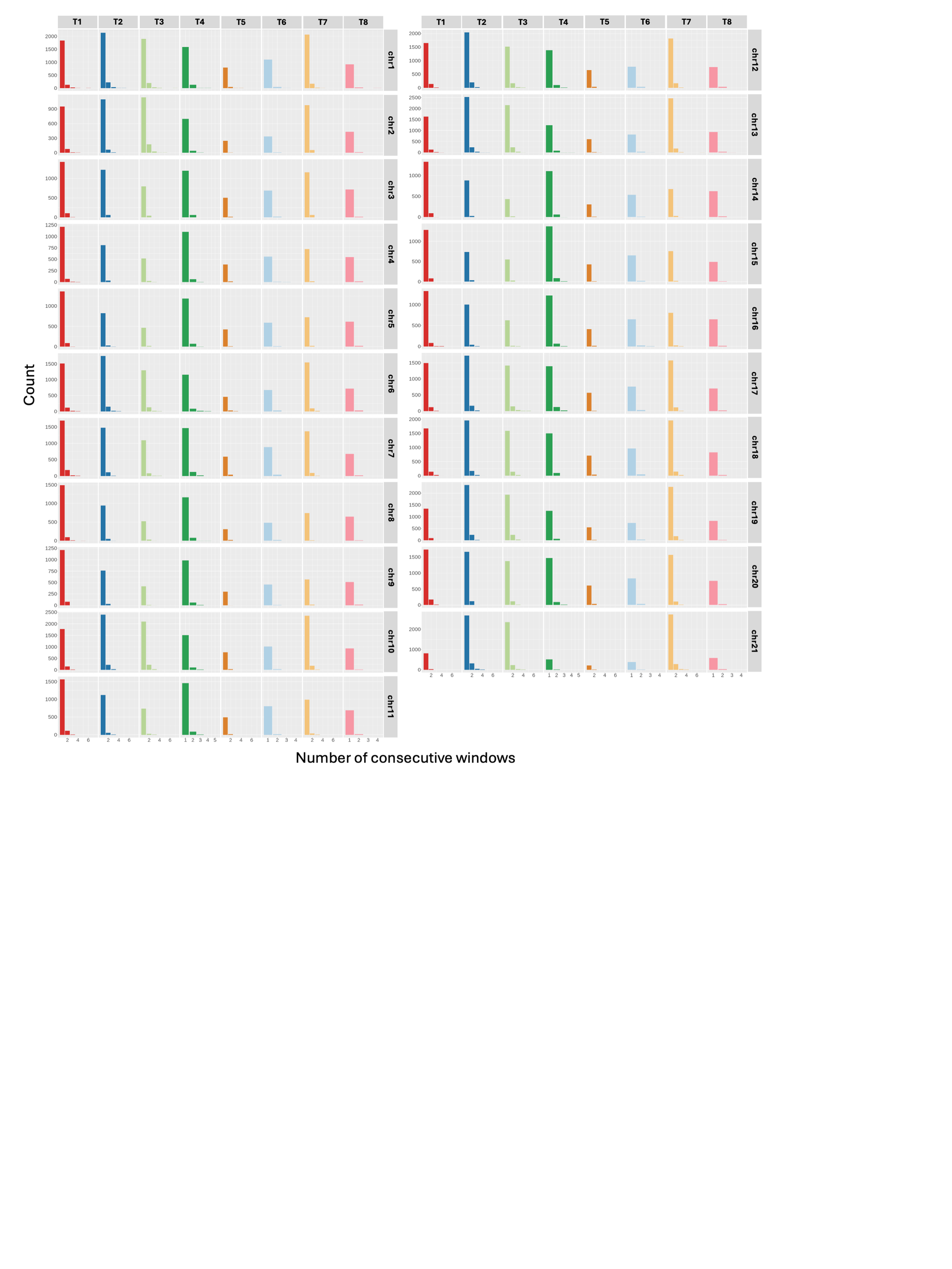

**Figure 22.** Per-chromosome count of consecutive windows that recover the same topology from stepwise non-overlapping windows analysis on *erato-sara Heliconius* butterflies’ genome based on the best window size for each chromosome (Table S4). Colouring is based on tree topology from Edelman et al. (2019).

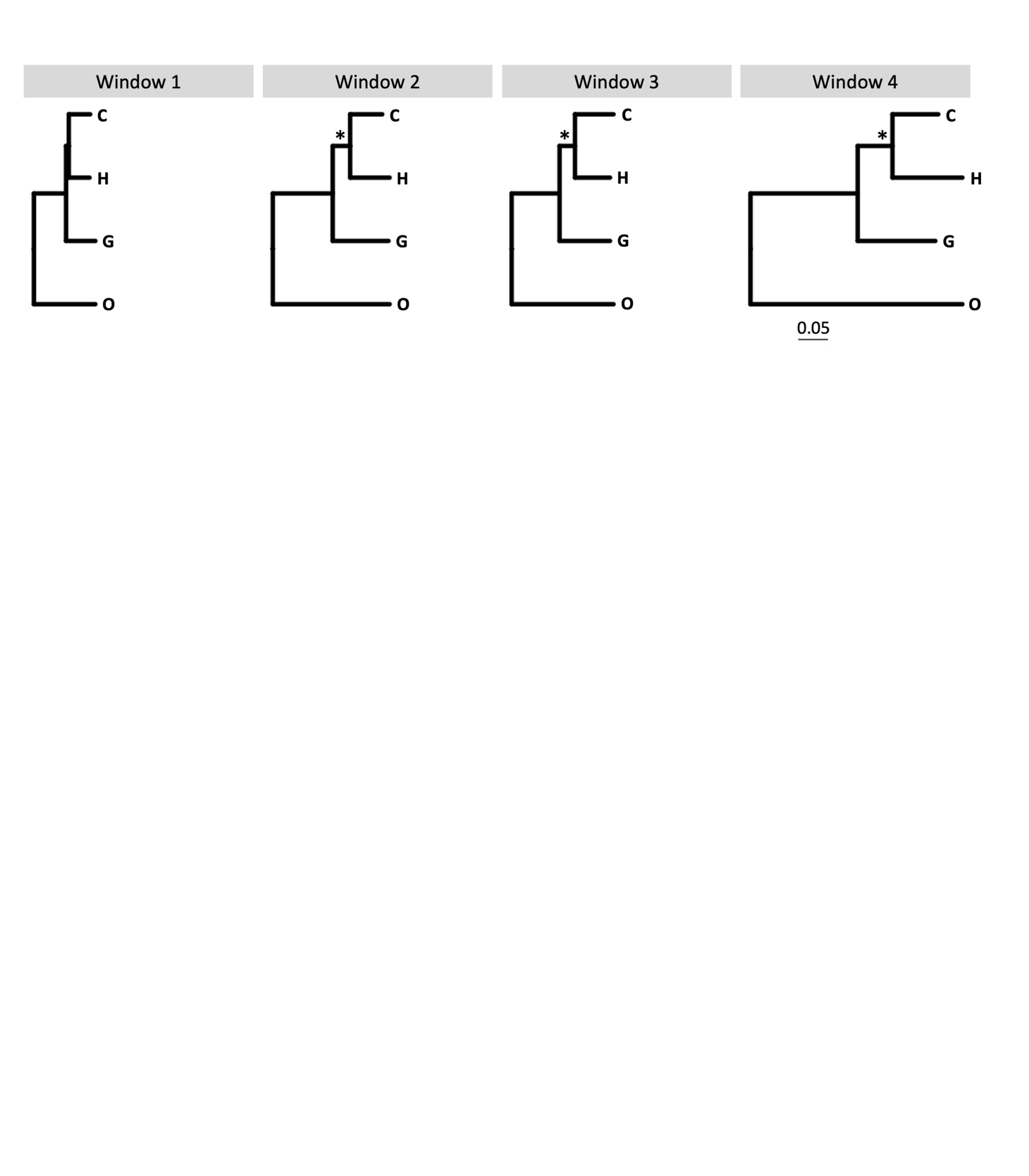

**Figure 23.** 4kb window trees for mitochondrial DNA of great apes. H: human, O: orangutan, C: chimpanzee, G: gorilla. Asterisk (*) denotes node with ≥95 UFBoot support.

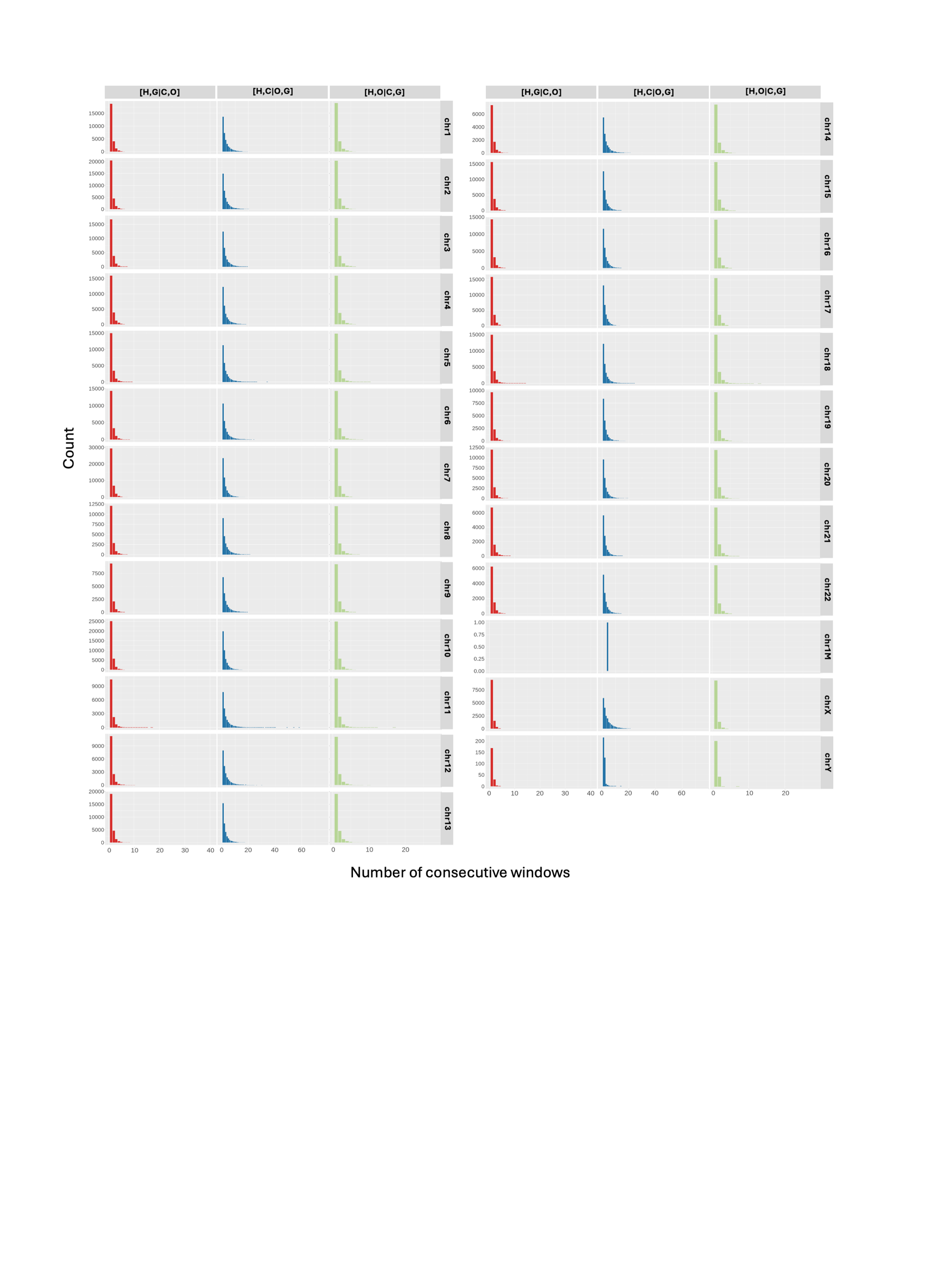

**Figure 24.** Per-chromosome count of consecutive windows that recover the same topology from stepwise non-overlapping windows analysis of great apes’ genome based on the best window size for each chromosome (Table S8). H: human, O: orangutan, C: chimpanzee, G: gorilla. Colouring is based on tree topology.

­­
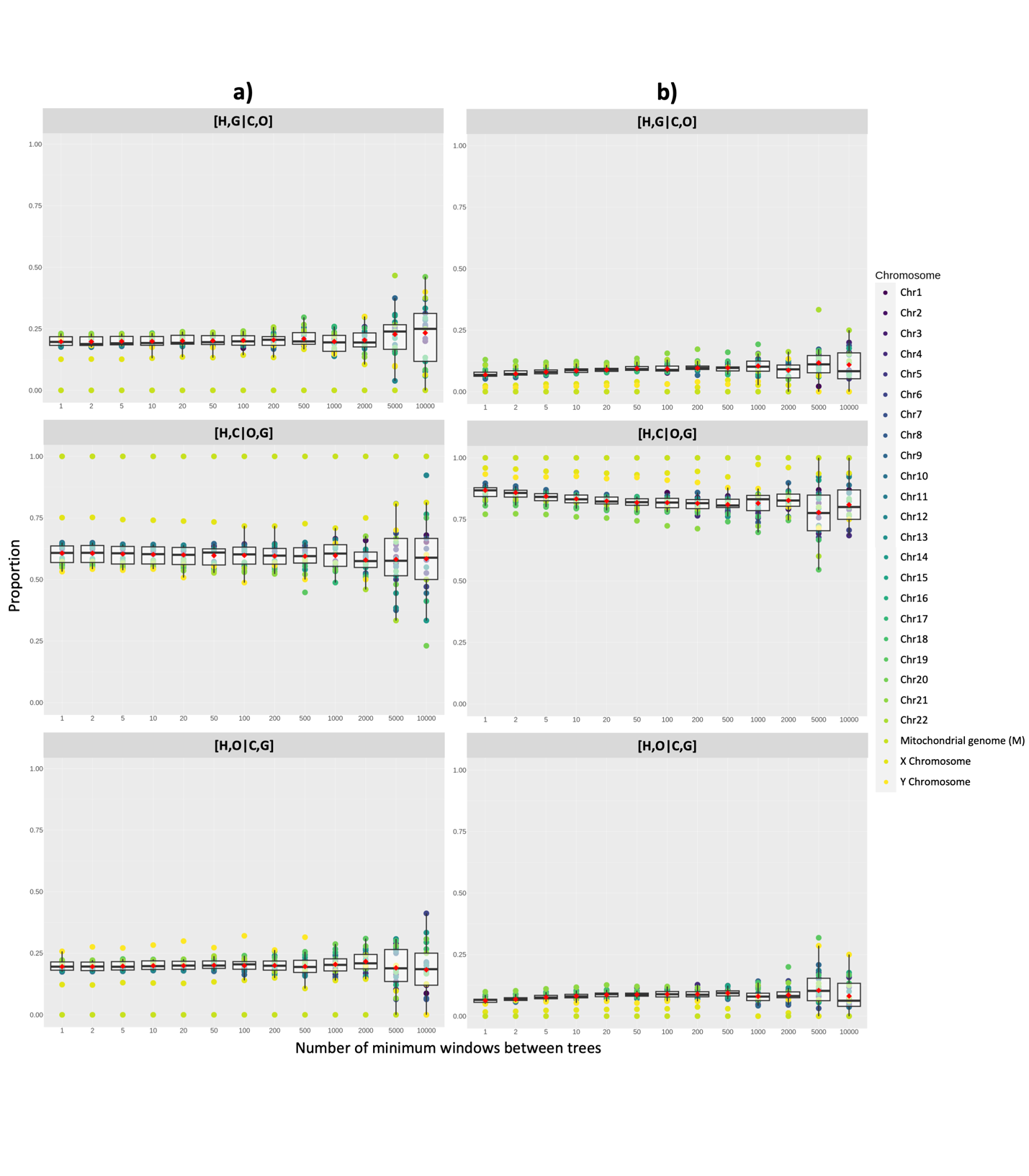

**Figure S25.** Proportion of tree topologies based on (a) all window trees and (b) ≥95 average UFBoot trees between human [H], orangutan [O], chimpanzee [C], and gorilla [G] across different minimum window intervals based on the best window size for each chromosome (Table S8). Red dots show the average proportion per window interval across chromosomes.

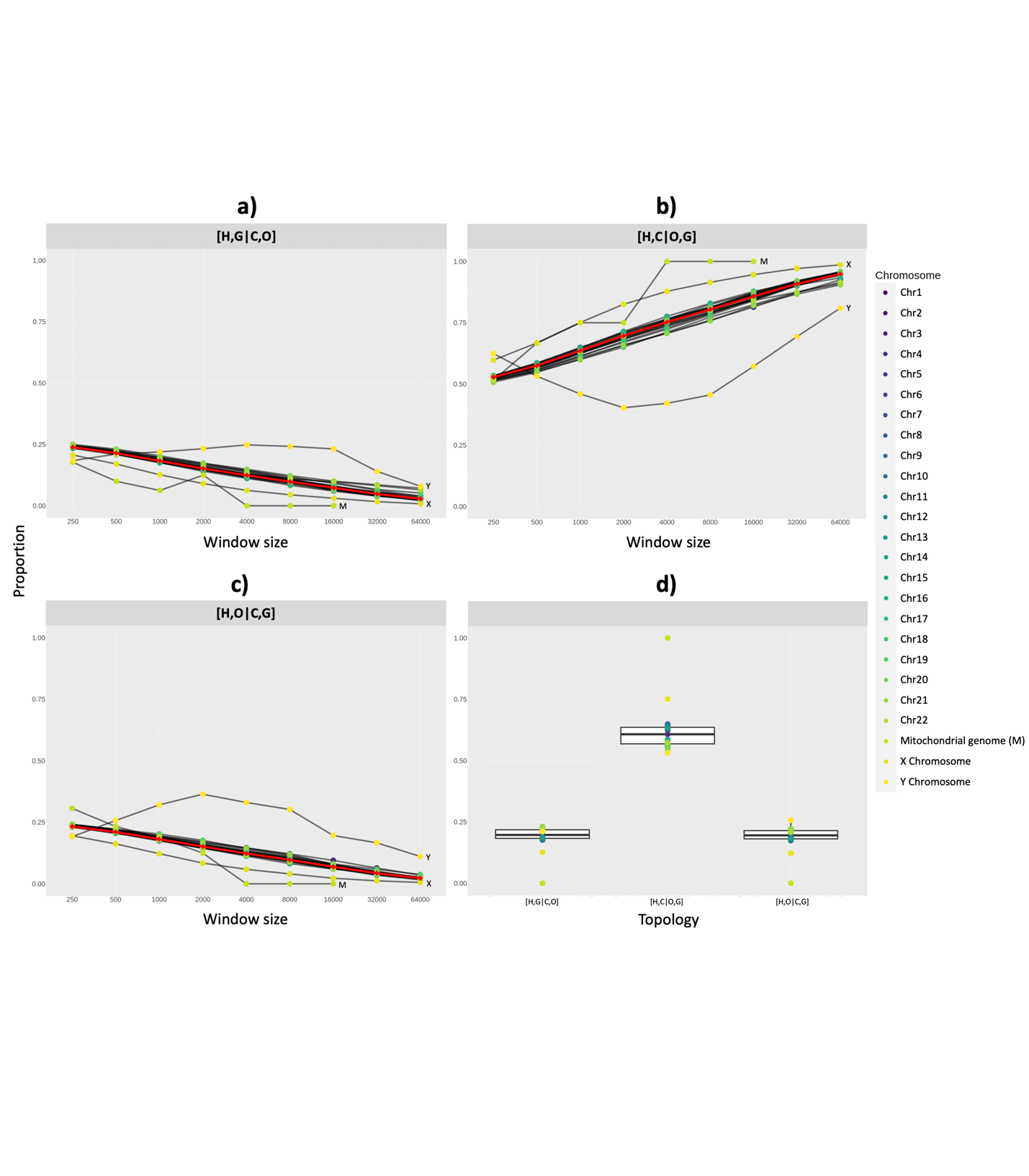

**Figure S26.** Proportion of all window tree topologies between human [H], orangutan [O], chimpanzee [C], and gorilla [G] across (a-c) constant window sizes and (d) the best window size for each chromosome (Table S8). Black lines connect the same chromosome across window sizes. Red lines show the average proportion per window size across chromosomes.

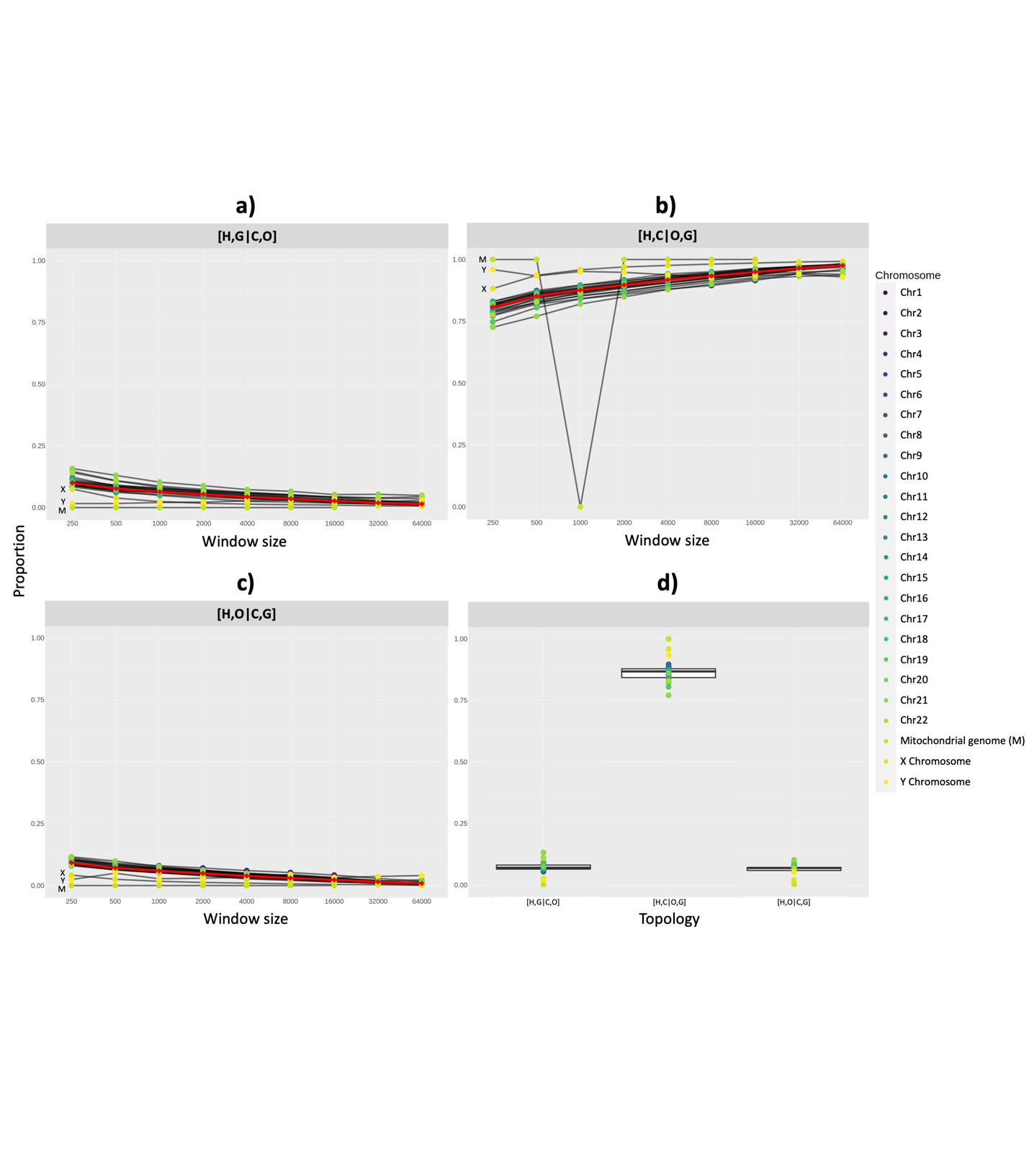

**Figure S27.** Proportion of window tree topologies with ≥95 average UFBoot values between human [H], orangutan [O], chimpanzee [C], and gorilla [G] across (a-c) constant window sizes and (d) the best window size for each chromosome (Table S8). Black lines connect the same chromosome across window sizes. Red lines show the average proportion per window size across chromosomes.
